# Detection of eco-evolutionary dynamics in communities using Joint Species Distribution Models

**DOI:** 10.1101/2024.11.30.626152

**Authors:** Jelena H. Pantel, Ruben J. Hermann

## Abstract

Biodiversity at the metacommunity scale is typically influenced by a number of environmental, spatial, biotic, and stochastic factors. At the same time, these factors impact the evolution of individual species, as sites present different local selection pressures and connectivity can impact gene flow and genetic drift. Identifying the relative impacts of environmental, spatial, biotic, and other drivers on community composition across spatial and temporal scales has been greatly facilitated by joint species distribution models, but these models have yet to consider the impact of microevolution on community composition. We used Heirarchical Models of Species Communities (HMSC) to analyze simulated data of an populations and communities, including in a large evolving metacommunity model, to establish whether HMSC can sufficiently quantify the contribution of phenotypic evolution for metacommunity composition. The models successfully partitioned variance contributed by environmental, spatial, and evolving phenotypic drivers, and also estimated site-and year-specific covariance. We also applied the HMSC with trait evolution model to an existing dataset studying trait change and community dynamics in an experimental aquatic plant system. The study of eco-evolutionary dynamics may require data that reflects numerous complex, interacting processes and it is necessary to have flexible, generalized statistical models to analyze this data. JSDM models such as HMSC present one promising path for analysis of eco-evolutionary dynamics in multi-species communities.

## 3 Introduction

Analysis of biodiversity at large spatial scales can be complex, because numerous environmental properties, spatial structure and connectivity, interactions between species, as well as stochastic processes such as drift and priority effects operate and determine observed patterns. Statistical models in ecology have made substantial progress in reflecting these numerous ecological processes at multiple scales. There are now a diversity of joint species distribution models [JSDMs; Pollock et al. (2014)] (Pichler and Hartig 2021) and multi-species occupancy models (Devarajan et al. 2020) (Poggiato et al. 2021), which can incorporate these diverse components that structure variance in biological data. Many of these models have implementations in R packages (e.g. spOccupancy, (Doser et al. 2022)), and are used to measure the relative importance of different ecological drivers in e.g. communities of fungi, birds, mammals, phytoplankton, fish, and other taxa, from local to continental scales (e.g. (Abrego et al. 2020) (Antão et al. 2022) (Keppeler et al. 2022)).

Despite the variety of components implemented in these models, there are some additional drivers of community composition which are not currently reflected. The dynamics of populations and communities, and resulting biodiversity patterns, are also influenced by the genotypic and phenotypic properties of organisms. Although traits can be considered in some JSDMs, they typically require fixed values for species’ traits (e.g. (Ovaskainen et al. 2017) (Tikhonov et al. 2020); but see Abrego et al. (2024)). However, ecological dynamics can impact selection on genes and phenotypes, potentially resulting in interactions or feedback loops between ecological and evolutionary processes (Lion 2018) (Barbour et al. 2022). Although an increasing number of theoretical and empirical studies have evaluated the impact of evolutionary dynamics for community assembly and metacommunity dynamics (Loeuille and Leibold 2008) (Osmond and Mazancourt 2013) (Pantel et al. 2015) (Toju et al. 2017), analytical methods to consider this additional complexity lag behind the scope of data collected in eco-evolutionary studies. Some methods exist that ultimately apply ANOVA to partition variance in traits or growth rates across contributions of ecological and evolutionary components ((Hairston Jr et al. 2005) (Govaert et al. 2016); methods for discrete and continuous analyses are described in Ellner et al. (2011)). These methods are more frequently cited for their potential rather than for their actual application to empirical data (but see (terHorst et al. 2014) (Hiltunen and Becks 2014)), because this approach is ultimately limited by the same requirements that can limit application of ANOVA - data should follow a normal distribution, the processes that structure the target response variable must be linear and additive, and complicated structuring mechanisms must be reduced into one or a few target categories labeled as ‘ecology’ or ‘evolution’. Flexible models, which can better reflect the mechanisms and structure of the diverse kinds of data that most studies of eco-evolutionary dynamics will collect, are needed for rigorous inference about eco-evolutionary processes (Pantel and Becks 2023).

Two existing statistical models are potentially well suited for analysis of eco-evolutionary data at the microevolutionary and metacommunity scale but also have some considerations before their application to complex community data. Lasky et al. (2020) (Lasky et al. 2020) developed an integrated reaction norm model, linking genetic, phenotypic, and demographic processes, and Benito Garzón et al. (2019) (Benito Garzón et al. 2019) presented a species distribution model with local adaptation and phenotypic plasticity (Δ SDMs). The model of Lasky et al. is an excellent candidate for spatially complex population dynamics, and awaits an extension to the multispecies level. The Δ SDM models described by Benito Garzón et al. truly disentangle the role that plastic and evolutionary trait divergence can play in species distributions, but they also require common garden experimental data across sites to estimate these effects. Observational data collected in metacommunity surveys may only have survey data across sites or time points, as large-scale experimental measures of reaction norms required by the Δ SDM model may be difficult to obtain.

One particular metacommunity statistical model, the Hierarchical Modelling of Species Communities (HMSC) framework (Ovaskainen et al. 2017) (Tikhonov et al. 2020), explicitly considers a multitude of ecological processes that can structure community composition across time and space (Leibold et al. 2022) and is potentially well suited to consider analysis of metacommunity eco-evolutionary processes. This model infers the role of environmental filtering via variation and covariation in how species respond to their environment, considers the impacts of (fixed) trait values and phylogenetic relationships, and uses latent variables to account for the numerous unobserved environmental and spatial features that are difficult to exhaustively measure in community surveys. Previous studies have demonstrated that this statistical model can reflect the diverse array of community assembly and metacommunity processes that structure species abundances (i.e. species sorting, environmental filtering, priority effects; (Little and Altermatt 2018) (Ovaskainen and Abrego 2020) (Leibold et al. 2022)). The associated R package has been applied to study environmental and abiotic drivers of diverse assemblages of organisms (e.g. (Sandal et al. 2022) (Weigel et al. 2023)). HMSC is potentially useful for studying eco-evolutionary drivers of community structure, but the model has not previously been applied to include heritable changes in trait values over time.

In this study, we use simulations of population dynamics, environmental drivers, species interactions, and trait evolution in populations, communities, at the local and metacommunity scale, to study eco-evolutionary dynamics. After generating simulated time series of population sizes and mean trait values, we then use HMSC to analyze the resulting data, to determine whether HMSC can successfully estimate the impacts of trait evolution for community composition, and to generate relative contributions of fixed environmental and trait evolution drivers as well as random effects of spatial and temporal covariance among species. We asked three main questions: (i) can linear statistical models (like HMSC) accurately capture the impact of non-linear population, community, and evolutionary dynamic processes?, (ii) can HMSC be used to evaluate hypothesized drivers of the relative importance of evolution for community composition at the metacommunity scale?, and (iii) can we better understand the role of evolution for community dynamics in experimental systems using HMSC?

## 4 Methods

### 4.1 Linear statistical models for one-and two-species models of growth, competition, and trait evolution

We evaluated whether linear regression models can appropriately capture the dynamics that drive species abundances in relatively simple populations and communities. We fit data generated from simulation models with increasing complexity (Table 1) to Bayesian linear regression models. These simulation models were (A) a single-species model of discrete-time logistic population growth, (B) expanded to include a randomly fluctuating environmental property *E* that impacts the species’ population growth rate, (C) a two-species model of logistic growth and competition that was then (D) expanded to consider a randomly fluctuating environmental property, (E) and further expanded to consider evolution in a trait *x* that is selected by the environmental property, and finally (F) the model of two-species growth, competition, environmental change, and evolution in a spatially structured (10-patch) habitat.

**Table 1:**
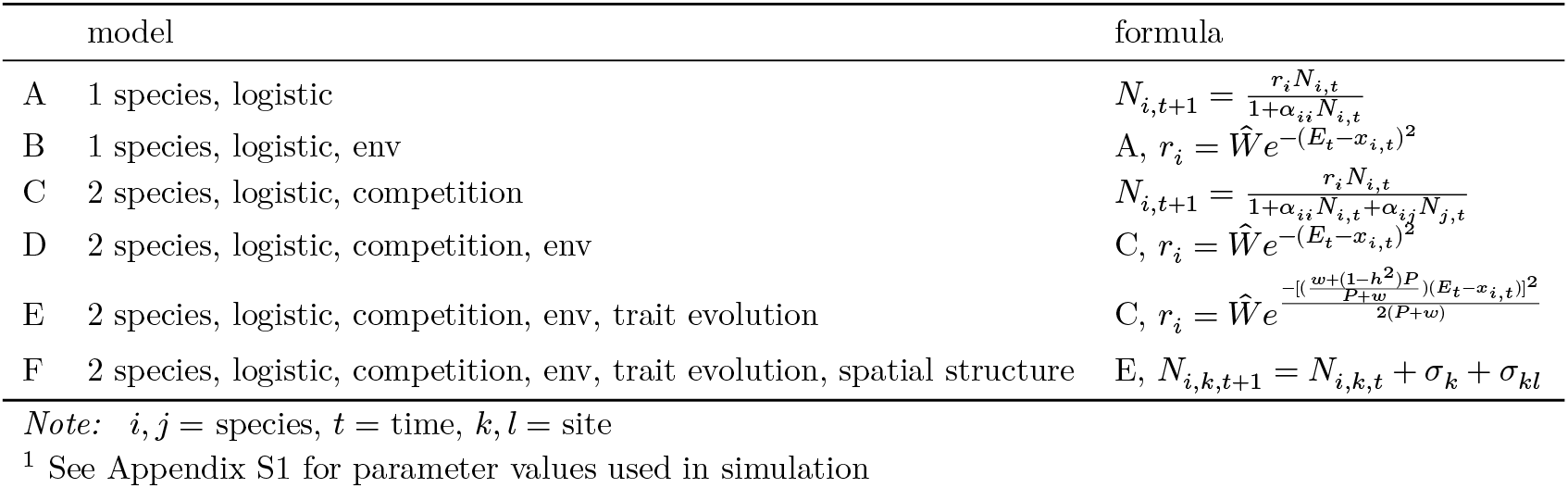
Models for population growth.

To fit linear regression models to the simulated population size data, we began with models using population size (model *A*) and environment (including the second-order term, to better reflect species optimum values; model *B*) as fixed effects 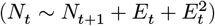, and progressed to a mixed model using environment and trait evolution as fixed effects 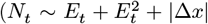 for simulation model *E*) and time (models *C, D, E*, and *F*) and spatial distance (model *F*) as random effects. For consistency, all linear regression models were implemented as hierarchical Bayesian models using HMSC (Hierarchical Modelling of Species Communities (Ovaskainen et al. 2017) (Tikhonov et al. 2020)). HMSC uses a latent variable approach as an alternative to directly modelling species interaction coefficients, estimating a reduced number of linear combinations of species abundances that best predict future population size values. For linear models fit to data generated in simulations *C, D, E*, and *F* above, the time random effect was implemented to capture species interactions (as part of the temporal species association matrix estimated by the model), and the spatial random effect was implemented in model *F* to capture spatial covariance in abundances as a function of distance between sites. Full methodological details are given in Appendix S1. The one-two species simulations were run in R (R Core Team 2024) using the ecoevoR package (Pantel and Becks 2023) and HMSC models were implemented using the Hmsc package (Tikhonov et al. 2022).

### 4.2 Evolution in metacommunities and HMSC

#### 4.2.1 Simulation model

We also simulated growth and competition dynamics for a multi-species assemblage in a patchy landscape, with site variation in one environmental property, and with evolution in a trait that is selected on by the environment. Population growth for species in the metacommunity simulation followed a Leslie-Gower model (a discrete-time version of a Lotka-Volterra model (Beverton and Holt 1957) (Leslie and Gower 1958)). We considered the impact of trait evolution for growth using a discrete time quantitative genetic model of evolutionary rescue (Gomulkiewicz and Holt 1995). The model for population size was:

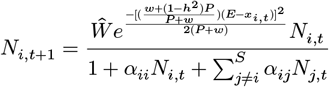

where *N*_*i*_, *t* is the population size of species *i* at time *t, Ŵ* is calculated as 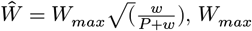 is the species’ maximum per-capita growth rate, *w* is the width of the Gaussian fitness function (which determines the strength of selection), *P* is the width of the distribution of the phenotype *x, h*^2^ is the heritability of the trait *x, E* is the local environmental optimum trait value (*x*_*opt*_), *x*_*i,t*_ is the trait value of species *i* at time *t, α*_*ii*_ is the intraspecific competition coefficient (the per capita impact of species *i* on itself) and *α*_*ij*_ is the interspecific competition coefficient. Populations have a critical density *N*_*c*_, below which the population is subject to extinction due to demographic stochasticity at a probability of *p* (Gomulkiewicz and Holt 1995).

This model was used in Pantel & Becks (Pantel and Becks 2023) to evaluate the consequences of adaptive evolution for coexistence in a three-species system. To expand this model to a metacommunity, we consider the evolution of multiple species (*S* = 15) in a landscape of patches (*k* = 50). The patches have values of an environmental property *E ∼ U*(0, 1) that determines the local optimum phenotype *E* = *x*_*opt*_, i.e. where species experience their maximum absolute fitness *W*_*max*_, and patches also have spatial locations *X* and *Y* (both drawn from *U*(0, 1)). The patches thus have a connectivity matrix **D** (here given by their Euclidean distance), as well as a connectivity matrix **C** that is a Gaussian function of **D** and species dispersal rate *d*_*i*_.

We used the evolving metacommunity model to test whether landscape connectivity (the overall dispersal rate *d*) and the speed of evolution (varying *h*^2^) influenced the relative importance of trait evolution (|Δ*x*|) for metacommunity composition *N*_*i,t,k*_ (evaluated as *V*_|Δ*x*|_, the proportion of variation in species abundances explained by |Δ*x*| as estimated by the HMSC model fit to the simulated data across *d* and *h*^2^ levels). We hypothesized that the overall importance of trait evolution (*V*_|Δ*x*|_) would decrease with increasing dispersal rate - at low dispersal rates, locally mal-adapted species are less likely to emigrate to more optimal patches and will experience strong local selection, leading to a large amount of trait evolution. Population sizes for initial, mal-adapted populations will also likely be lower, leading to reduced relative importance of density-dependent intra-and interspecific competition. We also hypothesized that heritability (*h*^2^) would mediate the relative importance of dispersal rate - at higher *h*^2^, species will evolve quickly and we expected the importance of trait evolution *V*_|Δ*x*|_ to depend on the amount of time that mal-adapted populations persist. Therefore, simulations with higher *h*^2^ would have decreased importance of evolution *V*_|Δ*x*|_ for community composition. For the same reason, we hypothesized generally that, for a given dispersal rate *d*, the importance of evolution *V*_|Δ*x*|_ would decrease with increasing *h*^2^.

All simulations used the same parameter values (*W*_*max*_ = 2, *P* = 1, *w* = 2, *N*_*c*_ = 100, *p* = 0.001), species interaction matrix (*α*_*ii*_ = 0.00125; *α*_*ij*_ *∼ U*(0, 0.0015)), and initial conditions (initial richness *s*_0*k*_ *∼ Pois*(0.75); initial population size *N*_0*i*_ *∼ P ois*(10); initial degree of maladaptation *β*_*i*0_ *∼ Gamma*(0.75, 1), initial distance to the local patch’s optimum phenotype 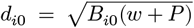, initial trait value *x*_*i*0_ = *E*_*k*_ *− d*_*i*0_; Appendix S4). Simulations were run across a range of dispersal values (identical for all species; *d* = 0, 10^*−*9^, 10^*−*8^, 10^*−*7^, 10^*−*6^, 10^*−*5^, 10^*−*4^, 10^*−*3^, 10^*−*2^, 0.1, 0.2, 0.3, 0.4, 0.5, 0.6, 0.7, 0.8, 0.9, 1) and heritability values (identical for all species: *h*^2^ = 0, 0.1, 0.2, 0.3, 0.4, 0.5, 0.6, 0.7, 0.8, 0.9, 1) for 200 time steps, producing records of *N*_*ikt*_ and *x*_*ikt*_. Code for all simulations and model fits is available at https://github.com/jhpantel/ecoevo-hmsc/tree/aRxiv.

#### 4.2.2 Statistical model

To determine the relative influence of environmental and spatial properties, intra-and interspecific population dynamics, and evolutionary dynamics for community structure, we applied HMSC to model the log-abundance at each site and year for each species. The log-abundance values were modeled as *In*(*N*_*ikt*_) *∼ N*(*L*_*ikt*_, *σ*_2_), where the linear predictors **L** are the sum of fixed and random effects **L**^**F**^ + **L**^**R**^ (Tikhonov et al. 2020). The fixed effects were 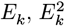, and the absolute value of the change in each species’ trait value from the previous time step to the next |Δ*x*_*ik,t−*1*→t*_|. Random effects included time (the day of the sample) and space. The spatial latent variables represent a spatial model, where species respond to some latent spatial predictor associated with the *xy* coordinates of the sites (Tikhonov et al. 2020). When there was no dispersal in the system (*d* = 0), spatial random effects were not included in the model.

#### 4.2.3 Applying HMSC to empirical data

To test the applicability for HMSC on ecological and evolutionary effects in empirical data, we applied HMSC to the experimental data of Jewel & Bell (2023) (Jewell and Bell 2023). The authors conducted a community selection experiment in which they grew four floating and short-lived aquatic plants (the angiosperms *Lemna minor* (Lm), *Spirodela polyrhiza* (Sp) and *Wolffia columbiana* (Wc), and the liverwort *Ricciocarpos natans* (Rn)) under different environmental treatments (shade cover and nutrient levels) for twelve weeks. The shade cover and nutrient levels for the experimental pools were a factorial combination of 3%, 12% and 50% shade, and low (dissolved nitrogen (DN) = 200 *μgL*^*−*1^, dissolved phosphorous (DP) = 10 *μgL*^*−*1^), medium (DN = 800 *μgL*^*−*1^, DP = 40 *μgL*^*−*1^), and high (DN = 3200 *μgL*^*−*1^, DP = 160 *μgL*^*−*1^) concentrations of nutrients, respectively. The frond size of ten individuals from each species and the density of each species were measured weekly for the eleven weeks of the experiment. To estimate the importance of trait change in the HMSC model, we calculated the average trait change as 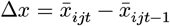, where 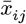 described the mean trait of species *i* in environment *j* and week *t*.

To determine the relative importance of trait change for community composition, we fit two HMSC models to the data: An ‘evolution model’ in which |Δ*x*| of each species and environmental treatment were used as explanatory variables, and a ‘no-evolution model’, in which only the environmental conditions were included. Species density was log-transformed species, and we included replicates (*n* = 2) and time as random effects. For each model we ran four MCMC chains to generate 1000 samples of the posterior distribution for all model parameters. To evaluate the model-estimated role of trait change for predicted population size, we used the posterior distributions of the estimated effect size of |Δ*x*| multiplied by the observed 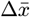 of each species, added to the model-estimated intercept.

## 5 Results

### 5.1 Linear statistical models for one-and two-species models of growth, competition, and trait evolution

Researchers often use linear regression models to study non-linear population and community dynamics. We evaluated the performance of linear models in simple cases (1-2 species, 1-few sites), to validate that effect size estimates and predicted densities accurately reflected mechanisms before moving onto more complex models in the next sections of our analysis. When only one species was present (models *A* and *B*), the regression coefficients (model 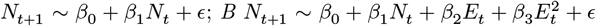) reflected the direction and magnitude of the mechanistic drivers of population size. *β*_1_ correctly captured the positive relationship between *N*_*t*_ and *N*_*t*+1_ under our simulation conditions, and *β*_2_ and *β*_3_ correctly captured the quadratic unimodal relationship between *E*_*t*_ and *N*_*t*_ + 1 (Appendix S1.1, S1.2).

When a second species was added (models *C* and *D*), the regression coefficients performed well for predicting density values at all time points, but the direction and magnitude of the regression coefficients did not reflect the intra-and interspecific competition strengths included in the mechanistic model (Appendix S1.3). We instead used a latent variable modelling approach (implemented in HMSC) to consider these species interactions as part of the temporal random effects included in the model. This approached produced effect size estimates that both predicted the population dynamics well (Figure 1a) and captured the conditional effects of environment for species densities (Figure 1b). The variation partition also successfully captured the relative importance of the environmental predictor and the temporal random effects for model *D* (which in this simple case were driven by intra-and interspecific species interactions; Figure 1c; Appendix S1.3, S1.4).

**Figure 1.**
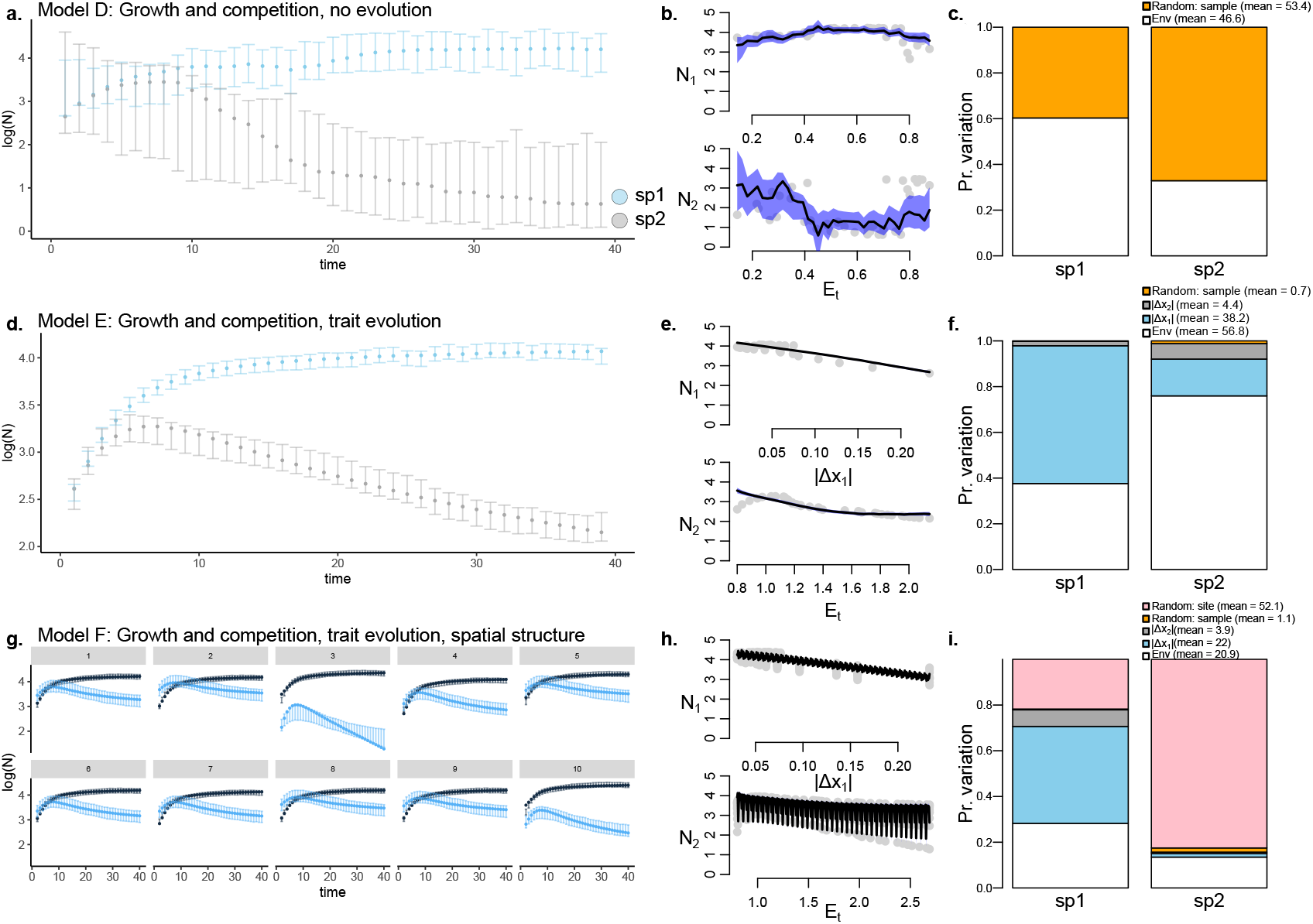
Simulation models for 2-species ecological and eco-evolutionary dynamics. Panels a, d, and g show log-population size (y-axis) over time (x-axis) for species 1 and 2 when they grow and compete for resources with random environmental fluctuations over time. Error bars give the 95% highest probability density interval (HPDI) for posterior predictions from the HMSC model fit. Species either have fixed (panel a) or evolving trait values (panels d, g) and are present in either 1 (panels a, d; simulation models D and E) or multiple sites (panel g; model F). Graphs in panels b, e, and h show observed (points) and model-predicted (black line with blue shading) log-population size (*Iog*(*N*)) for species 1 and 2 across gradients of the environment (*E* + *E*^2^) and the change in trait values of species 1 (|Δ*x*_1_|). Graphs in panels c, f, and i show the results of a variation partition after fitting data to the HMSC statistical model. The variation partition plot shows the proportion of variation explained by environment, trait evolution (|Δ*x*_1_| and |Δ*x*_2_|), and temporal and spatial random effects for *In*(*N*) for species 1 and 2. Error bars and shading in panels a, d, and g represent the 95% highest probability density interval for posterior predictions.

Trait evolution was included as a predictor when it was introduced in model *E*. The resulting linear regression coefficients could predict species dynamics (Figure 1d) and reflected the impact and magnitude of environmental and trait evolution predictors (Figure 1e). The variation partition also showed that trait evolution in species 1 (|Δ*x*_1_|) was a strong driver of variation in abundance (blue portion of bar plot, Figure 1f), primarily for species 1 but also for species 2 as well (Appendix S1.5; Appendix S2).

After adding spatial complexity to the 2-species simulation, we included a spatial random effect to capture species covariances as a function of distance between sites (model *F*). By properly considering species interactions as a temporal random effect and including spatial random effects, the HMSC model could estimate regression coefficients for the impacts of the environment *E*_*t*_ and trait evolution |Δ*x*_1_| and |Δ*x*_2_| for both species that fit well to the simulated data points (Figure 1g); Appendix S1.6) and capture conditional effects of environment and trait evolution for both species (Figure 1h). Spatial random effects explained a large proportion of variation in abundance for both species, but trait evolution remained a strong driver for species 1.

### 5.2 Evolution in metacommunities and HMSC

The simulation model followed population size and trait value dynamics for species in the landscape across 200 time steps (Figure 2a). Resulting biodiversity dynamics depended on the dispersal level and heritability values (Figure 2b). For dispersal, results mirrored some expectations from existing metacommunity models, i.e. that regional *γ* diversity decreases with increasing dispersal level *d* ((Mouquet and Loreau 2003); Figure 2b). The inclusion of trait heritability had a strong effect on community diversity, reflecting that local adaptation can rescue species and contribute to richness at the local and metacommunity scale. Regional species diversity (*γ*) increased with increasing *h*^2^ levels (Figure 2b).

**Figure 2.**
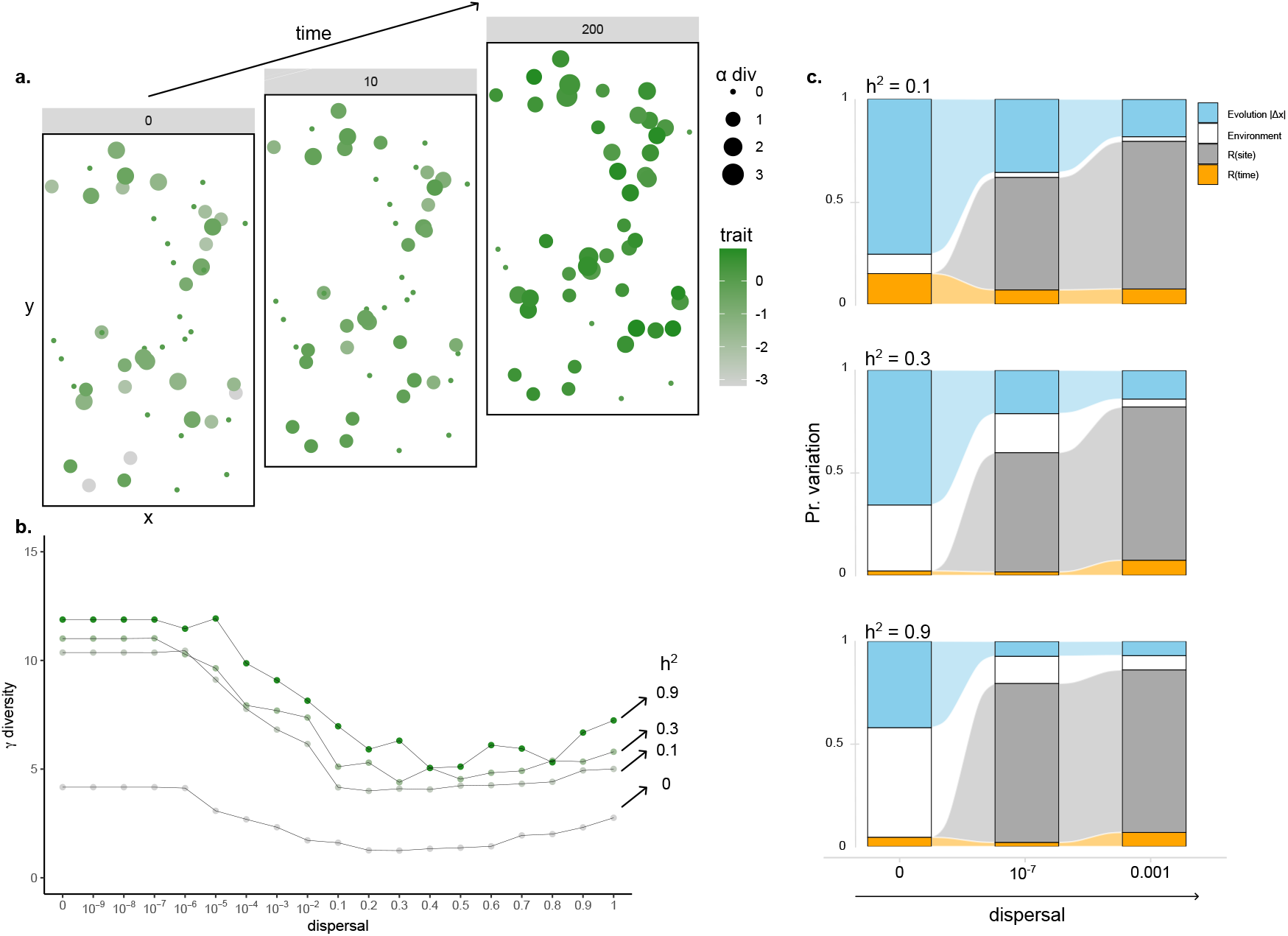
Importance of trait evolution for change in community composition at the metacommunity scale. (a) We developed a simulation model for population growth, competition, and trait evolution in 15 species that inhabit a patchy landscape. The landscape has 50 sites (colored points in the XY space), with distinct local environments that select for different patterns of local species diversity (size of points) and average trait values (color of points; results are shown at time *t* = 0, 10, 200 for a simulation with *d* = 10^*−*3^ and *h*^2^ = 0.1). (b) Regional metacommunity *γ* diversity decreased with increasing disperal level and increased with increasing heritability (*h*^2^). (c) Variation partition plots after fitting metacommunity time series of species abundances *Iog*(*N*_*ikt*_) to HMSC models with environment, trait evolution, and random effects (space and time) as predictors. These variation partition results are averaged across all species in the metacommunity. The top panel is for *h*^2^ = 0.1, the middle panel is for *h*^2^ = 0.3, and the bottom panel is for *h*^2^ = 0.9. The alluvial plots show the change in relative importance of different predictors for increasing dispersal levels.

Analysis using the HMSC model for community time series across *d* and *h*^2^ levels indicated that the importance of evolution for community composition (*V*_|Δ*x*|_) decreased with increasing dispersal and with increasing heritability (Figure 2c). The importance of spatial random effects increased with dispersal rate, and overall the importance of evolution decreased with increasing *h*^2^. These effects can be better understood by our supplemental analyses of 1-species, 1-site simulation models of evolutionary rescue (Appendix S3).

#### 5.2.1 Empirical data

We fit the empirical data (community dynamics and frond size of four aquatic plant species under different environmental treatments) to an HMSC model, and estimated the effect size of shade cover, nutrient level, and trait change (in all four species) for changes in species densities. The variation partition produced by the HMSC model fit shows the estimated relative importance of evolutionary (change in the frond size trait) and ecological (light and nutrient treatments) effects on species densities. Almost one fifth (18.6%) of the variation in species density could be explained by the variance in |Δ*x*| (Figure 3a; dLm, dRn, dSp, dWc). Including trait change in the statistical model accurately led to reductions for the estimated importance of ecological effects by a total of 14.4% (from 37% to 26.6% for the amount of shading and from 31% to 27.1% for the nutrient concentrations) and the reduction of 4.2% of the random effects (from 18.7% to 15% for time and from 13.3% to 12.8% for replicates) (Figure 3). While |Δ*x*| of the species Lm, Sp and Wc showed a similarly low impact (3.7%, 3.4%, and 3.7% respectively), |Δ*x*| of species Rn showed a higher impact on species density, 7.8%, which can be explained by its high |Δ*x*| changes over time (see Appendix S4 for data). The 95% HPDI for main effect sizes showed a negative effect of light shading for species density, while nutrient concentration had a positive estimated effect (Figure 3c). Changes in |Δ*x*| impacted densities in distinct ways for intra-and interspecific comparisons (Figure 3c). The change in frond size of species Rn (dRn) had a negative impact on Rn density, while change in Wc frond size (dWc) had a positive estimated effect on species Sp and Lm (Figure 3c). We show the estimated effect of trait change for species counts in Figure 3d, which reflects potentially strong impacts in some particular species and environmental treatment combinations, and weak effects with high uncertainty for many species and environmental treatments.

**Figure 3.**
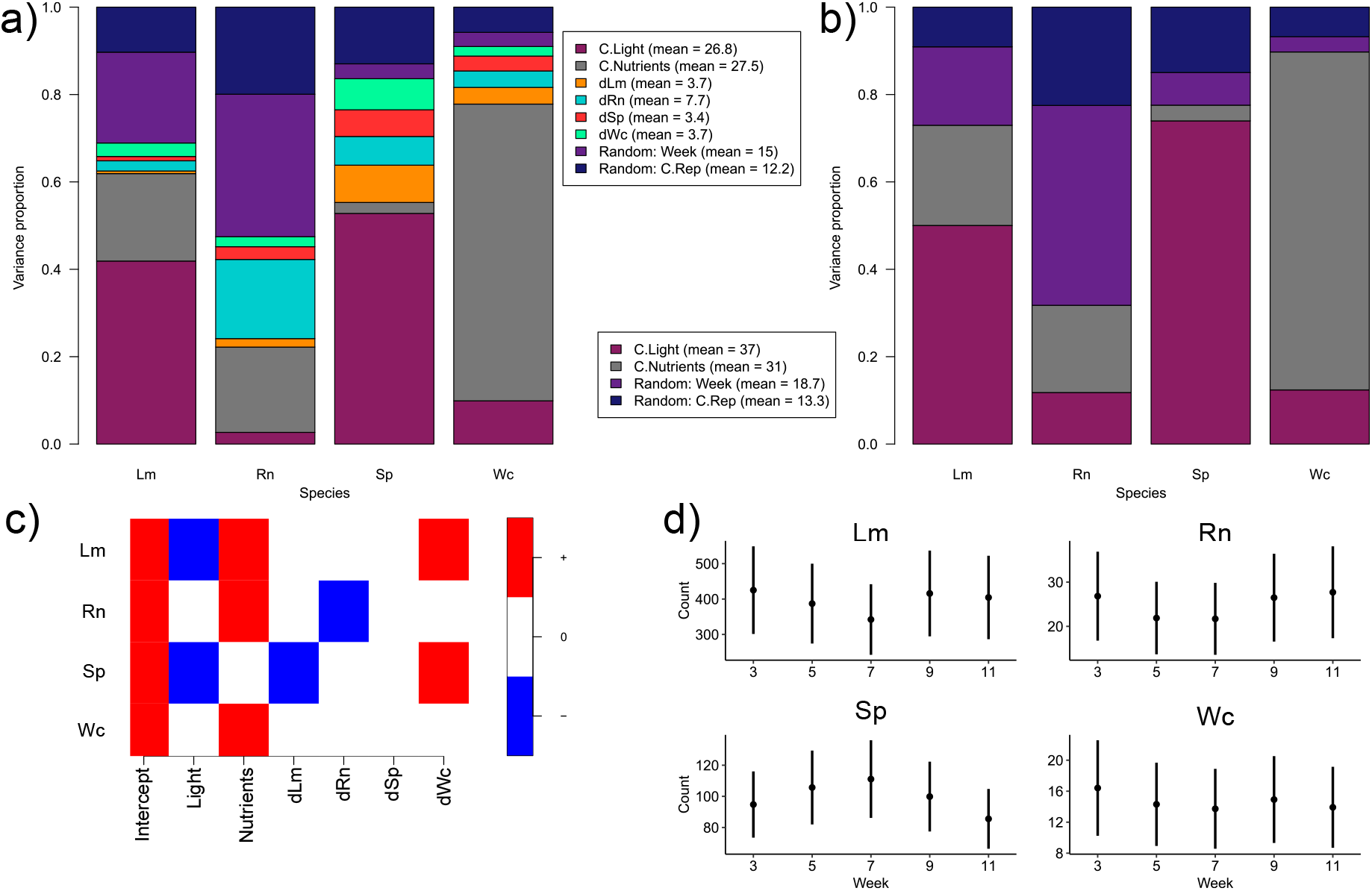
The estimated effects of species trait changes and environmental treatments for densities of experimental aquatic plants (species Lm, Rn, Sp, and Wc). The variation partition for the HMSC model with (a) and without (b) trait change |Δ*x*| is shown, as well as (c) the posterior estimated direction of effect (negative or positive; effect sizes with 95% HPDI that included 0 are colored white). The model estimated negative impacts of light, positive impacts of nutrients, negative intraspecific effects of |Δ*x*_*Rn*_| for *Iog*(*N*_*Rn*_), negative interspecific effects of |Δ*x*_*Lm*_| for *Iog*(*N*_*Sp*_), and positive interspecific effects of |Δ*x*_*Wc*_| for *Iog*(*N*_*Sp*_) and *Iog*(*N*_*Lm*_). (d) The posterior effect size estimates were used to predict species densities at each time step of the experiment. Here we show results only for the main effects of |Δ*x*| (for each of the four species, averaged across all of the other treatment effects). This captures the expected change in density due to trait evolution in all species.

## 6 Discussion

The analysis of community composition often requires statistical models that can consider the combined influences of environmental drivers and spatial connectivity. A more difficult, yet critical, element to incorporate is species interactions. Although the causal role of species interactions is difficult to interpret from statistical models (but see (Dubart et al. 2022) (Luo et al. 2022)) in systems with few species), joint species distribution models (JSDMs) have implemented spatial and temporal random effects that can effectively capture these species interactions using a latent variable approach (however it is important to note that the resulting species covariance matrices also reflect other processes besides species interactions, including unmeasured environmental effects; (Poggiato et al. 2021)). One element that has not been fully implemented into joint species distribution models is dynamic trait evolution (see (Benito Garzón et al. 2019) for a model that does consider reaction norms from in-situ reciprocal transplant experiments). We used an existing JSDM, and developed simulation models of evolving populations in increasingly complex scenarios to determine whether the signal of trait evolution could be estimated from data and compared to other drivers of community structure. We also applied this statistical model (HMSC; (Ovaskainen et al. 2017) (Tikhonov et al. 2020)) to an existing empirical dataset (from the experimental study of community eco-evolutionary dynamics in aquatic plants of (Jewell and Bell 2023)) to show how HMSC can be used to model and predict the changes in species composition over time as a function of environmental drivers, trait change, and temporal random effects. Overall, there is no barrier to including dynamic trait change as a fixed effect in a JSDM model, and the resulting effect size coefficient estimates can be used to (i) estimate the relative importance of trait evolution for community composition and (ii) predict the impacts of future trait change in the system.

At the same time, fixed effects for the impacts of trait evolution on population size should be interpreted with caution. For example, our simulations reveal instances where trait evolution may occur at the same time as large changes in population size, and this co-occurrence isn’t always causal (e.g. when selection is weak but trait heritability is high, and species are introduced at low initial population size). In this instance, the regression coefficient from the model may be negative and strong. By using simulations of trait evolution and population growth in simple systems (1-and 2-species systems, at single and simplified multi-patch scales; Figure 1; Appendix S1), we can better interpret effect size estimates. However, these precautions are not unique to including trait evolution as a predictor of community composition. Previous research has shown that the effect size coefficients from linear and AR models using *N*_*i,t−*1_ to predict *N*_*i,t*_ or *N*_*j,t*_ are not equivalent to the intra-and interspecific interaction coefficients that determine species density dependence and should not automatically be used to interpret the direction and magnitude of species inter-actions ((Kloppers and Greeff 2013) (Certain et al. 2018) (Mühlbauer et al. 2020) (Olivença et al. 2021)). Our findings for using HMSC to estimate the impact of trait evolution (|Δ*x*_*i*_|) for species abundances are similar -(i) the relative importance of trait evolution estimated by a variation partition accurately reflects the relative strength of simulated species interactions vs. magnitude of trait change and (ii) the coefficients greatly increase the predictive ability of the statistical model for simulated data (Appendix S1). HMSC is therefore a useful tool for understanding the role that evolution can play in community dynamics.

This was also observed when using HMSC to evaluate the importance of evolution for metacommunity structure, using simulated data from a model of population growth, competition, dispersal, and trait evolution in a spatially structured habitat (Figure 2a). First of all, the simulations themselves revealed an intriguing role of adaptive evolution for rescuing individual species from extinction (i.e. (Gomulkiewicz and Holt 1995)) and as a consequence greatly increasing biodiversity at the local and the landscape scale (i.e. community rescue, (Low-Décarie et al. 2015) (Bell et al. 2019); Figure 2b). The recovery of species via local adaptation had consequent effects for coexistence, as species that were initially mal-adapted could recover in population size and exert strong competitive effects on other species. Because HMSC can accurately estimate the relative importance of trait evolution for community composition (Appendix S3), we were thus able to test hypotheses about the drivers of increased or decreased impacts of trait evolution for community structure. Our simulation models fit to HMSC confirmed our hypotheses that the importance of evolution would decrease with increasing landscape connectivity (modeled as species dispersal rate *d*) and the overall speed of adaptive evolution (modeled as heritability *h*^2^). Previous theoretical models have shown that these two properties can have a strong influence on resulting community composition ((Urban et al. 2008) (Loeuille and Leibold 2008) (Urban and De Meester 2009) (De Meester et al. 2016)), but our demonstration that the HMSC statistical model can accurately estimate the proportion of variation explained by trait evolution can allow hypotheses from theoretical models to be tested using empirical data, with effect size estimates (regression coefficients from the model) that can be compared across different study systems.

The HMSC model was also applied to experimental aquatic plants, to estimate the importance of trait change (frond size), light levels, and nutrient composition for abundances of four species (Lm, Rn, Sp and Wc) over 11 weeks. This model currently does not take full advantage of the reciprocal transplant data (as implemented in the method of Jewell and Bell (2023)) and thus doesn’t separate the plastic elements of trait change (another key difference in our approach is that we use species densities as our response, not the community average trait value; our results are therefore not meant to be compared directly with those of Jewell and Bell (2023)). However, the HMSC model does not require derivation of new categories for effect sizes (this is the case for Pantel et al. (2015), Hattich et al. (2022), and Jewell and Bell (2023)), nor must data be modelled as normal. The HMSC model also allows for the flexible implementation of random effects according to the study design. As we show in Appendix S2 (see also Ovaskainen and Abrego (2020)), changes in population growth rate (Δ*r*) or community mean trait values (*x*) can also be fit as response variables, which may be more useful for mechanistic inference in some analyses. We also show an additional advantage of HMSC, which is the ability to predict species responses to environmental gradients and trait change (i.e. Figure 3c). It should therefore be possible to use models such as HMSC to study how evolution and trait change impact species densities under changing environmental conditions, or how the impact of trait change might depend on specific environmental conditions. These interactions are possible to consider in the Bayesian HMSC model. Researchers are thus free to fit a JSDM or HMSC model to match their study design, which is not the case for existing formulations of eco-evolutionary effect size estimates that use an ANOVA (i.e. Ellner et al. (2011); Govaert et al. (2016)).

As observed in the simulation results, correctly interpreting the estimated posterior effect sizes of evolution requires a thorough understanding of the experimental system. For example we often observed a negative effect of trait change for species density in model simulations, and this was also observed in the empirical results. However these effects derived from different mechanisms. In the simulations, trait change was greater when a species trait was further away from the optimal value (*E*_*k*_ *− x*_*ikt*_), where they also had lower growth rates (the measure of fitness in our models) and thus lower population size. Species with lower trait distance (*E*_*k*_ *− x*_*ikt*_) and thus higher population size had smaller trait changes. For this reason, HMSC estimates a negative effect of trait change |Δ*x*| for population size. Such effects should also be observed in studies of evolutionary rescue, where a species adapts to an initially lethal environment and avoids extinction via a gradually increasing growth rate ((Gomulkiewicz and Holt 1995); demonstrated in Appendix S3). In the empirical data this negative effect size estimate resulted as the species Rn invested early in the experiment in a larger frond size (Appendix S4). This may have been a tradeoff with a lower investment into reproduction, while still maintaining a similar total frond area on the water surface to capture the reduced light.

There has been a strong general interest in estimating effect sizes of evolution for ecological processes (Hairston Jr et al. 2005); (Ellner et al. 2011); (Pantel et al. 2015); (Barbour et al. 2022)), but in many empirical studies these estimates focus on one or a few species where the impacts of their evolution is strong enough to play a clear and detectable role in the corresponding ecological response variable of interest (e.g. population growth rate, community diversity, likelihood of food web collapse). There is no general knowledge for overall distributions of effect sizes for the impact of evolution for species composition in communities, nor are there studies that evaluate drivers of shifts in these distributions of effect sizes in complex communities (see (Hermann et al. 2024) for estimates in a predator-prey system). We present here distributions of effect sizes of trait change and evolution for species abundances (i.e. posterior estimates of regression coefficients; Figure 3c; Appendix S1), which allow estimation of the relative importance of evolution and trait change for community composition (i.e. Figure 1 c,f,i; Figure 2 c; Figure 3 a,b) in simulated and experimental aquatic plant communities. We expect that effect size coefficients can be useful for better understanding the relative importance of evolution as a driver of community composition.

An increasing number of studies collect and analyze high-resolution time series of populations and trait values for single (Rudman et al. 2022) and multiple (Thomas et al. 2018) (Gibert et al. 2023) (Han et al. 2023) (Shen et al. 2023) species. With increasing access to ecological and evolutionary data comes an increase in the number of interacting processes that should be considered to identify causal mechanisms that structure this data (Pantel and Becks 2023). HMSC, and joint species distribution models generally, can be expanded to further include dynamic trait change that occurs during the course of community ecological studies. Two critically important additions that will increase the application to field or experimental observation data will be (1) bidirectional causality and eco-evolutionary feedbacks and (2) the relative contribution of plasticity and genotypic evolution for trait change (i.e. (Gibert et al. 2022), (Jewell and Bell 2023)). It will also be necessary to improve knowledge of the distributions of eco-evolutionary effects and how these are influenced by causal processes that change the speed of evolution or the strength of species interactions ((Fronhofer et al. 2023)). Although these interacting eco-evolutionary dynamic processes represent an increased degree of complexity for data analysis and interpretation, these mechanistic processes can be translated into simpler statistical models to estimate ecological and evolutionary drivers of community composition.

## Notes

https://github.com/jhpantel/ecoevo-hmsc/tree/aRxiv

### Competing Interest Statement

The authors have declared no competing interest.

## References

Abrego, N., B. Crosier, P. Somervuo, N. Ivanova, A. Abrahamyan, A. Abdi, K. Hämäläinen, et al. 2020. Fungal communities decline with urbanization—more in air than in soil. The ISME Journal 14:2806–2815.

Abrego, N., P. Niittynen, J. Kemppinen, and O. Ovaskainen. 2024. Disentangling the role of intraspecific trait variation in community assembly with joint species-trait distribution modelling. Authorea Preprints.

Antão, L. H., B. Weigel, G. Strona, M. Hällfors, E. Kaarlejärvi, T. Dallas, Ø. H. Opedal, et al. 2022. Climate change reshuffles northern species within their niches. Nature Climate Change 12:587–592.

Barbour, M. A., D. J. Kliebenstein, and J. Bascompte. 2022. A keystone gene underlies the persistence of an experimental food web. Science 376:70–73.

Bell, G., V. Fugère, R. Barrett, B. Beisner, M. Cristescu, G. Fussmann, J. Shapiro, et al. 2019. Trophic structure modulates community rescue following acidification. Proceedings of the Royal Society B: Biological Sciences 286:20190856.

Benito Garzón, M., T. M. Robson, and A. Hampe. 2019. ΔTraitSDMs: Species distribution models that account for local adaptation and phenotypic plasticity. New Phytologist 222:1757–1765.

Beverton, R. J., and S. J. Holt. 1957. On the dynamics of exploited fish populations (Vol. 11). Springer Science & Business Media.

Certain, G., F. Barraquand, and A. Gårdmark. 2018. How do MAR(1) models cope with hidden nonlinearities in ecological dynamics? Methods in Ecology and Evolution 9:1975–1995.

De Meester, L., J. Vanoverbeke, L. J. Kilsdonk, and M. C. Urban. 2016. Evolving perspectives on monopolization and priority effects. Trends in Ecology & Evolution 31:136–146.

Devarajan, K., T. L. Morelli, and S. Tenan. 2020. Multi-species occupancy models: Review, roadmap, and recommendations. Ecography 43:1612–1624.

Doser, J. W., A. O. Finley, M. Kéry, and E. F. Zipkin. 2022. spOccupancy: An r package for single-species, multi-species, and integrated spatial occupancy models. Methods in Ecology and Evolution 13:1670–1678.

Dubart, M., J.-P. Pointier, P. Jarne, and P. David. 2022. Niche filtering, competition and species turnover in a metacommunity of freshwater molluscs. Oikos 2022:e09157.

Ellner, S. P., M. A. Geber, and N. G. Hairston Jr. 2011. Does rapid evolution matter? Measuring the rate of contemporary evolution and its impacts on ecological dynamics. Ecology Letters 14:603–614.

Fronhofer, E. A., D. Corenblit, J. N. Deshpande, L. Govaert, P. Huneman, F. Viard, P. Jarne, et al. 2023. Eco-evolution from deep time to contemporary dynamics: The role of timescales and rate modulators. Ecology Letters 26:S91–S108.

Gibert, J. P., Z.-Y. Han, D. J. Wieczynski, S. Votzke, and A. Yammine. 2022. Feedbacks between size and density determine rapid eco-phenotypic dynamics. Functional Ecology 36:1668–1680.

Gibert, J. P., D. J. Wieczynski, Z.-Y. Han, and A. Yammine. 2023. Rapid eco-phenotypic feedback and the temperature response of biomass dynamics. Ecology and Evolution 13:e9685.

Gomulkiewicz, R., and R. D. Holt. 1995. When does evolution by natural selection prevent extinction? Evolution 49:201–207.

Govaert, L., J. H. Pantel, and L. De Meester. 2016. Eco-evolutionary partitioning metrics: Assessing the importance of ecological and evolutionary contributions to population and community change. Ecology Letters 19:839–853.

Hairston Jr, N. G., S. P. Ellner, M. A. Geber, T. Yoshida, and J. A. Fox. 2005. Rapid evolution and the convergence of ecological and evolutionary time. Ecology Letters 8:1114–1127.

Han, Z.-Y., D. J. Wieczynski, A. Yammine, and J. P. Gibert. 2023. Temperature and nutrients drive eco-phenotypic dynamics in a microbial food web. Proceedings of the Royal Society B: Biological Sciences 290:20222263.

Hattich, G. S. I., L. Listmann, L. Govaert, C. Pansch, T. B. H. Reusch, and B. Matthiessen. 2022. Experimentally decomposing phytoplankton community change into ecological and evolutionary contributions. Functional Ecology 36:120–132.

Hermann, R. J., J. H. Pantel, T. Réveillon, and L. Becks. 2024. Range of trait variation in prey determines evolutionary contributions to predator growth rates. Journal of Evolutionary Biology 37:693–703.

Hiltunen, T., and L. Becks. 2014. Consumer co-evolution as an important component of the eco-evolutionary feedback. Nature communications 5:5226.

Jewell, M. D., and G. Bell. 2023. Eco-evolutionary contributions to community trait change in floating aquatic plants. Ecology 104:e4117.

Keppeler, F. W., M. C. Andrade, P. A. A. Trindade, L. M. Sousa, C. C. Arantes, K. O. Winemiller, O.P. Jensen, et al. 2022. Early impacts of the largest amazonian hydropower project on fish communities. Science of The Total Environment 838:155951.

Kloppers, P. H., and J. C. Greeff. 2013. Lotka–volterra model parameter estimation using experiential data. Applied Mathematics and Computation 224:817–825.

Lasky, J. R., M. B. Hooten, and P. B. Adler. 2020. What processes must we understand to forecast regional-scale population dynamics? Proceedings of the Royal Society B: Biological Sciences 287:20202219.

Leibold, M. A., F. J. Rudolph, F. G. Blanchet, L. De Meester, D. Gravel, F. Hartig, P. Peres-Neto, et al. 2022. The internal structure of metacommunities. Oikos 2022.

Leslie, P. H., and J. C. Gower. 1958. The properties of a stochastic model for two competing species. Biometrika 45:316–330.

Lion, S. 2018. Theoretical approaches in evolutionary ecology: Environmental feedback as a unifying perspective. The American Naturalist 191:21–44.

Little, C. J., and F. Altermatt. 2018. Do priority effects outweigh environmental filtering in a guild of dom-inant freshwater macroinvertebrates? Proceedings of the Royal Society B: Biological Sciences 285:20180205.

Loeuille, N., and M. A. Leibold. 2008. Evolution in metacommunities: On the relative importance of species sorting and monopolization in structuring communities. The American Naturalist 171:788–799.

Low-Décarie, E., M. Kolber, P. Homme, A. Lofano, A. Dumbrell, A. Gonzalez, and G. Bell. 2015. Community rescue in experimental metacommunities. Proceedings of the National Academy of Sciences 112:14307–14312.

Luo, M., S. Wang, S. Saavedra, D. Ebert, and F. Altermatt. 2022. Multispecies coexistence in fragmented landscapes. Proceedings of the National Academy of Sciences 119:e2201503119.

Mouquet, N., and M. Loreau. 2003. Community patterns in source-sink metacommunities. The American Naturalist 162:544–557.

Mühlbauer, L. K., M. Schulze, W. S. Harpole, and A. T. Clark. 2020. gauseR: Simple methods for fitting lotka-volterra models describing gause’s “struggle for existence”. Ecology and Evolution 10:13275–13283.

Olivença, D. V., J. D. Davis, and E. O. Voit. 2021. Comparison between lotka-volterra and multivariate autoregressive models of ecological interaction systems. bioRxiv.

Osmond, M. M., and C. de Mazancourt. 2013. How competition affects evolutionary rescue. Philosophical Transactions of the Royal Society B: Biological Sciences 368:20120085.

Ovaskainen, O., and N. Abrego. 2020. Joint species distribution modelling: With applications in r. Cambridge University Press.

Ovaskainen, O., G. Tikhonov, A. Norberg, F. Guillaume Blanchet, L. Duan, D. Dunson, T. Roslin, et al. 2017. How to make more out of community data? A conceptual framework and its implementation as models and software. Ecology Letters 20:561–576.

Pantel, J. H., and L. Becks. 2023. Statistical methods to identify mechanisms in studies of eco-evolutionary dynamics. Trends in Ecology & Evolution 38:760–772.

Pantel, J. H., C. Duvivier, and L. D. Meester. 2015. Rapid local adaptation mediates zooplankton community assembly in experimental mesocosms. Ecology Letters 18:992–1000.

Pichler, M., and F. Hartig. 2021. A new joint species distribution model for faster and more accurate inference of species associations from big community data. Methods in Ecology and Evolution 12:2159–2173.

Poggiato, G., T. Münkemüller, D. Bystrova, J. Arbel, J. S. Clark, and W. Thuiller. 2021. On the interpretations of joint modeling in community ecology. Trends in Ecology & Evolution 36:391–401.

Pollock, L. J., R. Tingley, W. K. Morris, N. Golding, R. B. O’Hara, K. M. Parris, P. A. Vesk, et al. 2014. Understanding co-occurrence by modelling species simultaneously with a joint species distribution model (JSDM). Methods in Ecology and Evolution 5:397–406.

R Core Team. 2024. R: A language and environment for statistical computing. R Foundation for Statistical Computing, Vienna, Austria.

Rudman, S. M., S. I. Greenblum, S. Rajpurohit, N. J. Betancourt, J. Hanna, S. Tilk, T. Yokoyama, et al. 2022. Direct observation of adaptive tracking on ecological time scales in <i>drosophila</i>. Science 375:eabj7484.

Sandal, L., V. Grøtan, B.-E. Sæther, R. P. Freckleton, D. G. Noble, and O. Ovaskainen. 2022. Effects of density, species interactions, and environmental stochasticity on the dynamics of british bird communities. Ecology 103:e3731.

Shen, C., K. Lemmen, J. Alexander, and F. Pennekamp. 2023. Connecting higher-order interactions with ecological stability in experimental aquatic food webs. Ecology and Evolution 13:e10502.

terHorst, C. P., J. T. Lennon, and J. A. Lau. 2014. The relative importance of rapid evolution for plantmicrobe interactions depends on ecological context. Proceedings of the Royal Society B: Biological Sciences 281:20140028.

Thomas, M. K., S. Fontana, M. Reyes, M. Kehoe, and F. Pomati. 2018. The predictability of a lake phytoplankton community, over time-scales of hours to years. Ecology Letters 21:619–628.

Tikhonov, G., Ø. H. Opedal, N. Abrego, A. Lehikoinen, M. M. J. de Jonge, J. Oksanen, and O. Ovaskainen. 2020. Joint species distribution modelling with the r-package hmsc. Methods in Ecology and Evolution 11:442–447.

Tikhonov, G., O. Ovaskainen, J. Oksanen, M. de Jonge, O. Opedal, and T. Dallas. 2022. Hmsc: Hierarchical model of species communities.

Toju, H., M. Yamamichi, P. R. Guimaraes Jr, J. M. Olesen, A. Mougi, T. Yoshida, and J. N. Thompson. 2017. Species-rich networks and eco-evolutionary synthesis at the metacommunity level. Nature Ecology & Evolution 1:0024.

Urban, M. C., and L. De Meester. 2009. Community monopolization: Local adaptation enhances priority effects in an evolving metacommunity. Proceedings of the Royal Society B: Biological Sciences 276:4129– 4138.

Urban, M. C., M. A. Leibold, P. Amarasekare, L. De Meester, R. Gomulkiewicz, M. E. Hochberg, C. A. Klausmeier, et al. 2008. The evolutionary ecology of metacommunities. Trends in ecology & evolution 23:311–317.

Weigel, B., N. Kotamäki, O. Malve, K. Vuorio, and O. Ovaskainen. 2023. Macrosystem community change in lake phytoplankton and its implications for diversity and function. Global Ecology and Biogeography 32:295–309.

