## Appendix S1 for "Detection of eco-evolutionary dynamics in communities using Joint Species Distribution Models"

### Appendix S1. Linear regression models and non-linear population dynamics

Jelena H. Pantel* Ruben J. Hermann†

26 November, 2024

#### 1 Model A: One species, logistic growth

Population growth over time in a single species is first modelled using a Beverton-Holt (discrete-time, logistic) model (Beverton and Holt (1957)), using an intra-specific competition coefficient for density-dependent growth (Hart and Marshall (2013)) - thus $\alpha = 1/K$.

$$N_{i,t+1} = \frac{r_i N_{i,t}}{1 + \alpha_{ii} N_{i,t}}$$

Note that in this model, the system is at equilibrium when $N_{i,t+1} = N_{i,t}$, and therefore:

$$N^* = N^* \frac{r_i}{1 + \alpha_{ii} N^*}$$

$$1 = \frac{r_i}{1 + \alpha_{ii} N^*}$$

$$N^* = \frac{r_i - 1}{\alpha_{ii}}$$

##### 1.1 Population dynamics simulation

We simulate population growth using $r_i = 1.67$ and $\alpha_{ii} = 0.00125$, with an initial population size $N_{1,0} = 10$. This is implemented using the `disc_log` function from the `ecoevor` package https://github.com/jhpantel/ecoevoR.

```
set.seed(42)
# Simulate initial species population growth
N1.0 <- 10
r1.0 <- 1.67
alpha.11 <- 0.00125
# Simulation of model for t time steps
t <- 30
N <- rep(NA, t)
```

*Laboratoire Chrono-environnement,UMR 6249 CNRS-UFC, 16 Route de Gray, 25030 Besançon cedex, France,

†University of Duisburg-Essen, Universitätsstraße 5, 45141 Essen, Germany,

```
N[1] <- N1.0
for (i in 2:t) {
    N[i] <- disc_log(r = r1.0, N0 = N[i - 1], alpha = alpha.11)
}
```

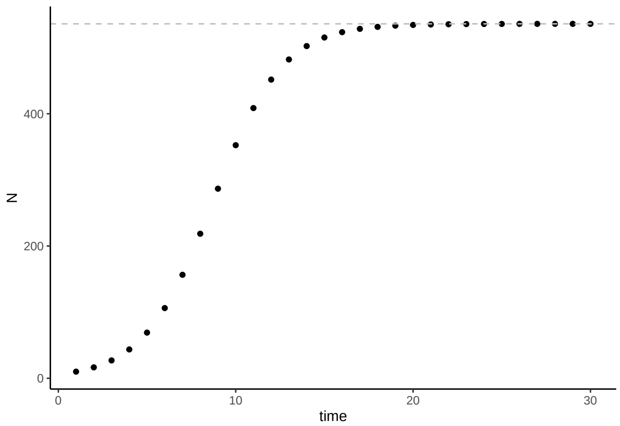


Figure 1: Population size $N$ over time $t$ for a discrete-time logistic growth model, with parameters $r_i = 1.67$, $N_{1,0} = 10$, and $\alpha_{11} = 0.00125$.

#### 1.2 Linear statistical model

We fit the population time series data to a first-order auto-regressive model to predict $N_{t+1}$ as a function of $N_t$, and compare that to a linear regression. We use ln-N after Ives (1995), as also discussed in Certain et al. (2018) and Olivença et al. (2021).

$$N_{t+1} = \beta_0 + \beta_1 N_t + \epsilon_t$$

```
# Fit the model
m.1.ar <- arima(x = log(N), order = c(1, 0, 0), include.mean = T, method = "CSS")
m.1.lm <- lm(log(dat$N[2:t]) ~ log(dat$N[1:(t - 1)]))
# plotting the series along with the fitted values
m.1.ar.fit <- log(N) - residuals(m.1.ar)
m.1.lm.fit <- log(dat$N[2:t]) - m.1.lm$resid
dat$ar1.fit <- m.1.ar.fit
dat$lm.fit <- NA
dat$lm.fit[2:t] <- m.1.lm.fit
```

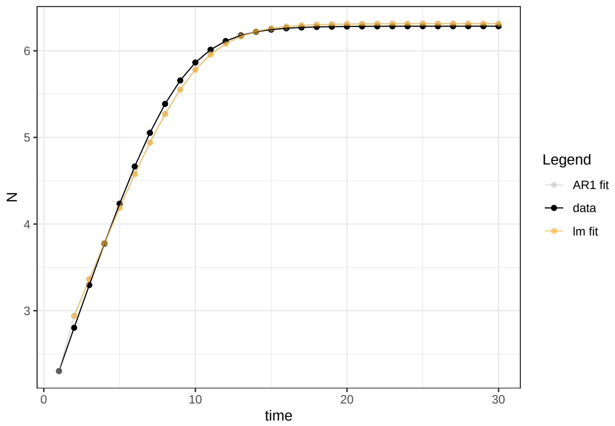


Figure 2: Population size over time (black line) with fitted values from a first-order autoregressive (grey) and linear regression (orange) model.

The linear model is a good fit, and $N_{t+1}$ and $N_t$ are well-represented by a linear function:

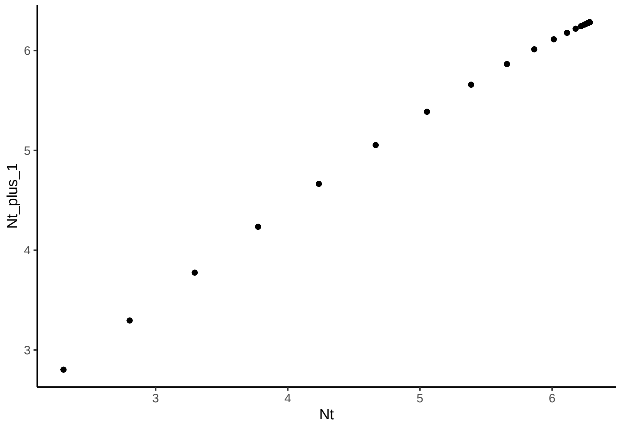


Figure 3: Population size (logarithm) at one time step $N_{t+1}$ as a function of log-population size in the previous time step $N_t$.

We also examine density dependence by plotting $\Delta N = N_{t+1} - N_t$ vs. $N_t$:

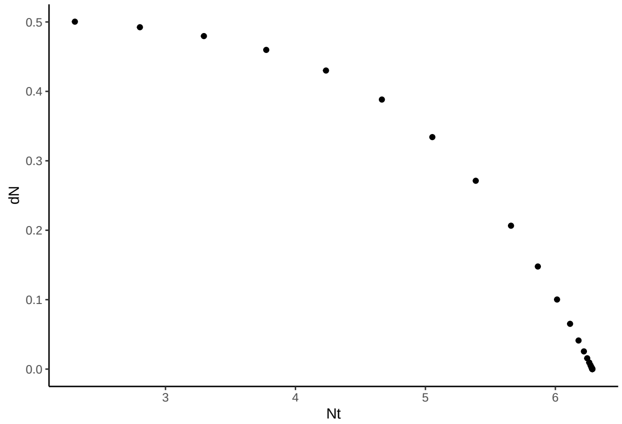


Figure 4: Change in population size from one time step to the next $\Delta N$ as a function of $N_{t+1}$

#### 1.3 Bayesian linear statistical model: HMSC

We can estimate the same model parameters using HMSC:

```
# prepare data in HMSC format
Y <- as.matrix(log(dat$N[2:t]))
XData <- data.frame(x = log(dat$N[1:(t - 1)]))
m.1.hmsc <- Hmsc(Y = Y, XData = XData, XFormula = ~x)
# Bayesian model parameters
nChains <- 2
thin <- 5
samples <- 1000
transient <- 500 * thin
verbose <- 500 * thin
# sample MCMC
m.1.sample <- sampleMcmc(m.1.hmsc, thin = thin, sample = samples, transient = transient,
    nChains = nChains, verbose = verbose)
```

```
m.post.hmsc <- convertToCodaObject(m.1.sample)
summary(m.post.hmsc$Beta)
#>
#> Iterations = 2505:7500
#> Thinning interval = 5
#> Number of chains = 2
#> Sample size per chain = 1000
#>
#> 1. Empirical mean and standard deviation for each variable,
#>    plus standard error of the mean:
#>
#>                                   Mean      SD  Naive SE Time-series SE
#> B[(Intercept) (C1), sp1 (S1)] 0.9887 0.06022 0.0013466      0.0013461
#> B[x (C2), sp1 (S1)]           0.8473 0.01055 0.0002358      0.0002421
#>
#> 2. Quantiles for each variable:
#>
#>                                   2.5%    25%   50%    75%  97.5%
```

```
#> B[(Intercept) (C1), sp1 (S1)] 0.8711 0.9476 0.990 1.0309 1.0983
#> B[x (C2), sp1 (S1)]           0.8274 0.8399 0.847 0.8543 0.8686
plot(m.post.hmsc$Beta)
```

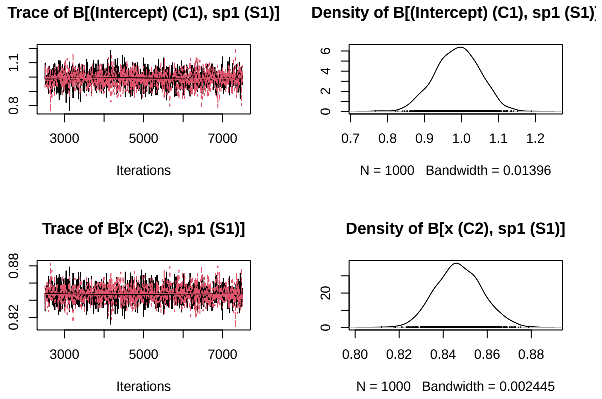


These estimates match well with those from the AR1 and linear model:

```
# AR1 coefficients (recall that the intercept is the term below multipled by 1
# - phi1)
m.1.ar$coef
#>       ar1 intercept
#> 0.8473883 6.4763916
m.1.ar$coef[2] * (1 - m.1.ar$coef[1])
#> intercept
#> 0.9883733
# linear model
summary(m.1.lm)$coefficients[1:2, 1:2]
#>                          Estimate  Std. Error
#> (Intercept)             0.9883734 0.053842285
#> log(dat$N[1:(t - 1)]) 0.8473883 0.009454972
# Bayesian estimates
summary(m.post.hmsc$Beta)$statistics[1:2, 1:2]
#>                                    Mean         SD
#> B[(Intercept) (C1), sp1 (S1)] 0.9887325 0.06022001
#> B[x (C2), sp1 (S1)]           0.8472922 0.01054677
```

```
Gradient <- constructGradient(m.1.sample, focalVariable = "x", ngrid = 29)
predY <- predict(m.1.sample, Gradient = Gradient, expected = TRUE)
plotGradient(m.1.sample, Gradient, pred = predY, showData = T, measure = "Y", main = "",
    xlab = "N_t", ylab = "predicted N_t+1")
```

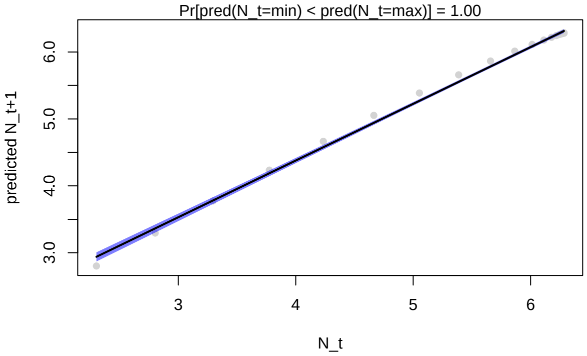


Figure 5: Observed (grey) and model-fit (blue) values for population size at time t (x-axis) and t+1 (y-axis) using HMSC.

```
lm_dat <- data.frame(cbind(log(dat$N[2:t]), log(dat$N[1:(t - 1)])))
colnames(lm_dat) <- c("Nt1", "Nt")
ggplot(lm_dat, aes(Nt, Nt1)) + stat_summary(fun.data = mean_cl_normal) + geom_smooth(method = "lm") +
    theme_classic()
```

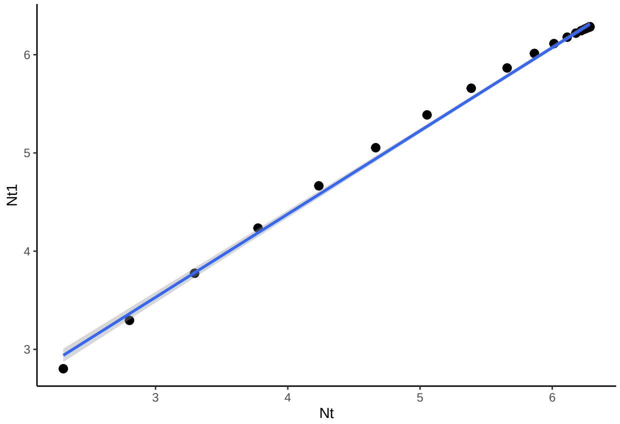


Figure 6: Observed (grey) and model-fit (blue) values for population size at time t (x-axis) and t+1 (y-axis) using lm.

##### 1.4 Conclusions

In this example, a first-order auto-regressive model works well, bypassing the need to estimate logistic growth parameters $r_i$ and $\alpha_{ii}$. The density-dependence dynamics ($\Delta N \sim f(N_t)$) show an overall declining trend over time. The Bayesian estimation implemented in HMSC gives good parameter estimates.

#### 2 Model B: One species, logistic growth, environmental covariate

We now consider using a linear model to analyze population growth when the species growth rate is impacted by a single environmental covariate.

#### 2.1 Growth depends on environment

First we add environment-dependent growth rate. The growth rate $r_i$ becomes:

$$r_i = \hat{W} e^{-(E - x_{i,t})^2}$$

Here, $\hat{W}$ is the maximal population growth rate (set to 1.67 as above), $E$ is the local environmental trait optimum value, and $x_{i,t}$ is species $i$ trait value at time $t$. We see that if $E = x_{i,t}$ then the growth rate is at the value $r = 1.67$. Here, we begin with $E = x_{i,t} = 0.8$, then simulate the environment $E$ value fluctuating randomly over time, and finally use a linear model to fit $E$ as a covariate. We implement the simulation using the `disc_log_E` function from the `ecoevor` package.

```
# Simulate initial species population growth with environment fluctuations
N1.0 <- 10
r1.0 <- 1.67
alpha.11 <- 0.00125
E.0 <- 0.8
x1.0 <- 0.8
# Simulation of model for t time steps
t <- 40
N <- rep(NA, t)
N[1] <- N1.0
E <- rep(NA, t)
E[1] <- E.0
for (i in 2:t) {
    N[i] <- disc_log_E(r = r1.0, N0 = N[i - 1], alpha = alpha.11, E = E[i - 1], x = x1.0)
    E[i] <- E[i - 1] + rnorm(1, 0, 0.1)
}
```

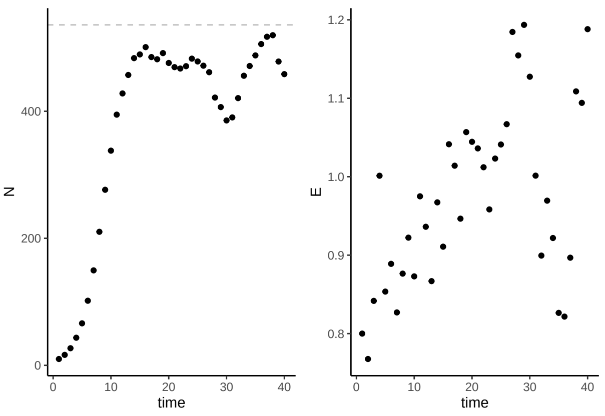


Figure 7: Population size $N$ over time $t$ for a discrete-time logistic growth model, with parameters $r_i = 1.67$, $N_{1,0} = 14$, and $\alpha_{11} = 0.00125$. The value of E over time is also shown.

#### 2.2 Linear statistical model with environmental covariate

We now include environment $E$ as a covariate in the linear model:

$$N_t = \beta_0 + \beta_1 N_{t-1} + \beta_2 E_{t-1} + \epsilon_t$$

```
# Fit the model
m.2.ar <- arima(x = log(N), order = c(1, 0, 0), include.mean = T, method = "CSS",
    xreg = E)
m.2.lm <- lm(log(dat$N[2:t]) ~ log(dat$N[1:(t - 1)]) + log(E[1:(t - 1)]))
# plotting the series along with the fitted values
m.2.ar.fit <- log(N) - residuals(m.2.ar)
m.2.lm.fit <- log(dat$N[2:t]) - m.2.lm$resid
dat$ar2.fit <- m.2.ar.fit
dat$lm2.fit <- NA
dat$lm2.fit[2:t] <- m.2.lm.fit
```

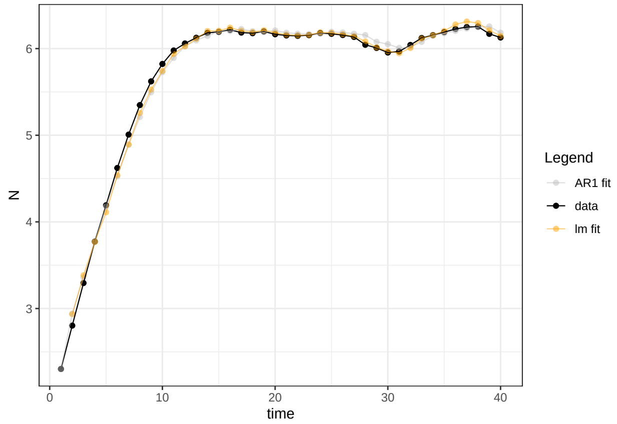


Figure 8: Population size over time (black line) with fitted values from a first-order autoregressive model (gray) and a linear model (orange).

The linear model is a good fit when including the environmental covariate. $N_{t+1}$ and $N_t$ can still be captured by a linear relationship. However we see that the relationship between $N_{t+1}$ and $E_t$ is non-linear.

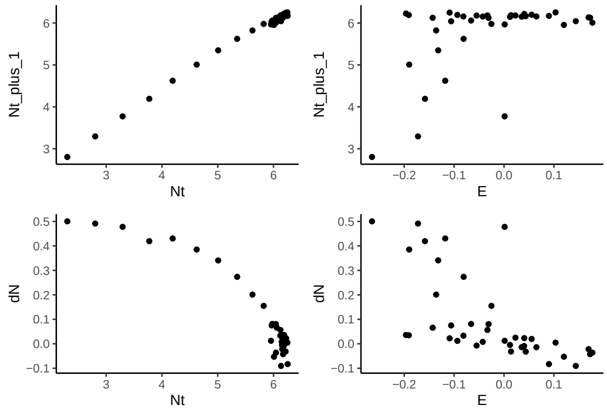


Figure 9: Population size (logarithm) at one time step $N_{t+1}$ as a function of log-population size in the previous time step $N_t$ and of environmental value $E_t$. Also shown are change in population size $\Delta N$ with population size and E.

This tells us that the lm is good for predictions, but not for inference (for capturing well the relationsip between the predictor and response variable). The use of linear relationships in JSDMs is discussed in

(Ingram et al. 2020), and in many applications (e.g. (Erickson and Smith 2023)) quadratic terms are used, which create bell-shaped response curves that may better match species with optimal niches (as opposed to linear, monotonically increasing relationships between population size and environmental predictors). We thus include a quadratic term for $E_t$ to provide a better fit to the data.

```
df <- data.frame(cbind(log(dat$N[2:t]), log(dat$N[1:(t - 1)]), E[1:(t - 1)], E[1:(t -
    1)]^2))
colnames(df) <- c("Nt1", "Nt", "E", "Esq")
m.2.lm <- lm(Nt1 ~ Nt + E + Esq, data = df)
```

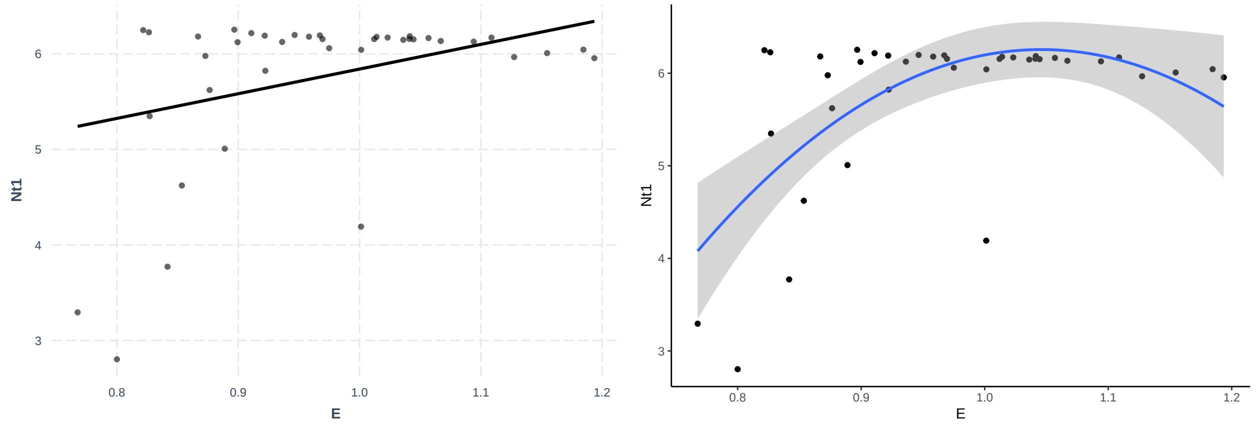


Figure 10: Population size (logarithm) at one time step $N_{t+1}$ as a function of environment E. Linear and quadratic fits are shown.

#### 2.3 Bayesian linear statistical model: HMSC

We can estimate the same model parameters using HMSC:

```
# prepare data in HMSC format
Y <- as.matrix(log(dat$N[2:t]))
XData <- df
m.2.hmsc <- Hmsc(Y = Y, XData = XData, XFormula = ~Nt + E + Esq)
# Bayesian model parameters
nChains <- 2
thin <- 5
samples <- 1000
transient <- 500 * thin
verbose <- 500 * thin
# sample MCMC
m.2.sample <- sampleMcmc(m.2.hmsc, thin = thin, sample = samples, transient = transient,
    nChains = nChains, verbose = verbose)
```

```
m2.post.hmsc <- convertToCodaObject(m.2.sample)
summary(m2.post.hmsc$Beta)
#>
#> Iterations = 2505:7500
#> Thinning interval = 5
#> Number of chains = 2
```

```
#> Sample size per chain = 1000
#>
#> 1. Empirical mean and standard deviation for each variable,
#>    plus standard error of the mean:
#>
#>                                    Mean      SD  Naive SE Time-series SE
#> B[(Intercept) (C1), sp1 (S1)] -0.1172 0.72629 0.0162404      0.0162433
#> B[Nt (C2), sp1 (S1)]           0.8521 0.01108 0.0002477      0.0002143
#> B[E (C3), sp1 (S1)]            2.5496 1.53081 0.0342299      0.0342367
#> B[Esq (C4), sp1 (S1)]         -1.4924 0.76626 0.0171341      0.0171372
#>
#> 2. Quantiles for each variable:
#>
#>                                    2.5%     25%     50%     75%   97.5%
#> B[(Intercept) (C1), sp1 (S1)] -1.5785 -0.5918 -0.1164  0.3659 1.33237
#> B[Nt (C2), sp1 (S1)]           0.8302  0.8448  0.8520  0.8594 0.87430
#> B[E (C3), sp1 (S1)]           -0.4942  1.5454  2.5787  3.5439 5.62748
#> B[Esq (C4), sp1 (S1)]         -3.0327 -1.9918 -1.5099 -0.9818 0.06678
bayesplot::mcmc_trace(m2.post.hmsc$Beta)
```

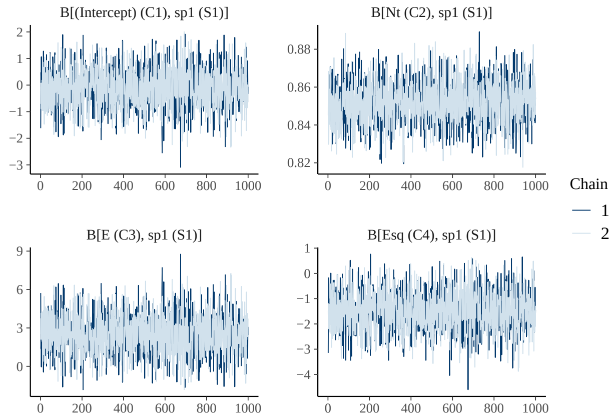


```
bayesplot::mcmc_areas(m2.post.hmsc$Beta, area_method = c("equal height"))
```

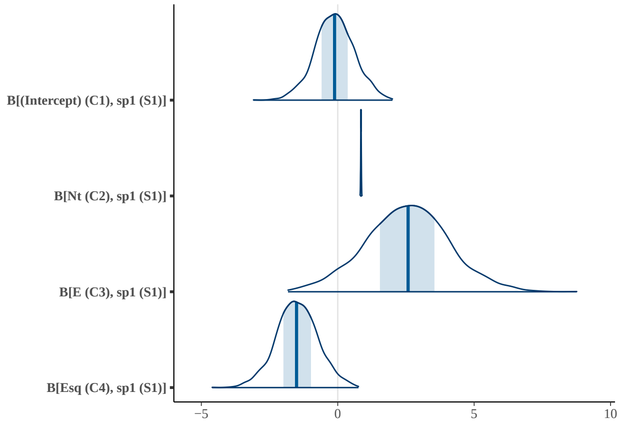


These estimates match well with those from the AR1 and linear model:

```r
# AR1 coefficients (recall that the intercept is the term below multipled by 1
# - phi1)
m.2.ar$coef
#>         ar1   intercept           E
#>  0.83925564  6.29776167 -0.02674129
m.2.ar$coef[2] * (1 - m.2.ar$coef[1])
#> intercept
#>   1.01233
# linear model
summary(m.2.lm)$coefficients[1:4, 1:2]
#>               Estimate Std. Error
#> (Intercept) -0.1317811 0.63418671
#> Nt           0.8523506 0.01002202
#> E            2.5819242 1.33687286
#> Esq         -1.5113693 0.66907748
# Bayesian estimates
summary(m2.post.hmsc$Beta)$statistics[1:4, 1:2]
#>                                     Mean         SD
#> B[(Intercept) (C1), sp1 (S1)] -0.1172053 0.72629078
#> B[Nt (C2), sp1 (S1)]           0.8521043 0.01107945
#> B[E (C3), sp1 (S1)]            2.5496453 1.53080925
#> B[Esq (C4), sp1 (S1)]         -1.4923591 0.76625968
```

We recall that the interpretation of the coefficients in an arimaX (arima with covariates) model is difficult. They do not give the impact on $N_t$ per unit increase in X as in a regression. So we do not interpret the causation implied by the coefficient in the arimaX model. In the regression model, we can see that $E$ has a positive impact on $N_t$.

```r
Gradient <- constructGradient(m.2.sample, focalVariable = "E", non.focalVariables = list(Nt = list(2),
    Esq = list(2)), ngrid = 39)
# Esq is manually constructed as gradient-produced E^2
Gradient$XDataNew$Esq <- Gradient$XDataNew$E^2
predY <- predict(m.2.sample, XData = Gradient$XDataNew, expected = TRUE)
plotGradient(m.2.sample, Gradient, pred = predY, showData = T, measure = "Y", main = "",
    xlab = "E_t", ylab = "predicted N_t+1")
```

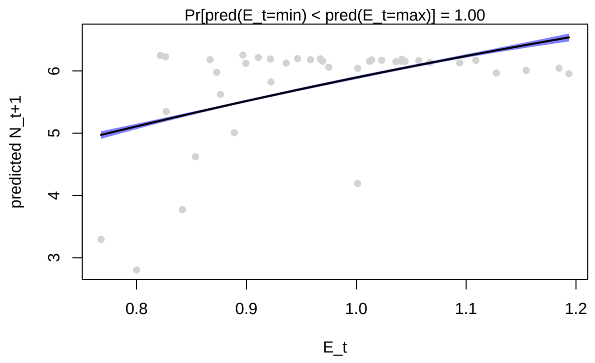


Figure 11: Observed (grey) and model-fit (blue) values for population size at time t+1 (y-axis) as a function of environment $E_t$ (x-axis).

##### 2.4 Conclusions

In this example, the linear regression again works well to describe the impact of $E_t$ for $N_t$ when using the quadratic formulation. The arimaX model works well for fitting and subsequent prediction, but less well for inference about the impacts of $E$. From the quadratic regression terms for $E$, we correctly see that the population size is maximal at the species trait value and decreases away from that value. We will continue to use log-transformed abundance and now introduce quadratic terms for the environmental parameter.

### 3 Model C: Two species, logistic growth, competition

Here we investigate how a second species impacts the inference we can make from linear models.

##### 3.1 Interspecific competition

The growth equation for each species now becomes:

$$N_{i,t+1} = \frac{r_i N_{i,t}}{1 + \alpha_{ii} N_{i,t} + \alpha_{ij} N_{j,t}}$$

We use a distinct growth rate for the 2nd species and introduce the interspecific interaction coefficient $\alpha_{ij}$. The simulation is generated using the `disc_LV_comp` function in the `ecoevor` package.

```r
# Initial conditions
N0 <- c(10, 10)
r <- c(1.67, 1.7)
alpha.11 <- 0.00125
alpha.22 <- 0.00125
alpha.12 <- 0.008
alpha.21 <- 0.008725
alpha <- matrix(c(alpha.11, alpha.21, alpha.12, alpha.22), nrow = 2, byrow = FALSE)
# make sure the interaction matrix is correct Simulation of model for t time
# steps
t <- 40
N <- array(NA, dim = c(t, 2))
N <- as.data.frame(N)
colnames(N) <- c("N1", "N2")
N[1, ] <- N0
for (i in 2:t) {
    N[i, ] <- disc_LV_comp(r = r, N0 = N[i - 1, ], alpha = alpha)
}
```

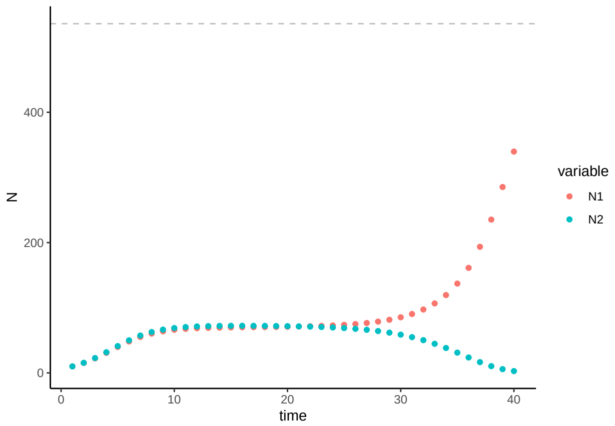


Figure 12: Population size $N$ over time $t$ for a discrete-time logistic growth model, with parameters $r_1 = 1.67$, $r_2 = 1.7$, $N_{i,0} = 10$, $\alpha_{ii} = 0.00125$, $\alpha_{12} = 0.008$, and $\alpha_{21} = 0.008725$.

#### 3.2 Linear statistical model with covariate for both species

We fit each species' population time series to an arimaX and lm with population size of the others species as a covariate.

$$N_{1,t} = \beta_0 + \beta_1 N_{1,t-1} + \beta_2 N_{2,t-1} + \epsilon_t$$

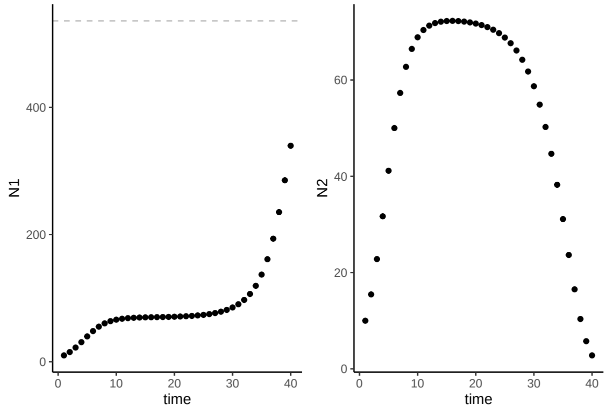


Figure 13: Population size $N$ over time $t$ for a discrete-time logistic growth model, with competition between 2 species ($N_1$ and $N_2$).

```
## Fit the model for Species 1
m.3.ar.n1 <- arima(x = log(N$N1), order = c(1, 0, 0), include.mean = T, method = "CSS",
    xreg = N$N2)
m.3.lm.n1 <- lm(log(dat$N1[2:t]) ~ log(dat$N1[1:(t - 1)]) + log(dat$N2[1:(t - 1)]))
# plotting the series along with the fitted values
m.3.ar.fit.n1 <- log(N$N1) - residuals(m.3.ar.n1)
m.3.lm.fit.n1 <- log(dat$N1[2:t]) - m.3.lm.n1$resid
dat$ar3.fit.n1 <- m.3.ar.fit.n1
dat$lm3.fit.n1 <- NA
```

```
dat$lm3.fit.n1[2:t] <- m.3.lm.fit.n1
## Species 2
m.3.ar.n2 <- arima(x = log(N$N2), order = c(1, 0, 0), include.mean = T, method = "CSS",
    xreg = N$N1)
m.3.lm.n2 <- lm(log(dat$N2[2:t]) ~ log(dat$N2[1:(t - 1)]) + log(dat$N1[1:(t - 1)]))
# plotting the series along with the fitted values
m.3.ar.fit.n2 <- log(N$N2) - residuals(m.3.ar.n2)
m.3.lm.fit.n2 <- log(dat$N2[2:t]) - m.3.lm.n2$resid
dat$ar3.fit.n2 <- m.3.ar.fit.n2
dat$lm3.fit.n2 <- NA
dat$lm3.fit.n2[2:t] <- m.3.lm.fit.n2
```

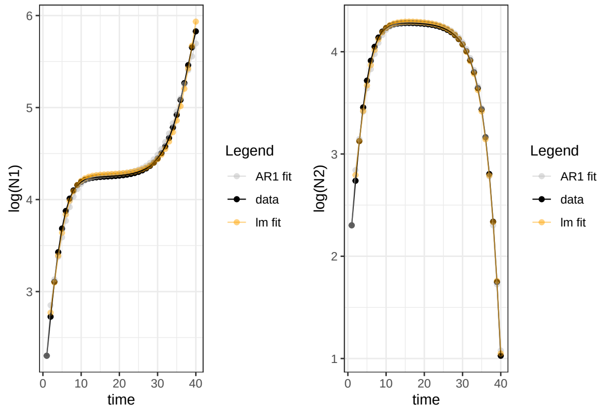


Figure 14: Population size over time (black line) with fitted values from a first-order autoregressive model (gray line) and from a linear model (orange line).

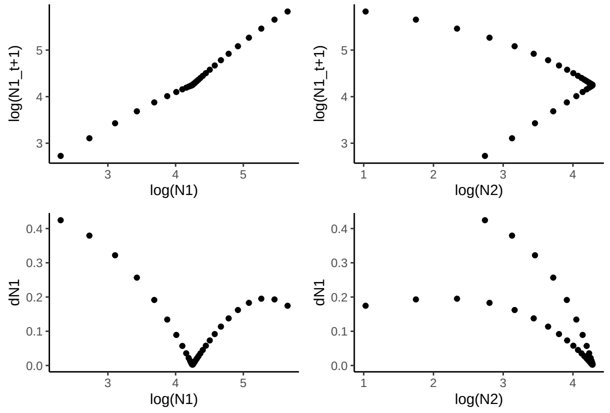


Figure 15: Population size (logarithm) in species 1 at one time step $N_{1,t+1}$ as a function of log-population size of Species 1 and Species 2 in the previous time step $N_t$. Also shown is the change in population size of Species 1 as a function of $N_t$ for both species.

We can see that the population size of Species 2 does not have a linear relationship with that of Species 1.

```
df <- data.frame(cbind(log(dat$N1[2:t]), log(dat$N1[1:(t - 1)]), log(dat$N2[1:(t -
    1)])))
colnames(df) <- c("Nt1", "N1", "N2")
m.3.lm <- lm(Nt1 ~ I(1/N1) + I(1/N2), data = df)
cor(df$Nt1, predict(m.3.lm))
#> [1] 0.9845708
summary(m.3.lm)
#>
#> Call:
#> lm(formula = Nt1 ~ I(1/N1) + I(1/N2), data = df)
#>
#> Residuals:
#>      Min       1Q   Median       3Q      Max
#> -0.30952 -0.04467 -0.03442  0.03430  0.22447
#>
#> Coefficients:
#>             Estimate Std. Error t value Pr(>|t|)
#> (Intercept)   6.4056     0.1060   60.45  < 2e-16 ***
#> I(1/N1)     -12.2863     0.3774  -32.55  < 2e-16 ***
#> I(1/N2)       3.3309     0.2448   13.61 9.24e-16 ***
#> ---
#> Signif. codes:  0 '***' 0.001 '**' 0.01 '*' 0.05 '.' 0.1 ' ' 1
#>
#> Residual standard error: 0.1063 on 36 degrees of freedom
#> Multiple R-squared:  0.9694, Adjusted R-squared:  0.9677
#> F-statistic: 569.8 on 2 and 36 DF,  p-value: < 2.2e-16
```

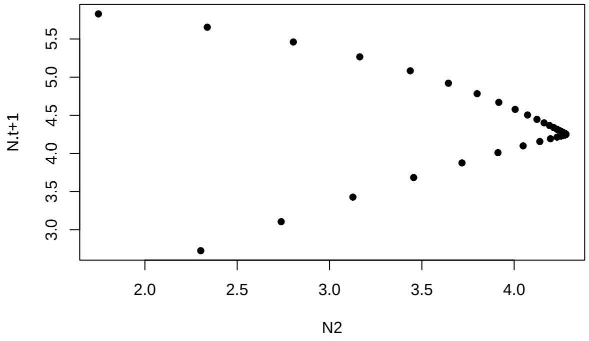


Figure 16: Population size (logarithm) in species 1 at one time step $N_{1,t+1}$ as a function of log-population size of Species 2 in the previous time step $N_t$.

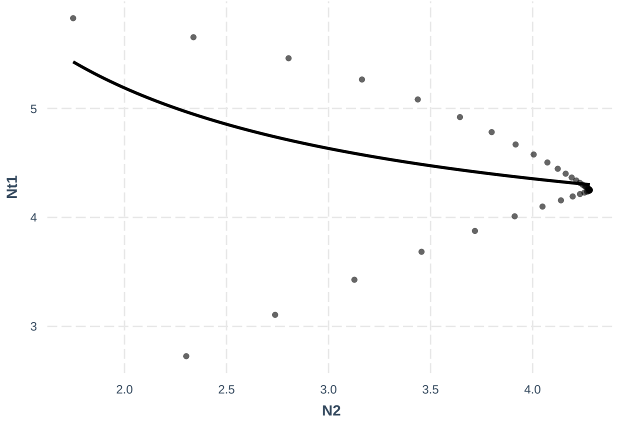


Figure 17: Linear model-predicted population size (logarithm) at one time step $N_{t+1}$ as a function of log-population size of the second species in the previous time step $N_t$. Fitted (line) and observed (points) values are shown.

We can already see that the linear model does not provide a useful fit to the data points (neither does taking the inverse function help). The regression parameters can predict well the population size of the other species, but they are not informative for inference. This has been observed previously (Certain et al. (2018)), and non-linear least squares models (among others) are more often used to estimate parameters for dynamics resulting from these kinds of species interaction models (Kloppers and Greeff (2013); Mühlbauer et al. (2020); Olivença et al. (2021)). The inability of the linear models to reproduce the interaction coefficients is more clearly demonstrated in Certain et al. (2018). However, they do indicate that the slope of the linear model can reflect the direction of effect for the interacting species.

Instead of further exploring techniques to fit to time-series derived from non-linear competition processes, we look more closely at how HMSC manages to fit data emerging from competition dynamics and make inference about species interactions. Instead of estimating full interaction coefficients, or assuming sparse interactions, they instead assume that a reduced number of linear combinations of species abundances that are most relevant to determining future growth rates for species in the community can bypass the 'curse of dimensionality' problem. This is well presented in Ovaskainen et al. (2017). We apply that approach here.

#### 3.3 Bayesian linear statistical model: HMSC

We can fit the data to a linear model using HMSC, with latent variables to capture the species associations. These are implemented by including a random effect at the time level. For this analysis, we do not expect the impact of Species 1 and 2 that results from competition to vary over time - competition will always have negative impacts for species abundances. Our goal is to estimate the overall impact of species covariance across all of the time points. We do not estimate these impacts as a function of the lag in time points. In other words, we assume that each time point has random variance and that species may covary in their response at each time point. We do not consider that species covariance is a function of distance between sampled time points (see Ovaskainen and Abrego (2020), equation 5.9). We instead assume that site loadings $\eta$ are independent among the sampled time points.

```
# prepare data in HMSC format
dat <- as.data.frame(log(N))
dat$time <- 1:t
df <- data.frame(dat[(2:t), -3])
Y <- as.matrix(df)
XData <- data.frame(dat[1:(t - 1), -3])
colnames(XData) <- c(paste0("n", 1:length(r)))
```

```
studyDesign = data.frame(sample = as.factor(1:(t - 1)))
studyDesign$sample <- as.factor(studyDesign$sample)
rL.sample = HmscRandomLevel(units = studyDesign$sample)
m.3.hmsc = Hmsc(Y = Y, XData = XData, XFormula = ~1, studyDesign = studyDesign, ranLevels = list(sample
# Bayesian model parameters
nChains <- 2
thin <- 5
samples <- 1000
transient <- 500 * thin
verbose <- 500 * thin
# sample MCMC
m.3.sample <- sampleMcmc(m.3.hmsc, thin = thin, sample = samples, transient = transient,
    nChains = nChains, verbose = verbose)
#> Computing chain 1
#> Chain 1, iteration 2500 of 7500 (transient)
#> Chain 1, iteration 5000 of 7500 (sampling)
#> Chain 1, iteration 7500 of 7500 (sampling)
#> Computing chain 2
#> Chain 2, iteration 2500 of 7500 (transient)
#> Chain 2, iteration 5000 of 7500 (sampling)
#> Chain 2, iteration 7500 of 7500 (sampling)
```

```
m3.post.hmsc <- convertToCodaObject(m.3.sample)
summary(m3.post.hmsc$Beta)
#>
#> Iterations = 2505:7500
#> Thinning interval = 5
#> Number of chains = 2
#> Sample size per chain = 1000
#>
#> 1. Empirical mean and standard deviation for each variable,
#>    plus standard error of the mean:
#>
#>                                 Mean      SD Naive SE Time-series SE
#> B[(Intercept) (C1), N1 (S1)] 4.34 0.09341 0.002089       0.002039
#> B[(Intercept) (C1), N2 (S2)] 3.81 0.11431 0.002556       0.002555
#>
#> 2. Quantiles for each variable:
#>
#>                                 2.5%   25%   50%   75% 97.5%
#> B[(Intercept) (C1), N1 (S1)] 4.154 4.279 4.341 4.401 4.511
#> B[(Intercept) (C1), N2 (S2)] 3.581 3.731 3.810 3.887 4.035
bayesplot::mcmc_trace(m3.post.hmsc$Beta)
```

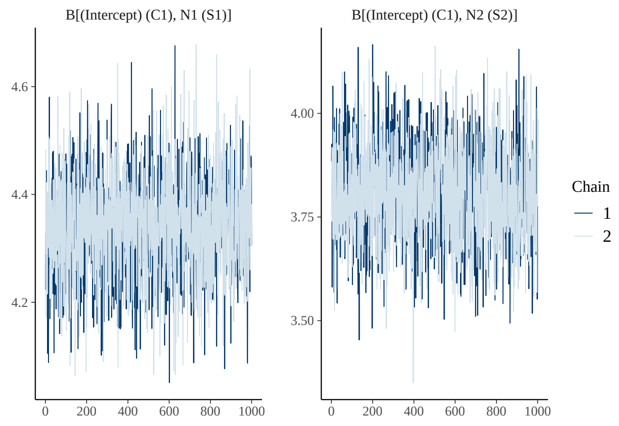


```
bayesplot::mcmc_areas(m3.post.hmsc$Beta, area_method = c("equal height"))
```

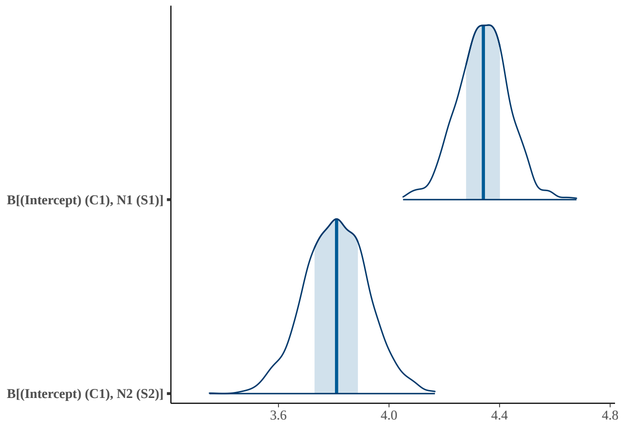


Now we look at the estimates for the species associations:

```
OmegaCor = computeAssociations(m.3.sample)
supportLevel = 0.95
toPlot = ((OmegaCor[[1]]$support > supportLevel) + (OmegaCor[[1]]$support < (1 -
    supportLevel)) > 0) * OmegaCor[[1]]$mean
corrplot::corrplot(toPlot, method = "color", col = colorRampPalette(c("blue", "white",
    "red"))(200), title = paste("random effect level:", m.3.sample$rLNames[1]), mar = c(0,
    0, 1, 0))
```

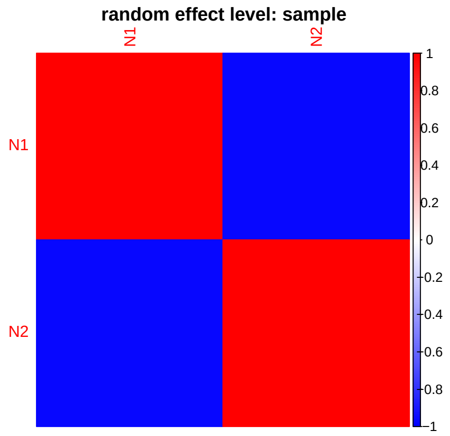

The direction of HMSC model fixed effect estimates matches well with those from the AR1 and linear model.

```
# AR1 coefficients (recall that the intercept is the term below multipled by 1
# - phi1)
m.3.ar.n1$coef
#>         ar1   intercept        N$N2
#>  0.79696422  5.69840618 -0.01881979
m.3.ar.n1$coef[2] * (1 - m.3.ar.n1$coef[1])
#> intercept
#>   1.15698
# linear model
summary(m.3.lm.n1)$coefficients[1:3, 1:2]
#>                          Estimate  Std. Error
#> (Intercept)             0.9839341 0.053084123
#> log(dat$N1[1:(t - 1)])  0.9201864 0.009157214
#> log(dat$N2[1:(t - 1)]) -0.1443693 0.008777527
# Bayesian estimates
summary(m3.post.hmsc$Beta)$statistics[1:2, 1:2]
#>                                 Mean         SD
#> B[(Intercept) (C1), N1 (S1)] 4.340340 0.09340565
#> B[(Intercept) (C1), N2 (S2)] 3.809673 0.11430500
```

The HMSC model correctly shows the negative association between the species (shown here as correlations - JSDM CHapter 7 tells us the intra-specific correlation is always 1).

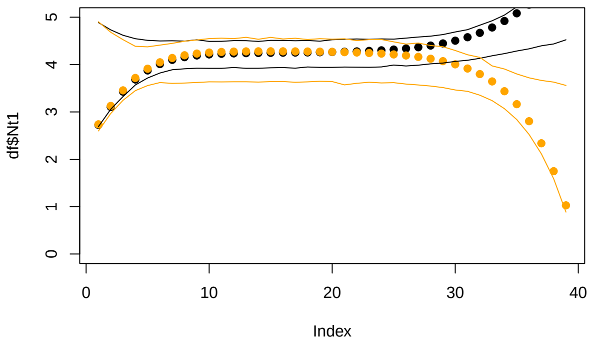


Figure 18: Observed (points) and HMSC model-predicted population size (logarithm) for Species 1 (black) and Species 2 (orange), using environment $E_t$ as a fixed effect predictor and specifying temporal random effects to capture the species interactions. Lines represent the 95% highest probability density interval (HPDI) for predictions at each time point.

To show that the HMSC model correctly allocates variance to the contributing components of the model (here environment, intra-specific and interspecific density-dependence, implemented as temporal random effects), we simulate a new example where the only difference between species is the $\alpha_{ij}$.

```
# Initial conditions
N0 <- c(10, 10)
r <- c(1.7, 1.7)
alpha.11 <- 0.01
```

```
alpha.22 <- 0.01
alpha.12 <- 0.005
alpha.21 <- 0.01
alpha <- matrix(c(alpha.11, alpha.21, alpha.12, alpha.22), nrow = 2, byrow = FALSE)  # careful to make
# Simulation of model for t time steps
t <- 40
N <- array(NA, dim = c(t, 2))
N <- as.data.frame(N)
colnames(N) <- c("N1", "N2")
N[1, ] <- N0
for (i in 2:t) {
    N[i, ] <- disc_LV_comp(r = r, N0 = N[i - 1, ], alpha = alpha)
}
```

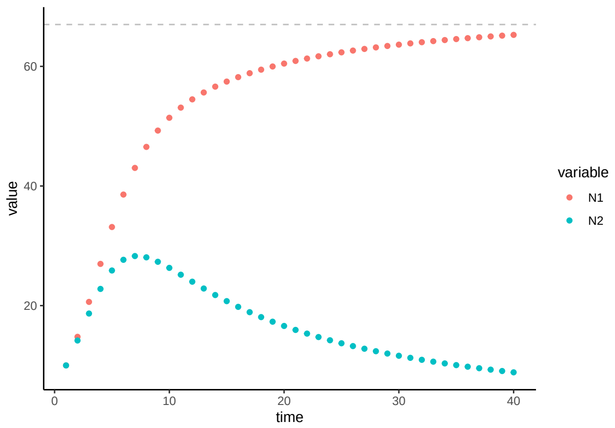


Figure 19: Population size $N$ over time $t$ for a discrete-time logistic growth model with competition, with parameters $r_1 = r_2 = 1.7$, $N_{i,0} = 10$, $\alpha_{ii} = 0.01$, $\alpha_{12} = 0.005$, and $\alpha_{21} = 0.01$.

```
# Plot simulation: ggplot
dat <- as.data.frame(cbind(N$N1, N$N2))
colnames(dat) <- c("N1", "N2")
dat$time <- 1:t
df <- data.frame(cbind(log(dat$N1[2:t]), log(dat$N2[2:t])))
colnames(df) <- c("Nt1", "Nt2")
# prepare data in HMSC format
Y <- as.matrix(cbind(df$Nt1, df$Nt2))
XData <- data.frame(cbind(log(dat$N1[1:(t - 1)]), log(dat$N2[1:(t - 1)])))
studyDesign = data.frame(sample = as.factor(1:(t - 1)))
studyDesign$sample <- as.factor(studyDesign$sample)
rL.sample = HmscRandomLevel(units = studyDesign$sample)
m.4.hmsc = Hmsc(Y = Y, XData = XData, XFormula = ~1, studyDesign = studyDesign, ranLevels = list(sample
# Bayesian model parameters
nChains <- 2
thin <- 5
samples <- 1000
transient <- 500 * thin
verbose <- 500 * thin
# sample MCMC
m.4.sample <- sampleMcmc(m.4.hmsc, thin = thin, sample = samples, transient = transient,
```

```
    nChains = nChains, verbose = verbose)
#> Computing chain 1
#> Chain 1, iteration 2500 of 7500 (transient)
#> Chain 1, iteration 5000 of 7500 (sampling)
#> Chain 1, iteration 7500 of 7500 (sampling)
#> Computing chain 2
#> Chain 2, iteration 2500 of 7500 (transient)
#> Chain 2, iteration 5000 of 7500 (sampling)
#> Chain 2, iteration 7500 of 7500 (sampling)
```

```
m4.post.hmsc <- convertToCodaObject(m.4.sample)
summary(m4.post.hmsc$Beta)
#>
#> Iterations = 2505:7500
#> Thinning interval = 5
#> Number of chains = 2
#> Sample size per chain = 1000
#>
#> 1. Empirical mean and standard deviation for each variable,
#>    plus standard error of the mean:
#>
#>                                   Mean      SD Naive SE Time-series SE
#> B[(Intercept) (C1), sp1 (S1)] 3.974 0.05288 0.001183       0.001153
#> B[(Intercept) (C1), sp2 (S2)] 2.767 0.06124 0.001369       0.001428
#>
#> 2. Quantiles for each variable:
#>
#>                                   2.5%   25%   50%   75% 97.5%
#> B[(Intercept) (C1), sp1 (S1)] 3.867 3.939 3.975 4.009 4.077
#> B[(Intercept) (C1), sp2 (S2)] 2.646 2.726 2.766 2.808 2.884
bayesplot::mcmc_trace(m4.post.hmsc$Beta)
```

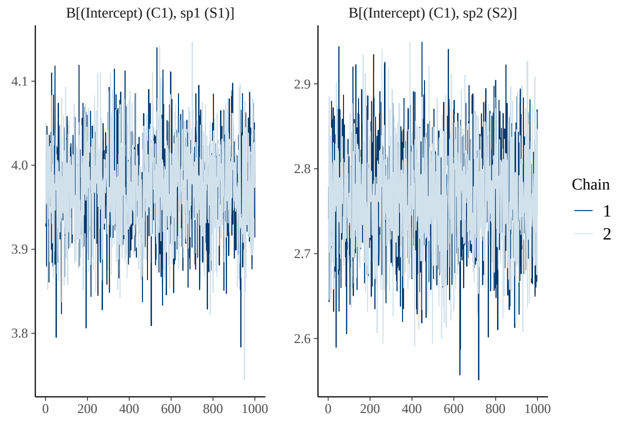


```
bayesplot::mcmc_areas(m4.post.hmsc$Beta, area_method = c("equal height"))
```

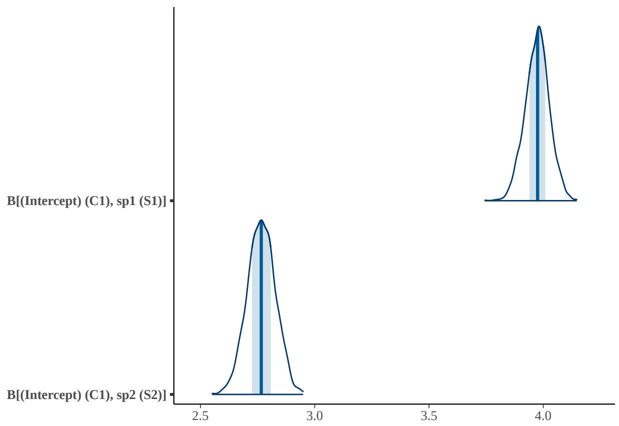


```
OmegaCor = computeAssociations(m.4.sample)
supportLevel = 0.95
toPlot = ((OmegaCor[[1]]$support > supportLevel) + (OmegaCor[[1]]$support < (1 -
    supportLevel)) > 0) * OmegaCor[[1]]$mean
corrplot::corrplot(toPlot, method = "color", col = colorRampPalette(c("blue", "white",
    "red"))(200), title = paste("random effect level:", m.4.sample$rLNames[1]), mar = c(0,
    0, 1, 0))
```

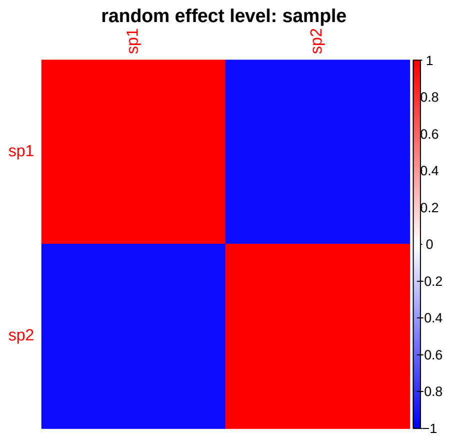


The HMSC model correctly shows the negative association between the species (shown here as correlations - JSDM CHapter 7 tells us the intra-specific correlation is always 1).

```
# explanatory power
preds = computePredictedValues(m.4.sample)
```

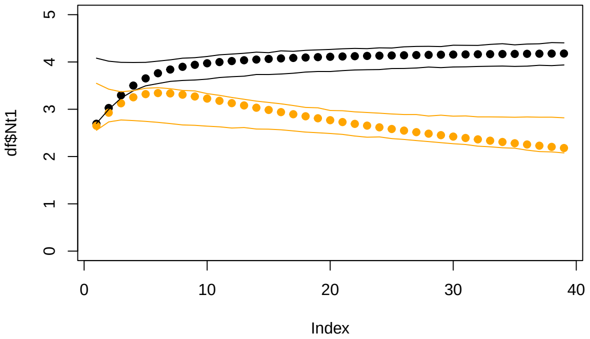


Figure 20: Observed (points) and HMSC model-predicted population size (logarithm) for Species 1 (black) and Species 2 (orange), using environment $E_t$ as a fixed effect predictor and specifying temporal random effects to capture the species interactions. Lines represent the 95% HPDI for predictions at each time point.

The residual associations should not be interpreted as interaction coefficients, but rather used for prediction. We can see here they are effective for this (shown above is the observed and model-predicted 95% highest probability density interval - HPDI).

### 4 Model D: Species growth, competition, with environmental change

The goal of the process was to use HMSC's latent variable approach to study the results of non-linear processes in ecology using a linear model. We can infer the direction and magnitude of the species interaction. We now consider an environmental covariate that changes over time and impacts each species growth. We will use only HMSC from this point forward, as the species interactions represent a departure from what a linear or AR model can manage.

In this example, the local environmental optimum trait value $E = 0.8$, and Species 1 $x_1 = 0.6$, while $x_2 = 0.8$. The simulation is generated using the `disc_LV_E` function in the `ecoevor` package.

```
# Initial conditions
N0 <- c(10, 10)
r <- c(1.7, 1.7)
alpha.11 <- 0.01
alpha.22 <- 0.01
alpha.12 <- 0.005
alpha.21 <- 0.01
alpha <- matrix(c(alpha.11, alpha.21, alpha.12, alpha.22), nrow = 2, byrow = FALSE)
E.0 <- 0.8
x <- c(0.6, 0.8)
# Simulation of model for t time steps
t <- 40
N <- array(NA, dim = c(t, length(N0)))
N <- as.data.frame(N)
colnames(N) <- paste0("N", 1:length(N0))
N[1, ] <- N0
E <- rep(NA, t)
```

```
E[1] <- E.0
for (i in 2:t) {
    N[i, ] <- disc_LV_E(r = r, N0 = N[i - 1, ], alpha = alpha, E = E[i - 1], x = x)
    E[i] <- E[i - 1] + rnorm(1, 0, 0.1)
}
```

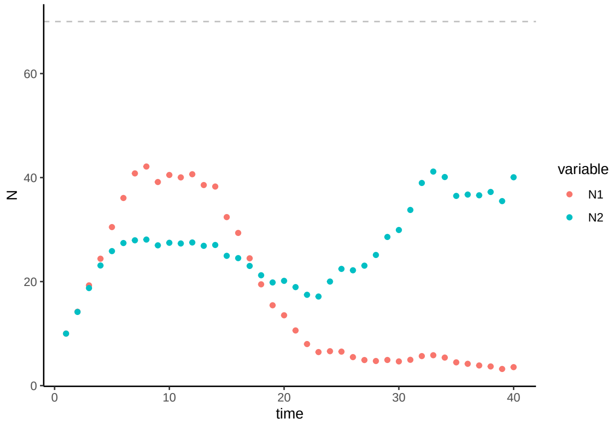


Figure 21: Population size $N$ over time $t$ for a discrete-time logistic growth model with competition and a changing environment, with parameters $r_1 = r_2 = 1.7$, $N_{i,0} = 10$, $\alpha_{ii} = 0.01$, $\alpha_{12} = 0.005$, and $\alpha_{21} = 0.01$.

```
dat <- as.data.frame(log(N))
dat$time <- 1:t
df <- data.frame(dat[(2:t), -3])
Y <- as.matrix(df)
# prepare data in HMSC format
XData <- data.frame(dat[1:(t - 1), -3], E[1:(t - 1)], E[1:(t - 1)]^2)
colnames(XData) <- c(paste0("n", 1:length(r)), "E", "Esq")
studyDesign = data.frame(sample = as.factor(1:(t - 1)))
rL = HmscRandomLevel(units = studyDesign$sample)
m.5.hmsc = Hmsc(Y = Y, XData = XData, XFormula = ~E + Esq, studyDesign = studyDesign,
    ranLevels = list(sample = rL))
# Bayesian model parameters
nChains <- 2
thin <- 5
samples <- 1000
transient <- 500 * thin
verbose <- 500 * thin
# sample MCMC
m.5.sample <- sampleMcmc(m.5.hmsc, thin = thin, sample = samples, transient = transient,
    nChains = nChains, verbose = verbose)
#> Computing chain 1
#> Chain 1, iteration 2500 of 7500 (transient)
#> Chain 1, iteration 5000 of 7500 (sampling)
#> Chain 1, iteration 7500 of 7500 (sampling)
#> Computing chain 2
#> Chain 2, iteration 2500 of 7500 (transient)
#> Chain 2, iteration 5000 of 7500 (sampling)
#> Chain 2, iteration 7500 of 7500 (sampling)
```

```
m5.post.hmsc <- convertToCodaObject(m.5.sample)
summary(m5.post.hmsc$Beta)
#>
#> Iterations = 2505:7500
#> Thinning interval = 5
#> Number of chains = 2
#> Sample size per chain = 1000
#>
#> 1. Empirical mean and standard deviation for each variable,
#>    plus standard error of the mean:
#>
#>                                   Mean     SD Naive SE Time-series SE
#> B[(Intercept) (C1), N1 (S1)]  3.7701 1.2830  0.02869        0.03113
#> B[E (C2), N1 (S1)]            0.9876 2.4710  0.05525        0.06060
#> B[Esq (C3), N1 (S1)]         -1.8970 1.1987  0.02680        0.02944
#> B[(Intercept) (C1), N2 (S2)]  1.4760 0.7564  0.01691        0.01636
#> B[E (C2), N2 (S2)]            3.7344 1.5249  0.03410        0.03307
#> B[Esq (C3), N2 (S2)]         -1.8451 0.7474  0.01671        0.01629
#>
#> 2. Quantiles for each variable:
#>
#>                                  2.5%     25%    50%    75%   97.5%
#> B[(Intercept) (C1), N1 (S1)]  1.27165  2.9198  3.764  4.588  6.4240
#> B[E (C2), N1 (S1)]           -4.02211 -0.6079  1.056  2.593  5.8051
#> B[Esq (C3), N1 (S1)]         -4.23771 -2.6601 -1.899 -1.167  0.4855
#> B[(Intercept) (C1), N2 (S2)]  0.03592  0.9845  1.465  1.960  2.9788
#> B[E (C2), N2 (S2)]            0.59060  2.7442  3.777  4.717  6.6489
#> B[Esq (C3), N2 (S2)]         -3.25030 -2.3306 -1.860 -1.364 -0.3263
bayesplot::mcmc_trace(m5.post.hmsc$Beta)
```

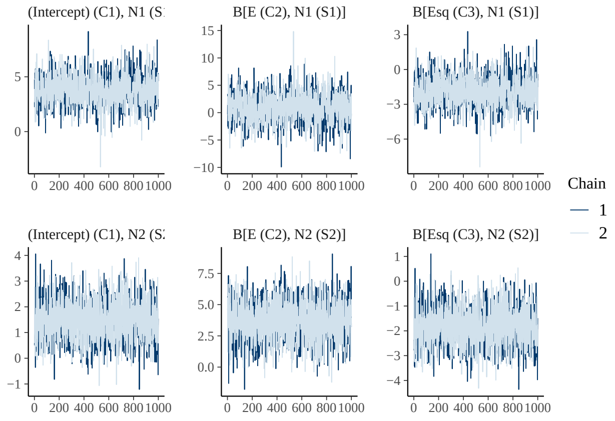


124

```
bayesplot::mcmc_areas(m5.post.hmsc$Beta, area_method = c("equal height"))
```

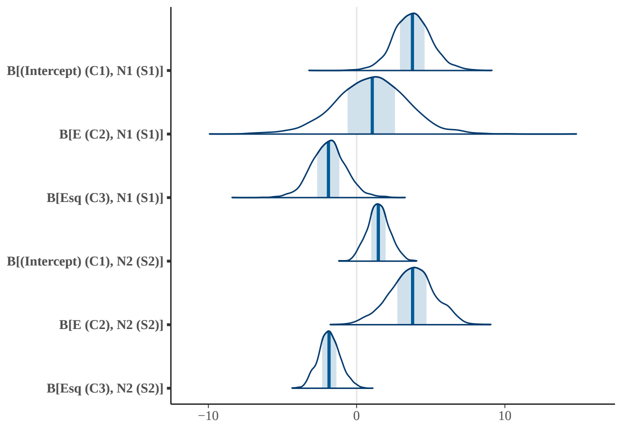


```
# Bayesian estimates
summary(m5.post.hmsc$Beta)$statistics[1:4, 1:2]
#>                                    Mean        SD
#> B[(Intercept) (C1), N1 (S1)]  3.7701166 1.2829827
#> B[E (C2), N1 (S1)]            0.9876064 2.4710204
#> B[Esq (C3), N1 (S1)]         -1.8969594 1.1987431
#> B[(Intercept) (C1), N2 (S2)]  1.4759806 0.7564327
```

```
OmegaCor = computeAssociations(m.5.sample)
supportLevel = 0.95
toPlot = ((OmegaCor[[1]]$support > supportLevel) + (OmegaCor[[1]]$support < (1 -
    supportLevel)) > 0) * OmegaCor[[1]]$mean
corrplot::corrplot(toPlot, method = "color", col = colorRampPalette(c("blue", "white",
    "red"))(200), title = paste("random effect level:", m.5.sample$rLNames[1]), mar = c(0,
    0, 1, 0))
```

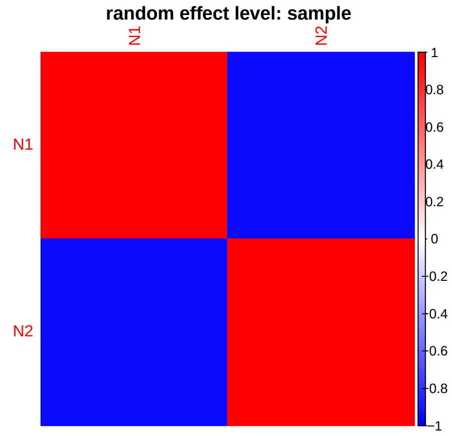


```
Gradient <- constructGradient(m.5.sample, focalVariable = "E", non.focalVariables = list(Esq = list(2))
    ngrid = 39)
predY <- predict(m.5.sample, XData = Gradient$XDataNew, expected = TRUE)
a <- plotGradient(m.5.sample, Gradient, pred = predY, showData = T, measure = "Y",
    index = 1, main = "", xlab = "E_t", ylab = "predicted N1_t+1")
```

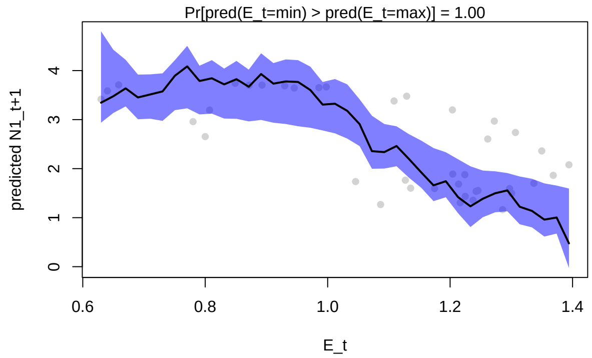


```r
b <- plotGradient(m.5.sample, Gradient, pred = predY, showData = T, measure = "Y",
    index = 2, main = "", xlab = "E_t", ylab = "predicted N2_t+1")
```

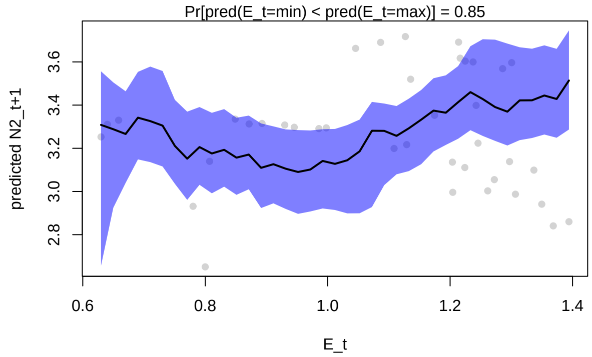


```r
postBeta = getPostEstimate(m.5.sample, parName = "Beta")
plotBeta(m.5.sample, post = postBeta, param = "Mean", supportLevel = 0.7)
```

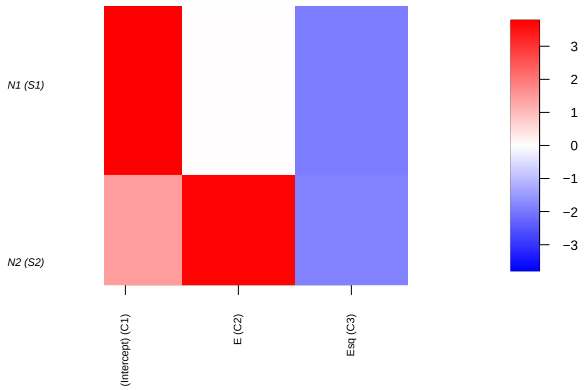


We can see that the Environment has an overall negative impact for Species 1 as $E$ drifts randomly towards more positive values (and thus further from Species 1 trait value). We also see that the residual species association for species 1 and 2 are negative.

We use variation partition to estimate the relative importance of environment and species associations for species abundances.

```r
VP <- computeVariancePartitioning(m.5.sample, group = c(1, 1, 1), groupnames = "Env")
plotVariancePartitioning(m.5.sample, VP, args.legend = list(cex = 0.75, bg = "transparent"))
```

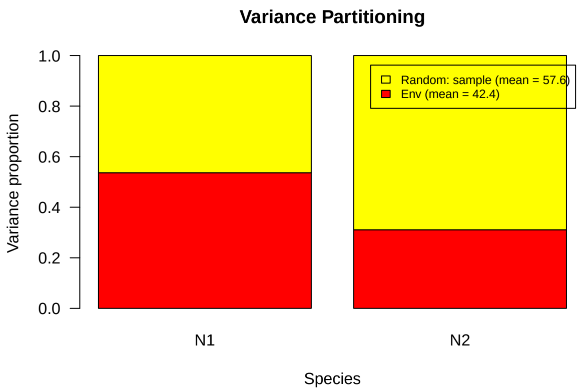


We evaluate whether variation partition correctly detects the relative importance of environment and species interactions by greatly increasing the effects of interspecific competition and decreasing the strength of environmental variation.

```r
# Simulate initial species population growth with environment fluctuations
N0 <- c(10, 10)
r <- c(1.7, 1.7)
alpha.11 <- 0.01
alpha.22 <- 0.01
alpha.12 <- 0.015
alpha.21 <- 0.02
alpha <- matrix(c(alpha.11, alpha.21, alpha.12, alpha.22), nrow = 2, byrow = FALSE)
E.0 <- 0.8
x <- c(0.8, 0.8)
# Simulation of model for t time steps
t <- 40
N <- array(NA, dim = c(t, length(N0)))
N <- as.data.frame(N)
colnames(N) <- paste0("N", 1:length(N0))
N[1, ] <- N0
E <- rep(NA, t)
E[1] <- E.0
for (i in 2:t) {
    N[i, ] <- disc_LV_E(r = r, N0 = N[i - 1, ], alpha = alpha, E = E[i - 1], x = x)
    E[i] <- E[i - 1] + rnorm(1, 0, 0.001)
}
```

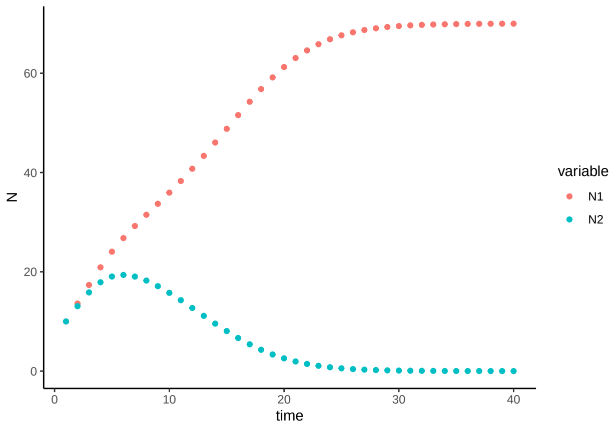


Figure 22: Population size $N$ over time $t$ for a discrete-time logistic growth model with competition and a changing environment, with parameters $r_1 = r_2 = 1.7$, $N_{i,0} = 10$, $\alpha_{ii} = 0.01$, $\alpha_{12} = 0.015$, and $\alpha_{21} = 0.02$.

```
## prepare data in HMSC format
dat <- as.data.frame(log(N))
dat$time <- 1:t
df <- data.frame(dat[(2:t), -3])
Y <- as.matrix(df)
XData <- data.frame(dat[1:(t - 1), -3], E[1:(t - 1)], E[1:(t - 1)]^2)
colnames(XData) <- c(paste0("n", 1:length(r)), "E", "Esq")
studyDesign = data.frame(sample = as.factor(1:(t - 1)))
rL = HmscRandomLevel(units = studyDesign$sample)
m.6.hmsc = Hmsc(Y = Y, XData = XData, XFormula = ~E + Esq, studyDesign = studyDesign,
    ranLevels = list(sample = rL))
# Bayesian model parameters
nChains <- 2
thin <- 5
samples <- 2000
transient <- 100 * thin
verbose <- 500 * thin
# sample MCMC
m.6.sample <- sampleMcmc(m.6.hmsc, thin = thin, sample = samples, transient = transient,
    nChains = nChains, verbose = verbose)
#> Computing chain 1
#> Chain 1, iteration 2500 of 10500 (sampling)
#> Chain 1, iteration 5000 of 10500 (sampling)
#> Chain 1, iteration 7500 of 10500 (sampling)
#> Chain 1, iteration 10000 of 10500 (sampling)
#> Computing chain 2
#> Chain 2, iteration 2500 of 10500 (sampling)
#> Chain 2, iteration 5000 of 10500 (sampling)
#> Chain 2, iteration 7500 of 10500 (sampling)
#> Chain 2, iteration 10000 of 10500 (sampling)
```

```
OmegaCor = computeAssociations(m.6.sample)
supportLevel = 0.95
toPlot = ((OmegaCor[[1]]$support > supportLevel) + (OmegaCor[[1]]$support < (1 -
    supportLevel)) > 0) * OmegaCor[[1]]$mean
```

```
corrplot::corrplot(toPlot, method = "color", col = colorRampPalette(c("blue", "white",
    "red"))(200), title = paste("random effect level:", m.6.sample$rLNames[1]), mar = c(0,
    0, 1, 0))
```

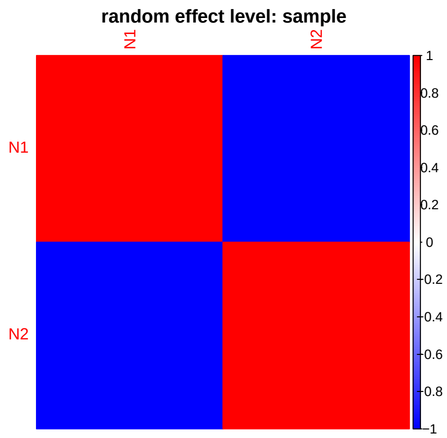


```
OmegaCor
#> [[1]]
#> [[1]]$mean
#>             N1         N2
#> N1   1.0000000 -0.9960107
#> N2 -0.9960107  1.0000000
#>
#> [[1]]$support
#>    N1 N2
#> N1  1  0
#> N2  0  1
```

```
Gradient <- constructGradient(m.6.sample, focalVariable = "E", non.focalVariables = list(Esq = list(2))
    ngrid = 39)
predY <- predict(m.6.sample, XData = Gradient$XDataNew, expected = TRUE)
a <- plotGradient(m.6.sample, Gradient, pred = predY, showData = T, measure = "Y",
    index = 1, main = "", xlab = "E_t", ylab = "predicted N1_t+1")
```

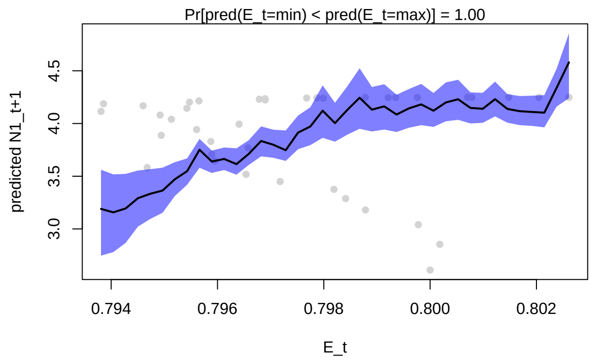

```r
b <- plotGradient(m.6.sample, Gradient, pred = predY, showData = T, measure = "Y",
    index = 2, main = "", xlab = "E_t", ylab = "predicted N2_t+1")
```

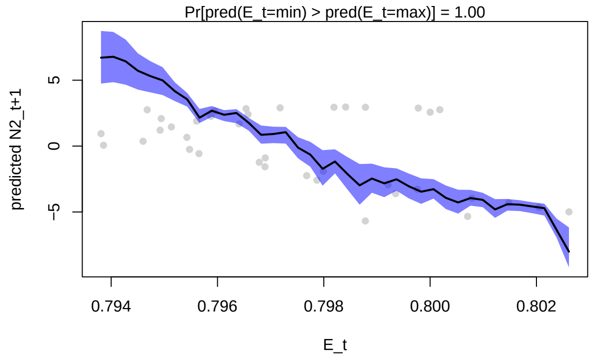


```r
postBeta = getPostEstimate(m.6.sample, parName = "Beta")
plotBeta(m.6.sample, post = postBeta, param = "Mean", supportLevel = 0.7)
```

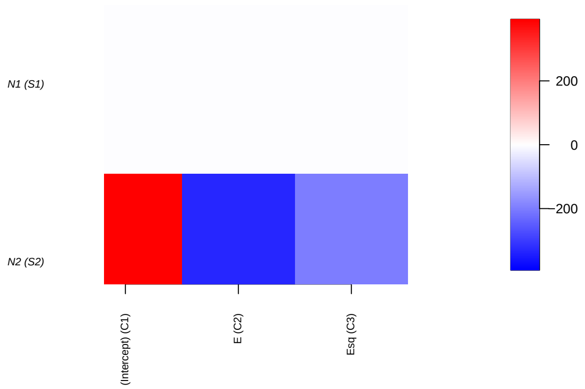


With the reduced impact of the environment, we now see the strong negative associations between the two competing species. We also correctly see a relatively weak impact of environment for Species 1 abundance and a stronger impact for Species 2.

For both species, the relative importance of environment has decreased relative to the species association effects.

```r
VP <- computeVariancePartitioning(m.6.sample, group = c(1, 1, 1), groupnames = "Env")
plotVariancePartitioning(m.6.sample, VP, args.legend = list(cex = 0.75, bg = "transparent"))
```

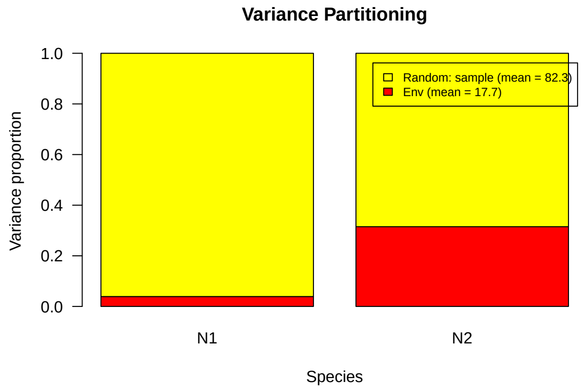


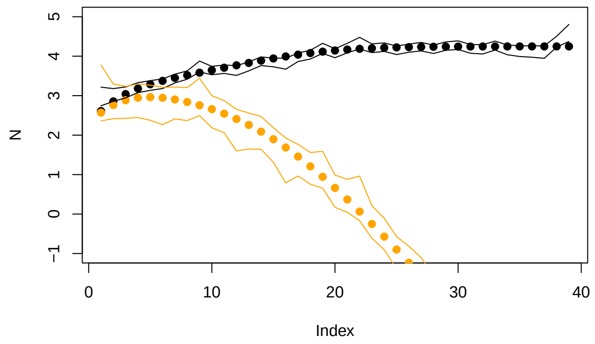


Figure 23: Observed (points) and HMSC model-predicted population size (logarithm) for Species 1 (black) and Species 2 (orange), using environment $E_t$ as a fixed effect predictor and specifying temporal random effects to capture the species interactions. Lines represent the 95% highest probability density interval (HPDI) for predictions at each time point.

### 5 Model E: Species growth, competition, environmental change, and trait evolution

We now consider that trait evolution can occur, with an impact on the species growth rate / fitness in the context of the environment.

#### 5.1 Trait evolution

Previously the population growth rate was fixed as $r_i = \hat{W} e^{-(E-x_{i,t})^2}$. However, we now introduce the quantitative genetic model of evolutionary rescue (Gomulkiewicz and Holt 1995).

The growth equation for each species now becomes:

$$N_{i,t+1} = \frac{\hat{W} e^{\frac{-[(\frac{w+(1-h^2)P}{P+w})(E-x_{i,t})]^2}{2(P+w)}} N_{i,t}}{1+\alpha_{ii}N_{i,t}+\alpha_{ij}N_{j,t}}$$

where $\hat{W}$ is calculated as $\hat{W} = W_{max}\sqrt{(\frac{w}{P+w})}$, $W_{max}$ is the species' maximum per-capita growth rate, $w$ is the width of the Gaussian fitness function (which determines the strength of selection, as increasing values indicate a weaker reduction in fitness with distance from optimum trait value), $P$ is the width of the distribution of the phenotype $x$, and $h^2$ is the heritability of the trait $x$. For the simulation we use $W_{max} = 2$, $P = 1$, and $w = 2$.

The change in the average trait value each time step is given by:

$$d_{i,t+1} = kd_{i,t}$$

where $k = \frac{w+(1-h^2)P}{w+P}$ and $d_{i,t} = E_t - x_{i,t}$.

#### 5.2 Population dynamics simulation

We use one of the same examples as above, with weak strength of interspecific interactions. We use initial trait values that favor the weaker competitor, and we simulate random increasing shifts in the local environment. The simulation uses the `disc_LV_evol` command in the `ecoevor` package.

```
N0 <- c(10, 10)
alpha.11 <- 0.01
alpha.22 <- 0.01
alpha.12 <- 0.005
alpha.21 <- 0.01
alpha <- matrix(c(alpha.11, alpha.21, alpha.12, alpha.22), nrow = 2, byrow = FALSE)
E.0 <- 0.8
x.0 <- c(0.1, 0.8)
P <- 1
w <- 2
Wmax <- 2
h2 <- 1
k <- (w + (1 - h2) * P)/(P + w)
# Simulation of model for t time steps
t <- 40
N <- array(NA, dim = c(t, length(N0)))
N <- as.data.frame(N)
colnames(N) <- paste0("N", 1:length(N0))
N[1, ] <- N0
E <- rep(NA, t)
E[1] <- E.0
x <- array(NA, dim = c(t, length(N0)))
x <- as.data.frame(x)
colnames(x) <- paste0("x", 1:length(N0))
x[1, ] <- x.0
r <- array(NA, dim = c(t, length(N0)))
r <- as.data.frame(r)
colnames(r) <- paste0("r", 1:length(N0))
What <- Wmax * sqrt(w/(P + w))
r[1, ] <- What * exp((-(((w + (1 - h2) * P)/(P + w)) * (E[1] - x[1, ]))^2)/(2 * (P +
    w)))
for (i in 2:t) {
    res <- disc_LV_evol(N0 = N[i - 1, ], alpha = alpha, E = E[i - 1], x = x[i - 1,
        ], P = P, w = w, Wmax = Wmax, h2 = h2)
    N[i, ] <- res$Nt1
```

```
    r[i, ] <- res$r
    # trait change
    d <- E[i - 1] - x[i - 1, ]
    d1 <- k * d
    x[i, ] <- E[i - 1] - d1
    # environmental change
    E[i] <- E[i - 1] + abs(rnorm(1, 0, 0.05))
}
```

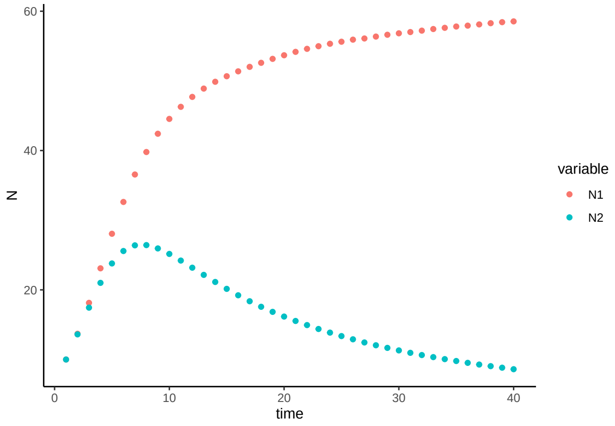


Figure 24: Population size $N$ over time $t$ for a discrete-time logistic growth model with competition, a changing environment, and trait evolution. Model parameters are $N_{i,0} = 10$, $\alpha_{ii} = 0.01$, $\alpha_{12} = 0.005$, and $\alpha_{21} = 0.01$.

165 The overall research goal is to determine the effect of trait evolution for species abundances. We hypothe-
166 size that $|x_{t+1} - x_{i,t}|$ is an important driver of species abundances, alongside the environment and species
167 interactions. We plan to fit a statistical model to estimate the impact of $E$, $E^2$, and $|x_{t+1} - x_{i,t}|$ as fixed
168 effects, and time points as a random effect (to estimate species-to-species associations). We first plot the
169 effects of interest as scatter plots with species abundances.

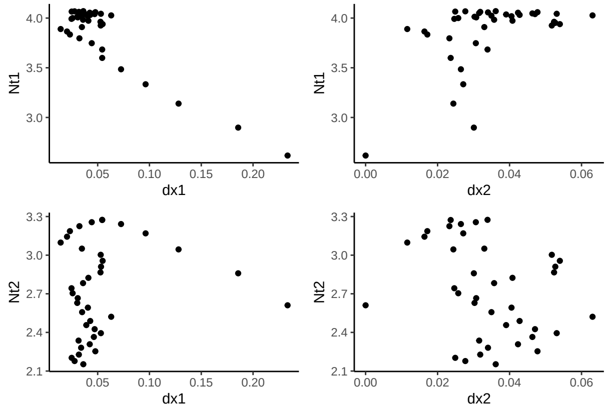


Figure 25: Population size (logarithm) in species 1 and 2 at one time step $N_{i,t+1}$ as a function of change in trait value of Species 1 and Species 2 in the previous time step $|\Delta x_{t,t-1}|$.

170 We expect that change in traits in Species 1 will have an effect on Species abundances, and we do not expect

a strong impact of change in traits in Species 2.

We now use the species abundances as the response variable in the model, with $E$, $E^2$, and $|x_{t+1} - x_{i,t}|$ as fixed effects, and time points as a random effect (to estimate species-to-species associations). We use the absolute value of the change in trait value from one time point to the next (the total amount of trait change, regardless of direction) as a fixed effect in the HMSC.

```
dat <- as.data.frame(cbind(log(N), x))
dat$time <- 1:t
df <- data.frame(dat[(2:t), -5])
colnames(df) <- c("Nt1", "Nt2", "xt1", "xt2")
df$dx1 <- abs(dat$x1[2:t] - dat$x1[1:(t - 1)])
df$dx2 <- abs(dat$x2[2:t] - dat$x2[1:(t - 1)])
Y <- as.matrix(cbind(df$Nt1, df$Nt2))
XData <- data.frame(cbind(E[1:(t - 1)], E[1:(t - 1)]^2, abs(dat$x1[2:t] - dat$x1[1:(t -
    1)]), abs(dat$x2[2:t] - dat$x2[1:(t - 1)])))
colnames(XData) <- c("E", "Esq", "dx1", "dx2")
studyDesign = data.frame(sample = as.factor(1:(t - 1)))
rL = HmscRandomLevel(units = studyDesign$sample)
m.7.hmsc = Hmsc(Y = Y, XData = XData, XFormula = ~E + Esq + dx1 + dx2, studyDesign = studyDesign,
    ranLevels = list(sample = rL))
# Bayesian model parameters
nChains <- 2
thin <- 5
samples <- 2000
transient <- 1000 * thin
verbose <- 500 * thin
# sample MCMC
m.7.sample <- sampleMcmc(m.7.hmsc, thin = thin, sample = samples, transient = transient,
    nChains = nChains, verbose = verbose)
#> Computing chain 1
#> Chain 1, iteration 2500 of 15000 (transient)
#> Chain 1, iteration 5000 of 15000 (transient)
#> Chain 1, iteration 7500 of 15000 (sampling)
#> Chain 1, iteration 10000 of 15000 (sampling)
#> Chain 1, iteration 12500 of 15000 (sampling)
#> Chain 1, iteration 15000 of 15000 (sampling)
#> Computing chain 2
#> Chain 2, iteration 2500 of 15000 (transient)
#> Chain 2, iteration 5000 of 15000 (transient)
#> Chain 2, iteration 7500 of 15000 (sampling)
#> Chain 2, iteration 10000 of 15000 (sampling)
#> Chain 2, iteration 12500 of 15000 (sampling)
#> Chain 2, iteration 15000 of 15000 (sampling)
```

```
m7.post.hmsc <- convertToCodaObject(m.7.sample)
summary(m7.post.hmsc$Beta)
#>
#> Iterations = 5005:15000
#> Thinning interval = 5
#> Number of chains = 2
#> Sample size per chain = 2000
#>
#> 1. Empirical mean and standard deviation for each variable,
```

```
#>    plus standard error of the mean:
#>
#>                                    Mean     SD  Naive SE Time-series SE
#> B[(Intercept) (C1), sp1 (S1)]  2.8689 0.1496 0.0023656      0.0023658
#> B[E (C2), sp1 (S1)]            1.2923 0.2061 0.0032581      0.0032036
#> B[Esq (C3), sp1 (S1)]         -0.3329 0.0639 0.0010104      0.0009934
#> B[dx1 (C4), sp1 (S1)]         -4.9707 0.2279 0.0036032      0.0035258
#> B[dx2 (C5), sp1 (S1)]          3.4148 0.7182 0.0113554      0.0120523
#> B[(Intercept) (C1), sp2 (S2)]  5.2820 0.1383 0.0021875      0.0022304
#> B[E (C2), sp2 (S2)]           -2.4741 0.1901 0.0030053      0.0030564
#> B[Esq (C3), sp2 (S2)]          0.4666 0.0587 0.0009282      0.0009283
#> B[dx1 (C4), sp2 (S2)]         -4.2172 0.2070 0.0032722      0.0032723
#> B[dx2 (C5), sp2 (S2)]          6.8562 0.6637 0.0104933      0.0098431
#>
#> 2. Quantiles for each variable:
#>
#>                                   2.5%     25%     50%     75%   97.5%
#> B[(Intercept) (C1), sp1 (S1)]  2.5793  2.7686  2.8651  2.9692  3.1651
#> B[E (C2), sp1 (S1)]            0.8843  1.1514  1.2964  1.4294  1.6897
#> B[Esq (C3), sp1 (S1)]         -0.4588 -0.3752 -0.3335 -0.2898 -0.2072
#> B[dx1 (C4), sp1 (S1)]         -5.4259 -5.1174 -4.9671 -4.8207 -4.5436
#> B[dx2 (C5), sp1 (S1)]          2.0188  2.9429  3.4159  3.8974  4.8244
#> B[(Intercept) (C1), sp2 (S2)]  5.0051  5.1913  5.2840  5.3748  5.5555
#> B[E (C2), sp2 (S2)]           -2.8438 -2.6006 -2.4787 -2.3492 -2.0933
#> B[Esq (C3), sp2 (S2)]          0.3496  0.4271  0.4678  0.5051  0.5806
#> B[dx1 (C4), sp2 (S2)]         -4.6360 -4.3514 -4.2158 -4.0857 -3.7977
#> B[dx2 (C5), sp2 (S2)]          5.5166  6.4150  6.8625  7.3011  8.1417
bayesplot::mcmc_trace(m7.post.hmsc$Beta)
```

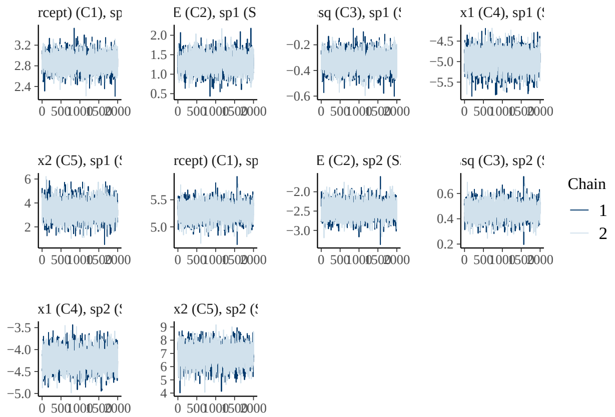


```
bayesplot::mcmc_areas(m7.post.hmsc$Beta, area_method = c("equal height"))
```

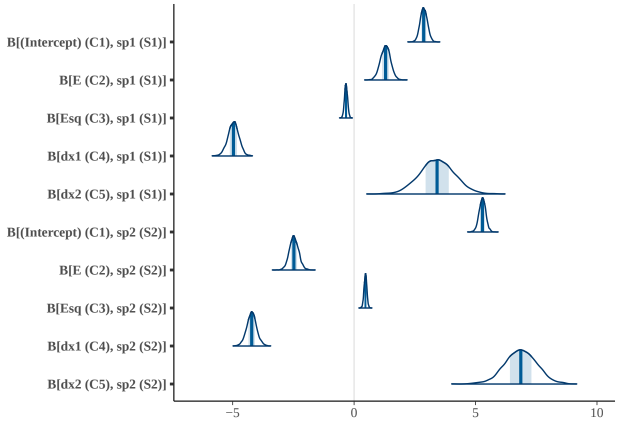


```r
# Bayesian estimates
cbind(summary(m7.post.hmsc$Beta)$statistics[1:10, 1], summary(m7.post.hmsc$Beta)$quantiles[,
    c(1, 5)])
#>                                                  2.5%      97.5%
#> B[(Intercept) (C1), sp1 (S1)]  2.8689310  2.5792716  3.1651279
#> B[E (C2), sp1 (S1)]            1.2923080  0.8842944  1.6897454
#> B[Esq (C3), sp1 (S1)]         -0.3328958 -0.4587869 -0.2072090
#> B[dx1 (C4), sp1 (S1)]         -4.9706907 -5.4258616 -4.5435503
#> B[dx2 (C5), sp1 (S1)]          3.4147944  2.0187669  4.8244125
#> B[(Intercept) (C1), sp2 (S2)]  5.2819918  5.0050821  5.5554974
#> B[E (C2), sp2 (S2)]           -2.4740977 -2.8437664 -2.0932774
#> B[Esq (C3), sp2 (S2)]          0.4665621  0.3495577  0.5805896
#> B[dx1 (C4), sp2 (S2)]         -4.2171885 -4.6359520 -3.7976972
#> B[dx2 (C5), sp2 (S2)]          6.8562285  5.5166052  8.1416526
```

```r
OmegaCor = computeAssociations(m.7.sample)
supportLevel = 0.95
toPlot = ((OmegaCor[[1]]$support > supportLevel) + (OmegaCor[[1]]$support < (1 -
    supportLevel)) > 0) * OmegaCor[[1]]$mean
corrplot::corrplot(toPlot, method = "color", col = colorRampPalette(c("blue", "white",
    "red"))(200), title = paste("random effect level:", m.7.sample$rLNames[1]), mar = c(0,
    0, 1, 0))
```

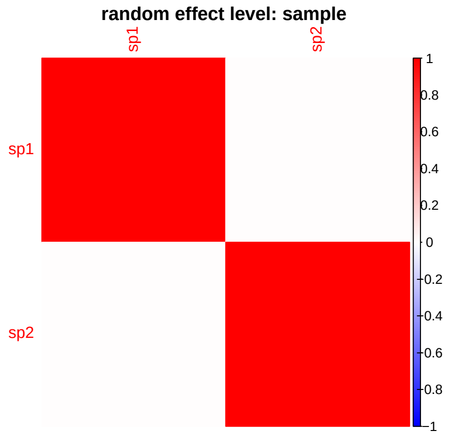

```
OmegaCor
#> [[1]]
#> [[1]]$mean
#>             sp1        sp2
#> sp1  1.0000000 -0.4103122
#> sp2 -0.4103122  1.0000000
#>
#> [[1]]$support
#>         sp1     sp2
#> sp1 1.00000 0.24775
#> sp2 0.24775 1.00000
```

```
Gradient <- constructGradient(m.7.sample, focalVariable = "E", non.focalVariables = list(Esq = list(2),
    dx1 = list(1), dx2 = list(1)), ngrid = 39)
# Esq is manually constructed as gradient-produced E^2
Gradient$XDataNew$Esq <- Gradient$XDataNew$E^2
predY <- predict(m.7.sample, XData = Gradient$XDataNew, expected = TRUE)
a <- plotGradient(m.7.sample, Gradient, pred = predY, showData = T, measure = "Y",
    index = 1, main = "", xlab = "E_t", ylab = "predicted N1_t+1")
```

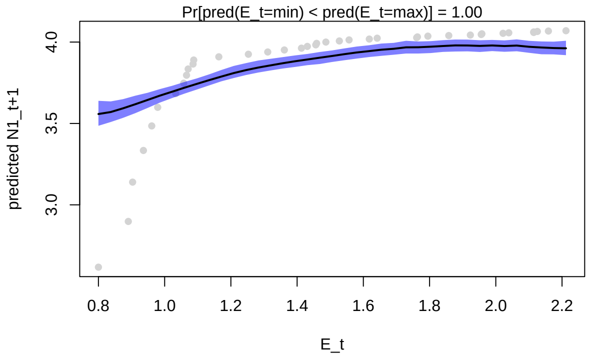


179

```
b <- plotGradient(m.7.sample, Gradient, pred = predY, showData = T, measure = "Y",
    index = 2, main = "", xlab = "E_t", ylab = "predicted N2_t+1")
```

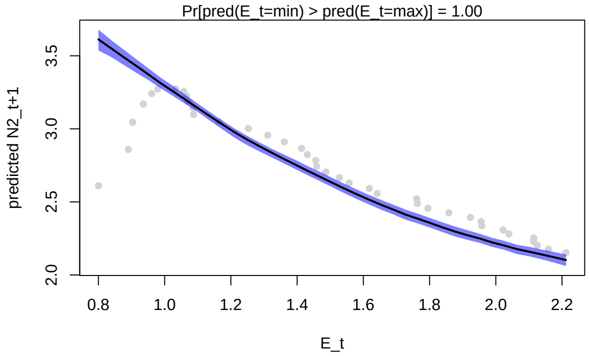


180

```
postBeta = getPostEstimate(m.7.sample, parName = "Beta")
plotBeta(m.7.sample, post = postBeta, param = "Mean", supportLevel = 0.95)
```

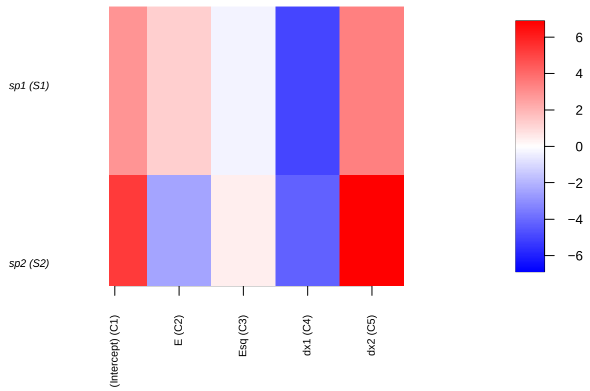


```
VP <- computeVariancePartitioning(m.7.sample, group = c(1, 1, 1, 2, 3), groupnames = c("Env",
    "Sp1", "Sp2"))
plotVariancePartitioning(m.7.sample, VP, cols = c("white", "skyblue", "darkgrey",
    "orange"), args.legend = list(cex = 0.75, bg = "transparent"))
```

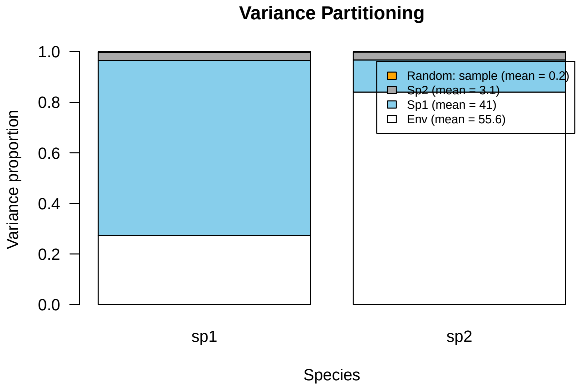


We can also look at a gradient plot for the effect of evolution in Species 1 for abundance in Species 1 and Species 2.

```
Gradient <- constructGradient(m.7.sample, focalVariable = "dx1", non.focalVariables = list(E = list(1),
    Esq = list(2), dx2 = list(1)), ngrid = 39)
Gradient$XDataNew$Esq <- Gradient$XDataNew$E^2
predY <- predict(m.7.sample, XData = Gradient$XDataNew, expected = TRUE)
a <- plotGradient(m.7.sample, Gradient, pred = predY, showData = T, measure = "Y",
    index = 1, main = "", xlab = "dx1_t", ylab = "predicted N1_t+1")
```

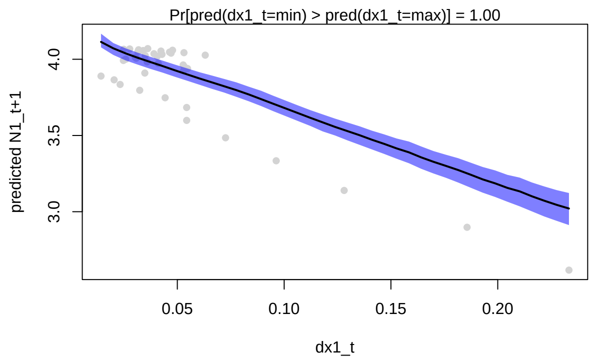


```
b <- plotGradient(m.7.sample, Gradient, pred = predY, showData = T, measure = "Y",
    index = 2, main = "", xlab = "dx1_t", ylab = "predicted N2_t+1")
```

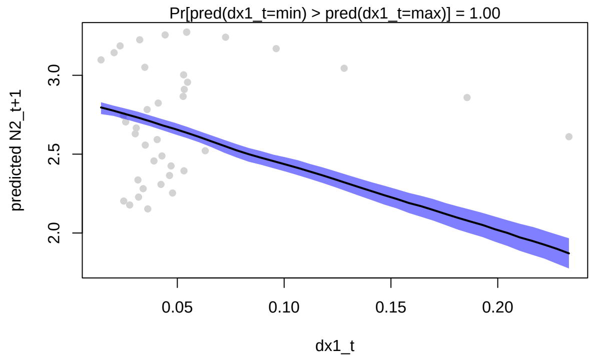


```
postBeta = getPostEstimate(m.7.sample, parName = "Beta")
plotBeta(m.7.sample, post = postBeta, param = "Mean", supportLevel = 0.95)
```

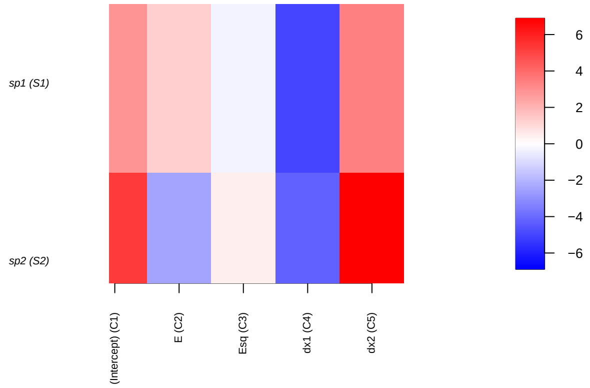


We repeat this plot for the effect of evolution in Species 2 for abundance in Species 1 and Species 2.

```
Gradient <- constructGradient(m.7.sample, focalVariable = "dx2", non.focalVariables = list(E = list(1),
    Esq = list(2), dx1 = list(1)), ngrid = 39)
Gradient$XDataNew$Esq <- Gradient$XDataNew$E^2
predY <- predict(m.7.sample, XData = Gradient$XDataNew, expected = TRUE)
a <- plotGradient(m.7.sample, Gradient, pred = predY, showData = T, measure = "Y",
    index = 1, main = "", xlab = "dx2_t", ylab = "predicted N1_t+1")
```

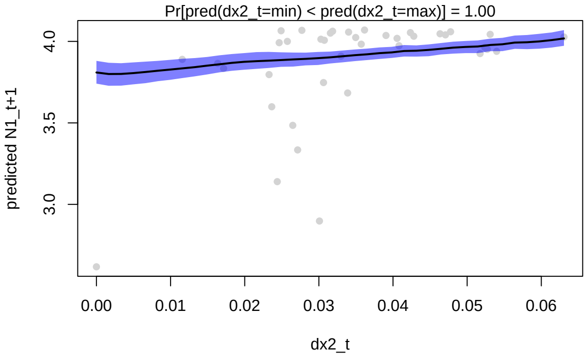


```
b <- plotGradient(m.7.sample, Gradient, pred = predY, showData = T, measure = "Y",
    index = 2, main = "", xlab = "dx2_t", ylab = "predicted N2_t+1")
```

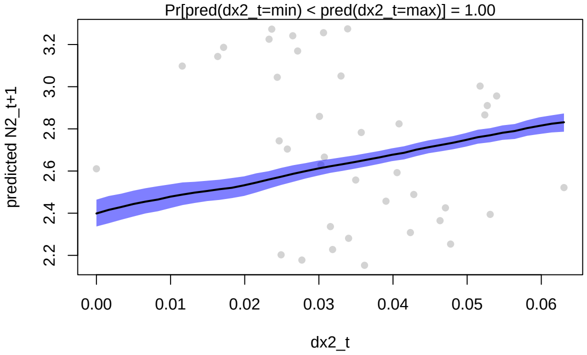


The analysis indicates that we can successfully use the HMSC analysis to partition drivers of species abundances that include evolving traits. We see that change in Species 1 trait is a strong driver of abundances in Species1 and 2, and that trait change in Species 2 has a weaker impact. This aligns well with the observed simulation results.

Though some of the main effect coefficients aren't well reflected in the data, we recall the model has strong predictive power when the elements are combined, with high predictive power to the observed data.

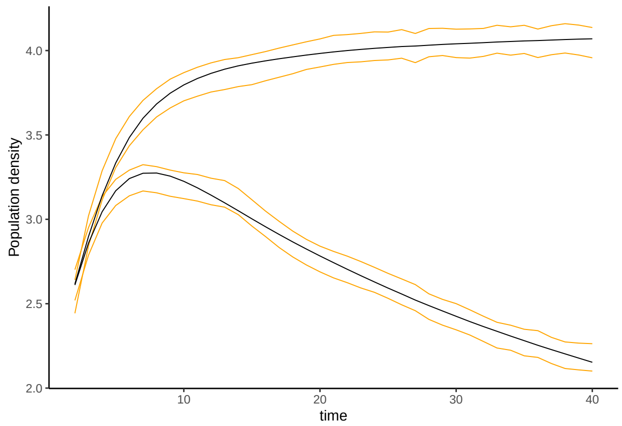


Figure 26: Observed (black) and HMSC model-predicted population size (logarithm, orange) for Species 1 and Species 2, using environment $E_t$ and change in trait value $|\Delta x|$ as a fixed effect predictor and specifying temporal random effects to capture the species interactions.

However, we can see the coefficients are less informative for inference. It is specific to this set of simulation conditions that the degree of evolution occurs at the beginning of the simulation, when the species is more mal-adapted. To better consider how trait evolution drives changes in population size, we use this change as the response variable Y in the predictive model.

```
# Plot simulation: ggplot
dat <- as.data.frame(cbind(log(N$N1), log(N$N2), x$x1, x$x2))
colnames(dat) <- c("N1", "N2", "x1", "x2")
dat$time <- 1:t
df <- data.frame(cbind(dat$N1[2:t], dat$N2[2:t], dat$x1[2:t], dat$x2[2:t]))
colnames(df) <- c("Nt1", "Nt2", "xt1", "xt2")
df$dN1 <- abs(dat$N1[2:t] - dat$N1[1:(t - 1)])
df$dN2 <- abs(dat$N2[2:t] - dat$N2[1:(t - 1)])
df$dx1 <- abs(dat$x1[2:t] - dat$x1[1:(t - 1)])
df$dx2 <- abs(dat$x2[2:t] - dat$x2[1:(t - 1)])
# prepare data in HMSC format
Y <- as.matrix(cbind(df$dN1, df$dN2))
XData <- data.frame(cbind(E[1:(t - 1)], E[1:(t - 1)]^2, abs(dat$x1[2:t] - dat$x1[1:(t -
    1)]), abs(dat$x2[2:t] - dat$x2[1:(t - 1)])))
colnames(XData) <- c("E", "Esq", "dx1", "dx2")
studyDesign = data.frame(sample = as.factor(1:(t - 1)))
rL = HmscRandomLevel(units = studyDesign$sample)
m.8.hmsc = Hmsc(Y = Y, XData = XData, XFormula = ~E + Esq + dx1 + dx2, studyDesign = studyDesign,
    ranLevels = list(sample = rL))
# Bayesian model parameters
nChains <- 2
thin <- 5
samples <- 2000
transient <- 1000 * thin
verbose <- 500 * thin
# sample MCMC
m.8.sample <- sampleMcmc(m.8.hmsc, thin = thin, sample = samples, transient = transient,
    nChains = nChains, verbose = verbose)
#> Computing chain 1
#> Chain 1, iteration 2500 of 15000 (transient)
```

```
#> Chain 1, iteration 5000 of 15000 (transient)
#> Chain 1, iteration 7500 of 15000 (sampling)
#> Chain 1, iteration 10000 of 15000 (sampling)
#> Chain 1, iteration 12500 of 15000 (sampling)
#> Chain 1, iteration 15000 of 15000 (sampling)
#> Computing chain 2
#> Chain 2, iteration 2500 of 15000 (transient)
#> Chain 2, iteration 5000 of 15000 (transient)
#> Chain 2, iteration 7500 of 15000 (sampling)
#> Chain 2, iteration 10000 of 15000 (sampling)
#> Chain 2, iteration 12500 of 15000 (sampling)
#> Chain 2, iteration 15000 of 15000 (sampling)
```

```
m8.post.hmsc <- convertToCodaObject(m.8.sample)
summary(m8.post.hmsc$Beta)
#>
#> Iterations = 5005:15000
#> Thinning interval = 5
#> Number of chains = 2
#> Sample size per chain = 2000
#>
#> 1. Empirical mean and standard deviation for each variable,
#>    plus standard error of the mean:
#>
#>                                    Mean      SD  Naive SE Time-series SE
#> B[(Intercept) (C1), sp1 (S1)]  0.34742 0.11089 0.0017533      0.0017037
#> B[E (C2), sp1 (S1)]           -0.41415 0.15298 0.0024189      0.0024189
#> B[Esq (C3), sp1 (S1)]          0.11514 0.04751 0.0007512      0.0007512
#> B[dx1 (C4), sp1 (S1)]          1.15744 0.17136 0.0027095      0.0027600
#> B[dx2 (C5), sp1 (S1)]         -0.63291 0.53307 0.0084285      0.0086952
#> B[(Intercept) (C1), sp2 (S2)] -0.10279 0.10751 0.0016999      0.0017953
#> B[E (C2), sp2 (S2)]            0.17311 0.14811 0.0023418      0.0024753
#> B[Esq (C3), sp2 (S2)]         -0.05427 0.04584 0.0007247      0.0007349
#> B[dx1 (C4), sp2 (S2)]          1.39762 0.16062 0.0025396      0.0025399
#> B[dx2 (C5), sp2 (S2)]         -1.28696 0.51197 0.0080949      0.0082622
#>
#> 2. Quantiles for each variable:
#>
#>                                   2.5%      25%      50%      75%    97.5%
#> B[(Intercept) (C1), sp1 (S1)]  0.12485  0.27474  0.34921  0.42026  0.56453
#> B[E (C2), sp1 (S1)]           -0.72011 -0.51493 -0.41532 -0.31373 -0.10810
#> B[Esq (C3), sp1 (S1)]          0.02018  0.08448  0.11563  0.14650  0.20978
#> B[dx1 (C4), sp1 (S1)]          0.82494  1.04445  1.15561  1.27024  1.49420
#> B[dx2 (C5), sp1 (S1)]         -1.67933 -0.99984 -0.62392 -0.27481  0.43213
#> B[(Intercept) (C1), sp2 (S2)] -0.32034 -0.17342 -0.10219 -0.03317  0.11076
#> B[E (C2), sp2 (S2)]           -0.11713  0.07718  0.17172  0.26905  0.47630
#> B[Esq (C3), sp2 (S2)]         -0.14704 -0.08439 -0.05379 -0.02421  0.03474
#> B[dx1 (C4), sp2 (S2)]          1.06702  1.29512  1.39933  1.50135  1.70843
#> B[dx2 (C5), sp2 (S2)]         -2.28991 -1.63008 -1.28566 -0.94407 -0.27855
bayesplot::mcmc_trace(m8.post.hmsc$Beta)
```

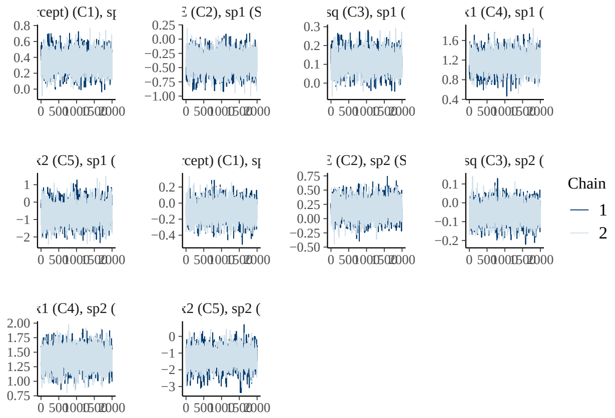


```r
bayesplot::mcmc_areas(m8.post.hmsc$Beta, area_method = c("equal height"))
```

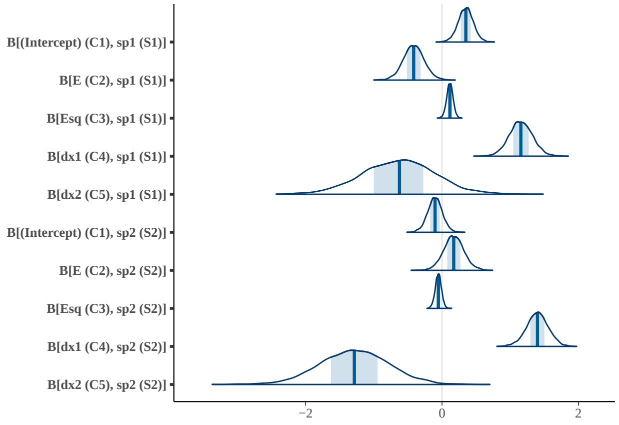


```r
Gradient <- constructGradient(m.8.sample, focalVariable = "dx1", non.focalVariables = list(E = list(1),
    Esq = list(2), dx2 = list(1)), ngrid = 39)
Gradient$XDataNew$Esq <- Gradient$XDataNew$E^2
predY <- predict(m.8.sample, XData = Gradient$XDataNew, expected = TRUE)
a <- plotGradient(m.8.sample, Gradient, pred = predY, showData = T, measure = "Y",
    index = 1, main = "", xlab = "dx1_t", ylab = "predicted dN1_t+1")
```

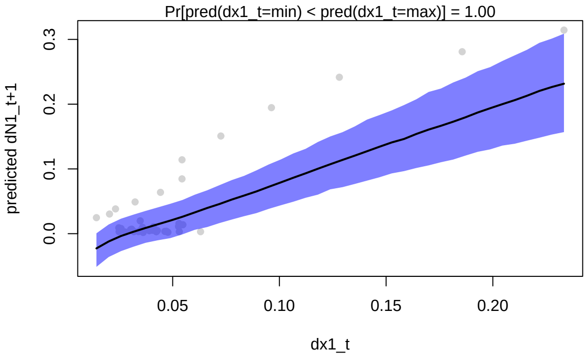

```
b <- plotGradient(m.8.sample, Gradient, pred = predY, showData = T, measure = "Y",
    index = 2, main = "", xlab = "dx1_t", ylab = "predicted dN2_t+1")
```

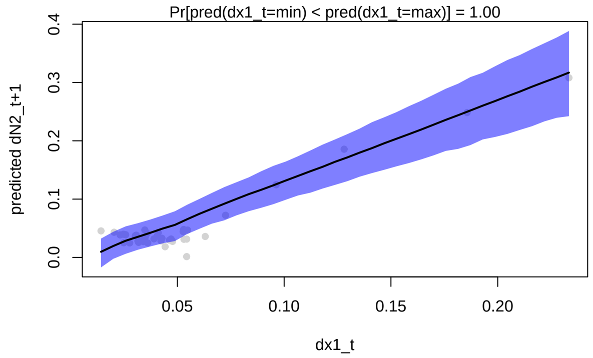


```
postBeta = getPostEstimate(m.8.sample, parName = "Beta")
plotBeta(m.8.sample, post = postBeta, param = "Mean", supportLevel = 0.95)
```

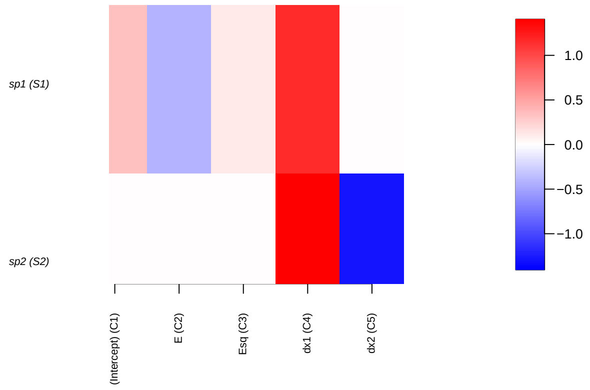


We repeat this plot for the effect of evolution in Species 2 for abundance in Species 1 and Species 2.

```
Gradient <- constructGradient(m.8.sample, focalVariable = "dx2", non.focalVariables = list(E = list(1),
    Esq = list(2), dx1 = list(1)), ngrid = 39)
Gradient$XDataNew$Esq <- Gradient$XDataNew$E^2
predY <- predict(m.8.sample, XData = Gradient$XDataNew, expected = TRUE)
a <- plotGradient(m.8.sample, Gradient, pred = predY, showData = T, measure = "Y",
    index = 1, main = "", xlab = "dx2_t", ylab = "predicted dN1_t+1")
```

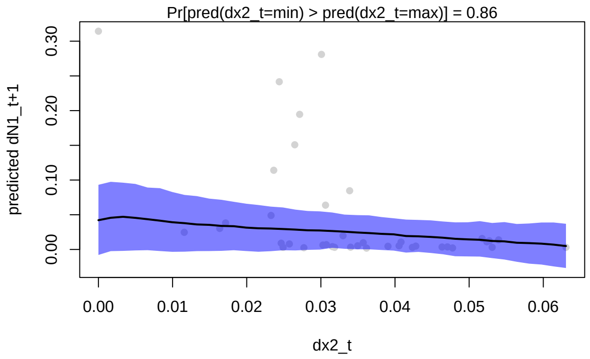


```r
b <- plotGradient(m.8.sample, Gradient, pred = predY, showData = T, measure = "Y",
    index = 2, main = "", xlab = "dx2_t", ylab = "predicted dN2_t+1")
```

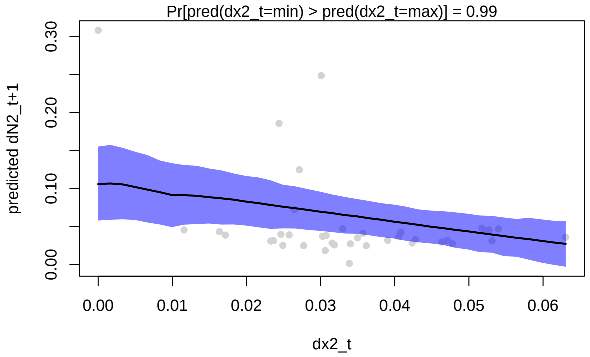


```r
VP <- computeVariancePartitioning(m.8.sample, group = c(1, 1, 1, 2, 3), groupnames = c("Env",
    "Sp1", "Sp2"))
plotVariancePartitioning(m.8.sample, VP, cols = c("white", "skyblue", "darkgrey",
    "orange"), args.legend = list(cex = 0.75, bg = "transparent"))
```

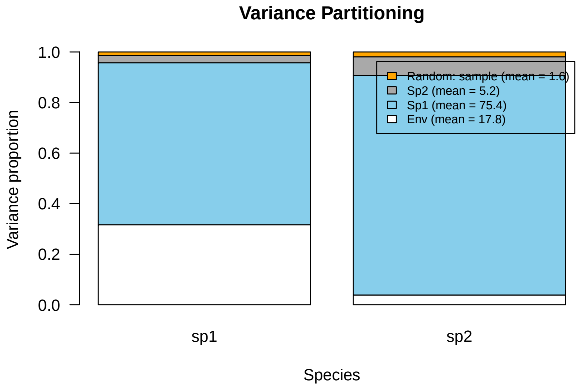


Here we can see that trait evolution in Species 1 had a strong positive relationship with the *change* in population size for both species. Trait evolution in Species 2 had much less of an effect.

### 6 Model F: Species growth, competition, environmental change, and trait evolution in a spatially structured environment

We now consider that trait evolution can occur, with an impact on the species growth rate / fitness in the context of the environment. We also add a final consideration to the model, the existence of multiple sites with correlation structure in the variance of species abundances. We consider this in our simulation by adding multiple (10) sites, with low vs. high spatial covariance in abundances, depending on distance between sites. We then include spatial random effects in the HMSC statistical model, where species associations are considered as a function of distance between sites.

#### 6.1 Population dynamics simulation

We use the same example as above, but now with 10 sites. To illustrate how HMSC can estimate the impacts of distance-dependent spatial covariance, we duplicate the population time series obtained in the example above, but we add in site variance and among-site covariance.

```
# Case 1: Weaker interactions, stronger environment Simulate initial species
# population growth with environment fluctuations
N0 <- c(10, 10)
E.0 <- 0.8
x.0 <- c(0.1, 0.8)
alpha.11 <- 0.01
alpha.22 <- 0.01
alpha.12 <- 0.005
alpha.21 <- 0.01
alpha <- matrix(c(alpha.11, alpha.21, alpha.12, alpha.22), nrow = 2, byrow = FALSE)
P <- 1
w <- 2
Wmax <- 2
h2 <- 1
k <- (w + (1 - h2) * P)/(P + w)
j <- 10
# Simulation of model for t time steps, i sites
xycoords = matrix(runif(2 * j), ncol = 2)
rownames(xycoords) <- 1:j
t <- 40
N <- array(NA, dim = c(j * t, length(N0)))
N <- as.data.frame(N)
colnames(N) <- paste0("N", 1:length(N0))
x <- array(NA, dim = c(j * t, length(N0)))
x <- as.data.frame(x)
colnames(x) <- paste0("x", 1:length(N0))
r <- array(NA, dim = c(j * t, length(N0)))
r <- as.data.frame(r)
colnames(r) <- paste0("r", 1:length(N0))
N[1:j, ] <- rep(N0, each = j)
E <- rep(NA, t)
E[1] <- E.0
x[1:j, ] <- rep(x.0, each = j)
What <- Wmax * sqrt(w/(P + w))
r0 <- as.numeric(What * exp((-(((w + (1 - h2) * P)/(P + w)) * (E[1] - x[1, ]))^2)/(2 *
    (P + w))))
```

```
r[1:j, ] <- rep(r0, each = j)
# Keep track of site and time
study <- array(NA, dim = c(j * t, 3))
study <- as.data.frame(study)
colnames(study) <- c("obs", "site", "time")
study$obs <- 1:dim(study)[1]
study$site <- rep(1:j, times = t)
study$time <- rep(1:t, each = j)

for (i in 2:t) {
    for (z in 1:j) {
        res <- disc_LV_evol(N0 = N[study$site == z & study$time == i - 1, ], alpha = alpha,
            E = E[i - 1], x = x[study$site == z & study$time == i - 1, ], P = P,
            w = w, Wmax = Wmax, h2 = h2)
        N[study$site == z & study$time == i, ] <- res$Nt1
        r[study$site == z & study$time == i, ] <- res$r
        # trait change
        d <- E[i - 1] - x[study$site == z & study$time == i - 1, ]
        d1 <- k * d
        x[study$site == z & study$time == i, ] <- E[i - 1] - d1
    }
    # environmental change
    E[i] <- E[i - 1] + abs(rnorm(1, 0, 0.05))
}
# site-level random effect
sigma <- 0
sigma.spatial <- 2
alpha.spatial <- 0.5
Sigma = sigma.spatial^2 * exp(-as.matrix(dist(xycoords))/alpha.spatial)
# draw from covariance matrix
a = mvrnorm(mu = rep(0, j), Sigma = Sigma)
for (i in 1:t) {
    N$N1[study$time == i] <- N$N1[study$time == i] + a
}
a = mvrnorm(mu = rep(0, j), Sigma = Sigma)
for (i in 1:t) {
    N$N2[study$time == i] <- N$N2[study$time == i] + a
}
```

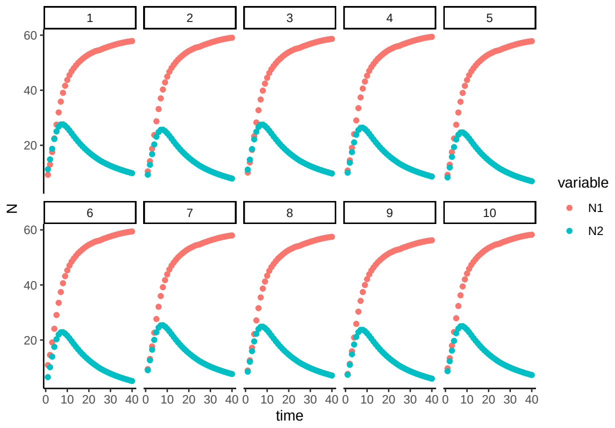


Figure 27: Population size $N$ over time $t$ for a discrete-time logistic growth model with competition, a changing environment, and trait evolution. Values are shown at 10 sites, with site covariance in species abundances depending on spatial distance.

223 We now fit a spatial random effect, and add that to the HMSC model.

```
# prepare data in HMSC format
dat <- as.data.frame(cbind(log(N), x))
dat$time <- study$time
df <- data.frame(cbind(dat$N1[dat$time != 1], dat$N2[dat$time != 1], dat$x1[dat$time !=
    1], dat$x2[dat$time != 1]))
colnames(df) <- c("Nt1", "Nt2", "xt1", "xt2")
Y <- as.matrix(cbind(df$Nt1, df$Nt2))
XData <- data.frame(E = rep(E[1:(t - 1)], each = j))
XData$Esq <- XData$E^2
XData$dx1 <- abs(dat$x1[dat$time != 1] - dat$x1[dat$time != t])
XData$dx2 <- abs(dat$x2[dat$time != 1] - dat$x2[dat$time != t])
# Update study design to include spatial random effects
samp1 <- 1:(j * t)
samp1 <- as.data.frame(samp1)
samp1 <- subset(samp1, samp1 > 10)
rownames(XData) <- row.names(samp1)
studyDesign = data.frame(sample = 1:(j * (t - 1)), site = study$site[study$time >
    1])
studyDesign$sample <- as.factor(studyDesign$sample)
studyDesign$site <- as.factor(studyDesign$site)
rL.spatial = HmscRandomLevel(sData = xycoords)
rL.sample = HmscRandomLevel(units = studyDesign$sample)
# Fit HMSC model
m.9.hmsc = Hmsc(Y = Y, XData = XData, XFormula = ~E + Esq + dx1 + dx2, studyDesign = studyDesign,
    ranLevels = list(sample = rL.sample, site = rL.spatial))
# Bayesian model parameters
nChains <- 2
thin <- 5
samples <- 2000
transient <- 1000 * thin
verbose <- 500 * thin
# sample MCMC
m.9.sample <- sampleMcmc(m.9.hmsc, thin = thin, sample = samples, transient = transient,
```

```
    nChains = nChains, verbose = verbose)
#> Computing chain 1
#> Chain 1, iteration 2500 of 15000 (transient)
#> Chain 1, iteration 5000 of 15000 (transient)
#> Chain 1, iteration 7500 of 15000 (sampling)
#> Chain 1, iteration 10000 of 15000 (sampling)
#> Chain 1, iteration 12500 of 15000 (sampling)
#> Chain 1, iteration 15000 of 15000 (sampling)
#> Computing chain 2
#> Chain 2, iteration 2500 of 15000 (transient)
#> Chain 2, iteration 5000 of 15000 (transient)
#> Chain 2, iteration 7500 of 15000 (sampling)
#> Chain 2, iteration 10000 of 15000 (sampling)
#> Chain 2, iteration 12500 of 15000 (sampling)
#> Chain 2, iteration 15000 of 15000 (sampling)
```

```
m9.post.hmsc <- convertToCodaObject(m.9.sample)
summary(m9.post.hmsc$Beta)
#>
#> Iterations = 5005:15000
#> Thinning interval = 5
#> Number of chains = 2
#> Sample size per chain = 2000
#>
#> 1. Empirical mean and standard deviation for each variable,
#>    plus standard error of the mean:
#>
#>                                    Mean       SD  Naive SE Time-series SE
#> B[(Intercept) (C1), sp1 (S1)]  2.9084 0.032736 0.0005176      0.0005175
#> B[E (C2), sp1 (S1)]            1.1219 0.029259 0.0004626      0.0004626
#> B[Esq (C3), sp1 (S1)]         -0.2641 0.008319 0.0001315      0.0001315
#> B[dx1 (C4), sp1 (S1)]         -4.6592 0.047055 0.0007440      0.0007827
#> B[dx2 (C5), sp1 (S1)]          3.6255 0.093157 0.0014729      0.0014727
#> B[(Intercept) (C1), sp2 (S2)]  4.6626 0.139811 0.0022106      0.0021717
#> B[E (C2), sp2 (S2)]           -1.5738 0.068967 0.0010905      0.0010906
#> B[Esq (C3), sp2 (S2)]          0.1966 0.019616 0.0003102      0.0003102
#> B[dx1 (C4), sp2 (S2)]         -4.4286 0.111548 0.0017637      0.0018169
#> B[dx2 (C5), sp2 (S2)]          6.9817 0.222005 0.0035102      0.0035103
#>
#> 2. Quantiles for each variable:
#>
#>                                   2.5%     25%     50%     75%   97.5%
#> B[(Intercept) (C1), sp1 (S1)]  2.8459  2.8873  2.9080  2.9285  2.9739
#> B[E (C2), sp1 (S1)]            1.0640  1.1021  1.1215  1.1419  1.1808
#> B[Esq (C3), sp1 (S1)]         -0.2809 -0.2697 -0.2640 -0.2584 -0.2476
#> B[dx1 (C4), sp1 (S1)]         -4.7500 -4.6908 -4.6599 -4.6272 -4.5679
#> B[dx2 (C5), sp1 (S1)]          3.4420  3.5626  3.6249  3.6892  3.8084
#> B[(Intercept) (C1), sp2 (S2)]  4.3755  4.5838  4.6604  4.7432  4.9433
#> B[E (C2), sp2 (S2)]           -1.7094 -1.6195 -1.5742 -1.5265 -1.4392
#> B[Esq (C3), sp2 (S2)]          0.1586  0.1830  0.1966  0.2098  0.2356
#> B[dx1 (C4), sp2 (S2)]         -4.6469 -4.5039 -4.4287 -4.3533 -4.2125
#> B[dx2 (C5), sp2 (S2)]          6.5435  6.8324  6.9859  7.1306  7.4120
bayesplot::mcmc_trace(m9.post.hmsc$Beta)
```

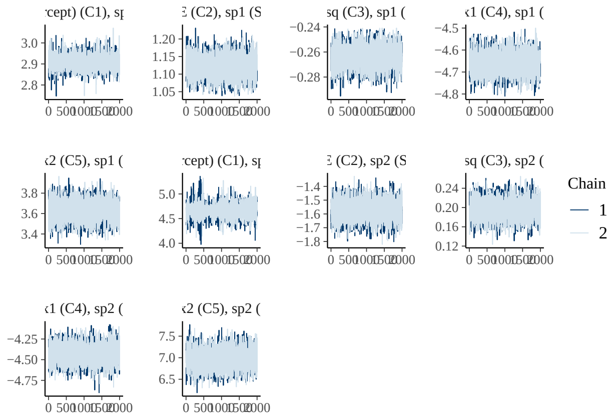


```r
bayesplot::mcmc_areas(m9.post.hmsc$Beta, area_method = c("equal height"))
```

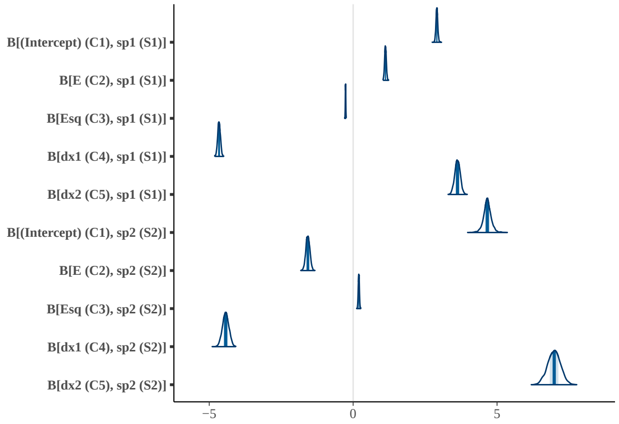


```r
VP <- computeVariancePartitioning(m.9.sample, group = c(1, 1, 1, 2, 3), groupnames = c("Env",
    "Sp1", "Sp2"))
plotVariancePartitioning(m.9.sample, VP, cols = c("white", "skyblue", "darkgrey",
    "orange", "pink"), args.legend = list(cex = 0.75, bg = "transparent"))
```

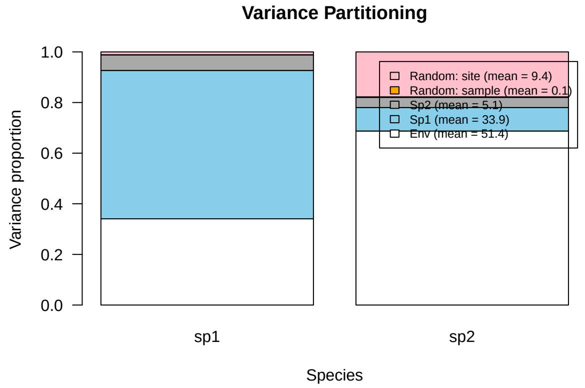


```r
OmegaCor = computeAssociations(m.9.sample)
supportLevel = 0.95
```

```
toPlot = ((OmegaCor[[1]]$support > supportLevel) + (OmegaCor[[1]]$support < (1 -
    supportLevel)) > 0) * OmegaCor[[1]]$mean
corrplot::corrplot(toPlot, method = "color", col = colorRampPalette(c("blue", "white",
    "red"))(200), title = paste("random effect level:", m.9.sample$rLNames[1]), mar = c(0,
    0, 1, 0))
```

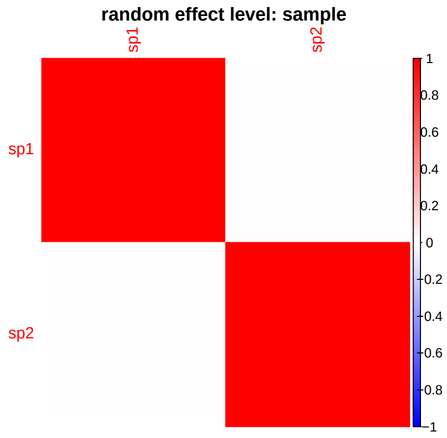


```
OmegaCor
#> [[1]]
#> [[1]]$mean
#>            sp1        sp2
#> sp1 1.00000000 0.02623117
#> sp2 0.02623117 1.00000000
#>
#> [[1]]$support
#>         sp1     sp2
#> sp1 1.00000 0.52125
#> sp2 0.52125 1.00000
#>
#>
#> [[2]]
#> [[2]]$mean
#>           sp1       sp2
#> sp1 1.0000000 0.1357261
#> sp2 0.1357261 1.0000000
#>
#> [[2]]$support
#>         sp1     sp2
#> sp1 1.00000 0.63175
#> sp2 0.63175 1.00000
toPlot = ((OmegaCor[[2]]$support > supportLevel) + (OmegaCor[[2]]$support < (1 -
    supportLevel)) > 0) * OmegaCor[[2]]$mean
corrplot::corrplot(toPlot, method = "color", col = colorRampPalette(c("blue", "white",
    "red"))(200), title = paste("random effect level:", m.9.sample$rLNames[2]), mar = c(0,
    0, 1, 0))
```

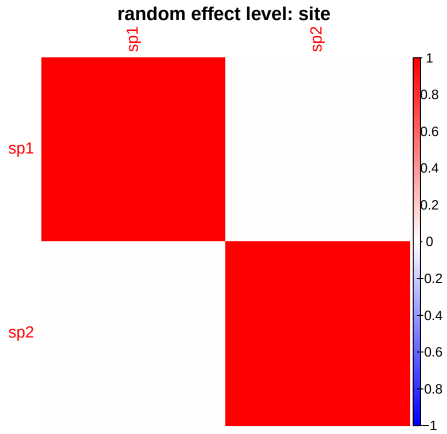


The HMSC model correctly considers spatial random effects, but they are not strong in this example. We evaluate the model's ability to detect increased spatial signal by increasing the spatial covariance in species abundances.

```
# Simulation of model for t time steps, i sites
N0 <- c(10, 10)
E.0 <- 0.8
x.0 <- c(0.1, 0.8)
alpha.11 <- 0.01
alpha.22 <- 0.01
alpha.12 <- 0.005
alpha.21 <- 0.01
alpha <- matrix(c(alpha.11, alpha.21, alpha.12, alpha.22), nrow = 2, byrow = FALSE)  # careful to make
P <- 1
w <- 2
Wmax <- 2
h2 <- 1
k <- (w + (1 - h2) * P)/(P + w)
j <- 10
# random site locations
xycoords = matrix(runif(2 * j), ncol = 2)
rownames(xycoords) <- 1:j
t <- 40
N <- array(NA, dim = c(j * t, length(N0)))
N <- as.data.frame(N)
colnames(N) <- paste0("N", 1:length(N0))
x <- array(NA, dim = c(j * t, length(N0)))
x <- as.data.frame(x)
colnames(x) <- paste0("x", 1:length(N0))
r <- array(NA, dim = c(j * t, length(N0)))
r <- as.data.frame(r)
colnames(r) <- paste0("r", 1:length(N0))
N[1:j, ] <- rep(N0, each = j)
E <- rep(NA, t)
E[1] <- E.0
x[1:j, ] <- rep(x.0, each = j)
What <- Wmax * sqrt(w/(P + w))
r0 <- as.numeric(What * exp((-(((w + (1 - h2) * P)/(P + w)) * (E[1] - x[1, ]))^2)/(2 *
    (P + w))))
r[1:j, ] <- rep(r0, each = j)
# Keep track of site and time
```

```
study <- array(NA, dim = c(j * t, 3))
study <- as.data.frame(study)
colnames(study) <- c("obs", "site", "time")
study$obs <- 1:dim(study)[1]
study$site <- rep(1:j, times = t)
study$time <- rep(1:t, each = j)

for (i in 2:t) {
    for (z in 1:j) {
        res <- disc_LV_evol(N0 = N[study$site == z & study$time == i - 1, ], alpha = alpha,
            E = E[i - 1], x = x[study$site == z & study$time == i - 1, ], P = P,
            w = w, Wmax = Wmax, h2 = h2)
        N[study$site == z & study$time == i, ] <- res$Nt1
        r[study$site == z & study$time == i, ] <- res$r
        # trait change
        d <- E[i - 1] - x[study$site == z & study$time == i - 1, ]
        d1 <- k * d
        x[study$site == z & study$time == i, ] <- E[i - 1] - d1
    }
    # environmental change
    E[i] <- E[i - 1] + abs(rnorm(1, 0, 0.05))
}
# site-level random effect
sigma <- 0
sigma.spatial <- 10
alpha.spatial <- 0.5
Sigma = sigma.spatial^2 * exp(-as.matrix(dist(xycoords))/alpha.spatial)
# draw from covariance matrix
a = mvrnorm(mu = rep(0, j), Sigma = Sigma)
for (i in 1:t) {
    N$N1[study$time == i] <- N$N1[study$time == i] + a
}
a = mvrnorm(mu = rep(0, j), Sigma = Sigma)
for (i in 1:t) {
    N$N2[study$time == i] <- N$N2[study$time == i] + a
}
```

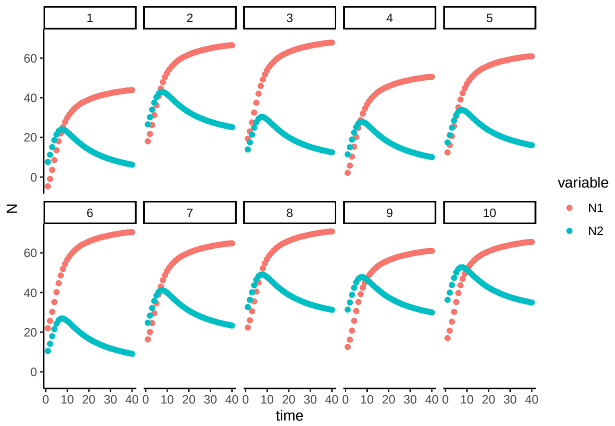


Figure 28: Population size $N$ over time $t$ for a discrete-time logistic growth model with competition, a changing environment, and trait evolution. Values are shown at 10 sites, with an increased site covariance in species abundances, depending on spatial distance.

```
# prepare data in HMSC format
dat <- as.data.frame(cbind(log(N), x))
#> Warning in FUN(X[[i]], ...): NaNs produced
dat$time <- study$time
df <- data.frame(cbind(dat$N1[dat$time != 1], dat$N2[dat$time != 1], dat$x1[dat$time !=
    1], dat$x2[dat$time != 1]))
colnames(df) <- c("Nt1", "Nt2", "xt1", "xt2")
Y <- as.matrix(cbind(df$Nt1, df$Nt2))
XData <- data.frame(E = rep(E[1:(t - 1)], each = j))
XData$Esq <- XData$E^2
XData$dx1 <- abs(dat$x1[dat$time != 1] - dat$x1[dat$time != t])
XData$dx2 <- abs(dat$x2[dat$time != 1] - dat$x2[dat$time != t])
# Update study design to include spatial random effects
samp1 <- 1:(j * t)
samp1 <- as.data.frame(samp1)
samp1 <- subset(samp1, samp1 > 10)
rownames(XData) <- row.names(samp1)
studyDesign = data.frame(sample = 1:(j * (t - 1)), site = study$site[study$time >
    1])
studyDesign$sample <- as.factor(studyDesign$sample)
studyDesign$site <- as.factor(studyDesign$site)
rL.spatial = HmscRandomLevel(sData = xycoords)
rL.sample = HmscRandomLevel(units = studyDesign$sample)
# Fit HMSC model
m.10.hmsc = Hmsc(Y = Y, XData = XData, XFormula = ~E + Esq + dx1 + dx2, studyDesign = studyDesign,
    ranLevels = list(sample = rL.sample, site = rL.spatial))
# Bayesian model parameters
nChains <- 2
thin <- 5
samples <- 2000
transient <- 1000 * thin
verbose <- 500 * thin
# sample MCMC
m.10.sample <- sampleMcmc(m.10.hmsc, thin = thin, sample = samples, transient = transient,
    nChains = nChains, verbose = verbose)
```

```
#> Computing chain 1
#> Chain 1, iteration 2500 of 15000 (transient)
#> Chain 1, iteration 5000 of 15000 (transient)
#> Chain 1, iteration 7500 of 15000 (sampling)
#> Chain 1, iteration 10000 of 15000 (sampling)
#> Chain 1, iteration 12500 of 15000 (sampling)
#> Chain 1, iteration 15000 of 15000 (sampling)
#> Computing chain 2
#> Chain 2, iteration 2500 of 15000 (transient)
#> Chain 2, iteration 5000 of 15000 (transient)
#> Chain 2, iteration 7500 of 15000 (sampling)
#> Chain 2, iteration 10000 of 15000 (sampling)
#> Chain 2, iteration 12500 of 15000 (sampling)
#> Chain 2, iteration 15000 of 15000 (sampling)
```

```
m10.post.hmsc <- convertToCodaObject(m.10.sample)
summary(m10.post.hmsc$Beta)
#>
#> Iterations = 5005:15000
#> Thinning interval = 5
#> Number of chains = 2
#> Sample size per chain = 2000
#>
#> 1. Empirical mean and standard deviation for each variable,
#>    plus standard error of the mean:
#>
#>                                   Mean      SD  Naive SE Time-series SE
#> B[(Intercept) (C1), sp1 (S1)]  2.6103 0.22509 0.0035589      0.0069848
#> B[E (C2), sp1 (S1)]            1.5400 0.19704 0.0031155      0.0030612
#> B[Esq (C3), sp1 (S1)]         -0.4194 0.06505 0.0010286      0.0010086
#> B[dx1 (C4), sp1 (S1)]         -4.1819 0.23232 0.0036734      0.0038031
#> B[dx2 (C5), sp1 (S1)]          3.5610 0.57421 0.0090790      0.0090793
#> B[(Intercept) (C1), sp2 (S2)]  4.7049 0.39613 0.0062633      0.0105717
#> B[E (C2), sp2 (S2)]           -1.7966 0.17965 0.0028405      0.0028796
#> B[Esq (C3), sp2 (S2)]          0.3351 0.05908 0.0009341      0.0009483
#> B[dx1 (C4), sp2 (S2)]         -3.1127 0.20976 0.0033166      0.0033170
#> B[dx2 (C5), sp2 (S2)]          4.9941 0.51922 0.0082096      0.0082072
#>
#> 2. Quantiles for each variable:
#>
#>                                   2.5%     25%     50%     75%   97.5%
#> B[(Intercept) (C1), sp1 (S1)]  2.1501  2.4738  2.6196  2.7615  3.0171
#> B[E (C2), sp1 (S1)]            1.1496  1.4080  1.5447  1.6725  1.9187
#> B[Esq (C3), sp1 (S1)]         -0.5450 -0.4631 -0.4212 -0.3753 -0.2923
#> B[dx1 (C4), sp1 (S1)]         -4.6397 -4.3383 -4.1818 -4.0228 -3.7336
#> B[dx2 (C5), sp1 (S1)]          2.4726  3.1741  3.5585  3.9446  4.7081
#> B[(Intercept) (C1), sp2 (S2)]  3.8424  4.4525  4.7484  4.9852  5.3821
#> B[E (C2), sp2 (S2)]           -2.1491 -1.9120 -1.7976 -1.6794 -1.4455
#> B[Esq (C3), sp2 (S2)]          0.2183  0.2966  0.3354  0.3731  0.4494
#> B[dx1 (C4), sp2 (S2)]         -3.5241 -3.2549 -3.1125 -2.9741 -2.6955
#> B[dx2 (C5), sp2 (S2)]          3.9950  4.6326  4.9966  5.3519  6.0137
bayesplot::mcmc_trace(m10.post.hmsc$Beta)
```

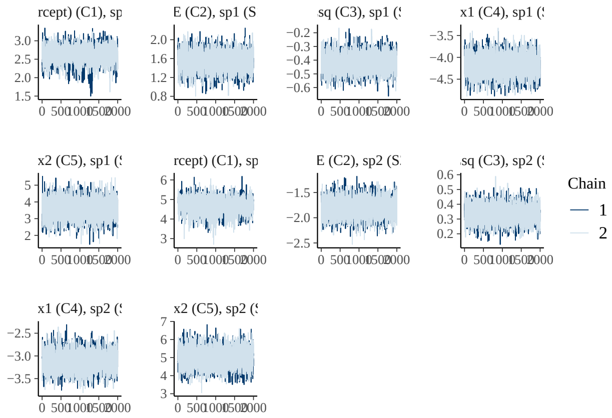


```r
bayesplot::mcmc_areas(m10.post.hmsc$Beta, area_method = c("equal height"))
```

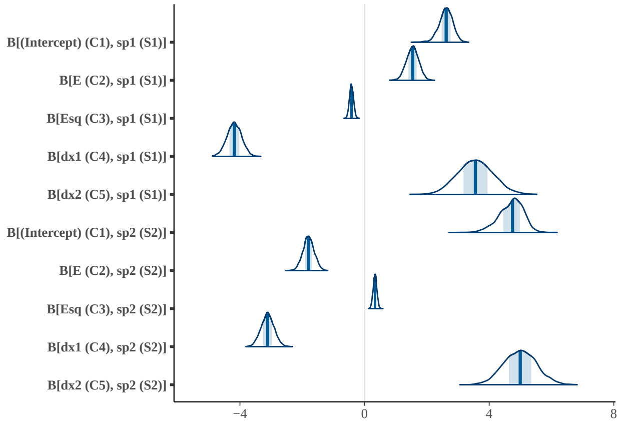


```r
VP <- computeVariancePartitioning(m.10.sample, group = c(1, 1, 1, 2, 3), groupnames = c("Env",
    "Sp1", "Sp2"))
plotVariancePartitioning(m.10.sample, VP, cols = c("white", "skyblue", "darkgrey",
    "orange", "pink"), args.legend = list(cex = 0.75, bg = "transparent"))
```

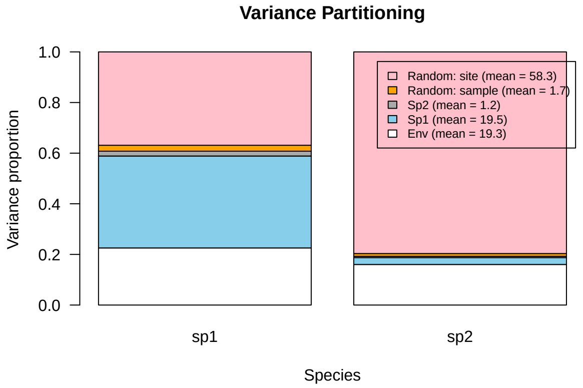


```r
OmegaCor = computeAssociations(m.9.sample)
supportLevel = 0.95
```

```
toPlot = ((OmegaCor[[1]]$support > supportLevel) + (OmegaCor[[1]]$support < (1 -
    supportLevel)) > 0) * OmegaCor[[1]]$mean
corrplot::corrplot(toPlot, method = "color", col = colorRampPalette(c("blue", "white",
    "red"))(200), title = paste("random effect level:", m.9.sample$rLNames[1]), mar = c(0,
    0, 1, 0))
```

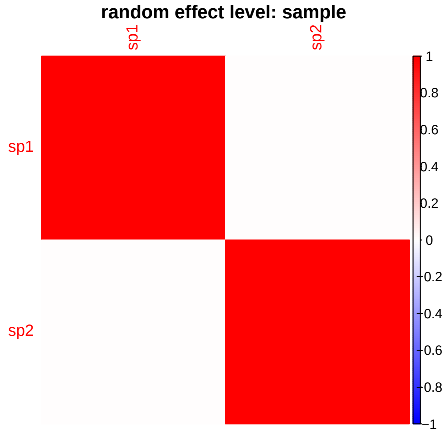


```
OmegaCor
#> [[1]]
#> [[1]]$mean
#>            sp1        sp2
#> sp1 1.00000000 0.02623117
#> sp2 0.02623117 1.00000000
#>
#> [[1]]$support
#>         sp1     sp2
#> sp1 1.00000 0.52125
#> sp2 0.52125 1.00000
#>
#>
#> [[2]]
#> [[2]]$mean
#>           sp1       sp2
#> sp1 1.0000000 0.1357261
#> sp2 0.1357261 1.0000000
#>
#> [[2]]$support
#>         sp1     sp2
#> sp1 1.00000 0.63175
#> sp2 0.63175 1.00000
toPlot = ((OmegaCor[[2]]$support > supportLevel) + (OmegaCor[[2]]$support < (1 -
    supportLevel)) > 0) * OmegaCor[[2]]$mean
corrplot::corrplot(toPlot, method = "color", col = colorRampPalette(c("blue", "white",
    "red"))(200), title = paste("random effect level:", m.9.sample$rLNames[2]), mar = c(0,
    0, 1, 0))
```

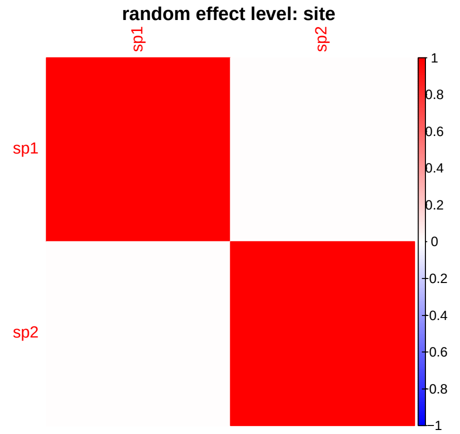


We observe a much greater fraction of variation explained by the spatial random effects.

### References

Beverton, R. J., and S. J. Holt. 1957. On the dynamics of exploited fish populations (Vol. 11). Springer Science & Business Media.

Certain, G., F. Barraquand, and A. Gårdmark. 2018. How do MAR(1) models cope with hidden nonlinearities in ecological dynamics? Methods in Ecology and Evolution 9:1975–1995.

Erickson, K. D., and A. B. Smith. 2023. Modeling the rarest of the rare: A comparison between multi-species distribution models, ensembles of small models, and single-species models at extremely low sample sizes. Ecography 2023:e06500.

Gomulkiewicz, R., and R. D. Holt. 1995. When does evolution by natural selection prevent extinction? Evolution 49:201–207.

Hart, S. P., and D. J. Marshall. 2013. Environmental stress, facilitation, competition, and coexistence. Ecology 94:2719–2731.

Ingram, M., D. Vukcevic, and N. Golding. 2020. Multi-output gaussian processes for species distribution modelling. Methods in Ecology and Evolution 11:1587–1598.

Ives, A. R. 1995. Predicting the response of populations to environmental change. Ecology 76:926–941.

Kloppers, P. H., and J. C. Greeff. 2013. Lotka–volterra model parameter estimation using experiential data. Applied Mathematics and Computation 224:817–825.

Mühlbauer, L. K., M. Schulze, W. S. Harpole, and A. T. Clark. 2020. gauseR: Simple methods for fitting lotka-volterra models describing gause's "struggle for existence". Ecology and Evolution 10:13275–13283.

Olivença, D. V., J. D. Davis, and E. O. Voit. 2021. Comparison between lotka-volterra and multivariate autoregressive models of ecological interaction systems. bioRxiv.

Ovaskainen, O., and N. Abrego. 2020. Joint species distribution modelling: With applications in r. Ecology biodiversity and conservation. Cambridge University Press, United Kingdom.

Ovaskainen, O., G. Tikhonov, D. Dunson, V. Grøtan, S. Engen, B.-E. Sæther, and N. Abrego. 2017. How are species interactions structured in species-rich communities? A new method for analysing time-series data. Proceedings of the Royal Society B: Biological Sciences 284:20170768.
