## Appendix S2 for "Detection of eco-evolutionary dynamics in communities using Joint Species Distribution Models"

### Appendix S2. Evolutionary drivers of species and community dynamics

Jelena H. Pantel* Ruben J. Hermann†

29 November, 2024

#### 1 Background

In Appendix S1.5, we evaluated the ability of the statistical model to estimate the impact of trait evolution for species abundances. Trait evolution's impact for species abundances can be modeled as a predictor in different ways. Here we derive the numerical form of the trait $x_i$ that we hypothesize has the most impact on species abundances.

#### 2 Model for population growth, competition, environmental change, and evolution

The growth equation for each species is:

$$N_{i,t+1} = \frac{\hat{W} e^{\frac{-[(\frac{w+(1-h^2)P}{P+w})(E-x_{i,t})]^2}{2(P+w)}} N_{i,t}}{1 + \alpha_{ii} N_{i,t} + \alpha_{ij} N_{j,t}}$$

We simulate population growth under particular parameter values, to generate population dynamics where species interactions, environment, and trait evolution all drive population dynamics. The simulation uses the `disc_LV_evol` command in the `ecoevor` package.

```
set.seed(42)
N0 <- c(10, 10)
alpha.11 <- 0.01
alpha.22 <- 0.01
alpha.12 <- 0.005
alpha.21 <- 0.01
alpha <- matrix(c(alpha.11, alpha.21, alpha.12, alpha.22), nrow = 2, byrow = FALSE)
E.0 <- 0.8
x.0 <- c(0.1, 0.8)
P <- 1
w <- 2
Wmax <- 2
```

*Laboratoire Chrono-environnement,UMR 6249 CNRS-UFC, 16 Route de Gray, 25030 Besançon cedex, France,

†University of Duisburg-Essen, Universitätsstraße 5, 45141 Essen, Germany,

```
h2 <- 1
k <- (w + (1 - h2) * P)/(P + w)
# Simulation of model for t time steps
t <- 40
N <- array(NA, dim = c(t, length(N0)))
N <- as.data.frame(N)
colnames(N) <- paste0("N", 1:length(N0))
N[1, ] <- N0
E <- rep(NA, t)
E[1] <- E.0
x <- array(NA, dim = c(t, length(N0)))
x <- as.data.frame(x)
colnames(x) <- paste0("x", 1:length(N0))
x[1, ] <- x.0
r <- array(NA, dim = c(t, length(N0)))
r <- as.data.frame(r)
colnames(r) <- paste0("r", 1:length(N0))
What <- Wmax * sqrt(w/(P + w))
r[1, ] <- What * exp((-(((w + (1 - h2) * P)/(P + w)) * (E[1] - x[1, ]))^2)/(2 * (P +
    w)))
for (i in 2:t) {
    res <- disc_LV_evol(N0 = N[i - 1, ], alpha = alpha, E = E[i - 1], x = x[i - 1,
        ], P = P, w = w, Wmax = Wmax, h2 = h2)
    N[i, ] <- res$Nt1
    r[i, ] <- res$r
    # trait change
    d <- E[i - 1] - x[i - 1, ]
    d1 <- k * d
    x[i, ] <- E[i - 1] - d1
    # environmental change
    E[i] <- E[i - 1] + abs(rnorm(1, 0, 0.05))
}
```

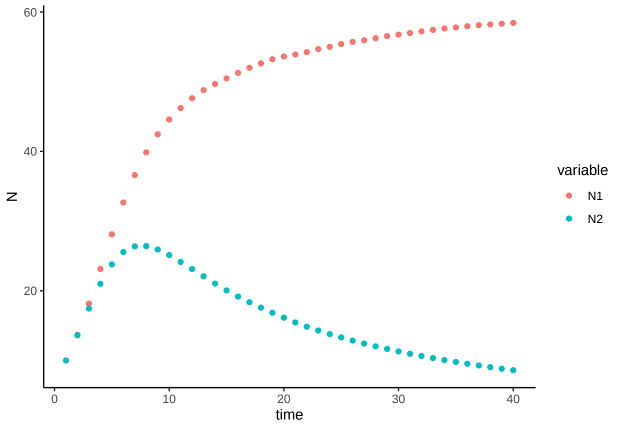


Figure 1: Population size $N$ over time $t$ for a discrete-time logistic growth model with competition, a changing environment, and trait evolution. Model parameters are $N_{i,0} = 10$, $\alpha_{ii} = 0.01$, $\alpha_{12} = 0.005$, $\alpha_{21} = 0.01$, and $h^2 = 1$.

16 We can check the impact evolution had for this system by re-running dynamics with $h^2 = 0$:

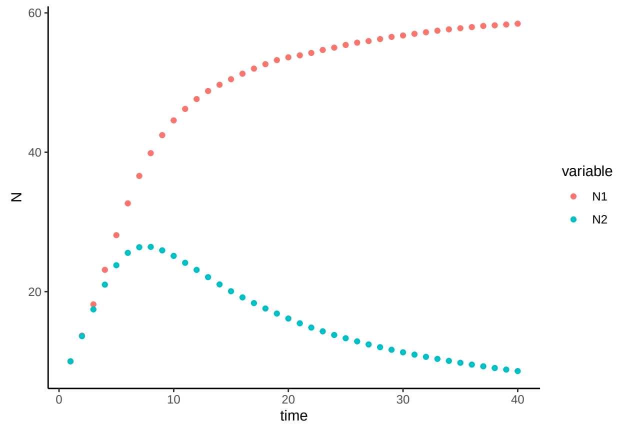


Figure 2: Population size $N$ over time $t$ for a discrete-time logistic growth model with competition, a changing environment, and no trait evolution. Model parameters are $N_{i,0} = 10$, $\alpha_{ii} = 0.01$, $\alpha_{12} = 0.005$, $\alpha_{21} = 0.01$, and $h^2 = 0$.

### 3 Hypotheses for evolution as a driver of species abundances

The growth equation is not linear, yet we fit abundance data to a hierarchical model with linear predictors because in most natural systems, a single theoretical growth model can't adequately capture the complexity in a given system. Linear models don't capture mechanistic relationships, but instead correlative. They are useful for inferring direction and magnitude of effect sizes. In Appendix S1, we explore how non-linear dynamics are captured when fit to a linear model. Here, we explore different forms to encode evolution in the trait $x$ that determines the organism's fitness in this system.

#### 3.1 H1. Raw trait value drives species abundances

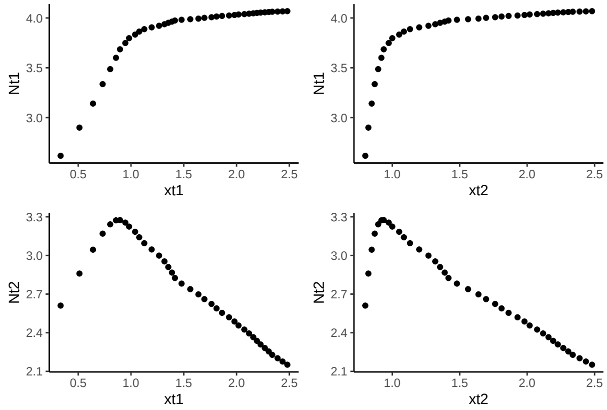


Figure 3: Population size (logarithm) in species 1 and 2 at one time step $N_{i,t+1}$ as a function of trait value of Species 1 and Species 2 in the previous time step $x_t$.

We do not hypothesize that the raw trait value is the best predictor of species abundances, because the impacts of this evolved trait depend on the context of the environment.

#### 3.2 H2. Absolute value of trait distance from optimum drives species abundances

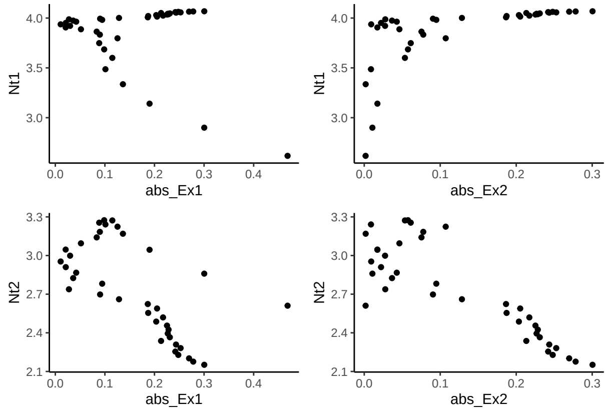


Figure 4: Population size (logarithm) in species 1 and 2 at one time step $N_{i,t+1}$ as a function of the absolute value of distance between the environmental optimum trait value given by $E_t$ and the trait value of Species 1 and Species 2 $x_t$.

We hypothesize that this represents the true impact on species abundances, via the direct impact this measure has on fitness and consequent population growth. However, we also acknowledge this is likely difficult to measure in most empirical systems.

#### 3.3 H3. Absolute value of trait change from one time to the next drives species abundances

We now consider the magnitude of trait change from one time step to the next as a driver for species abundances.

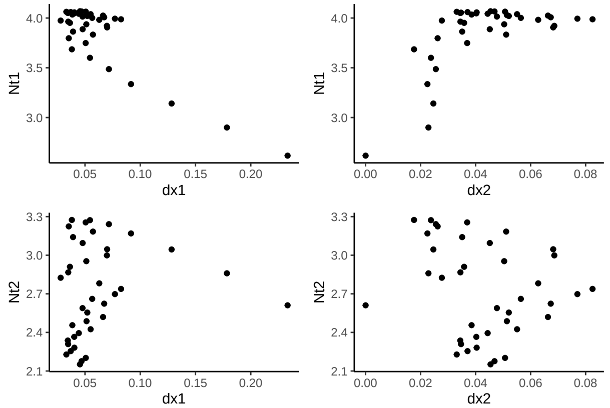


Figure 5: Population size (logarithm) in species 1 and 2 at one time step $N_{i,t+1}$ as a function of change in trait value of Species 1 and Species 2 in the previous time step $|\Delta x_{t,t-1}|$.

The measures correlate highly with one another, as the amount of trait change (the amount of evolution) is determined by the distance to the optimum trait value (given that P and w remain constant in the simulation).

```r
cor(df$dx1, df$abs_Ex1)
#> [1] 0.406851
cor(df$dx2, df$abs_Ex2)
#> [1] 0.1952239
```

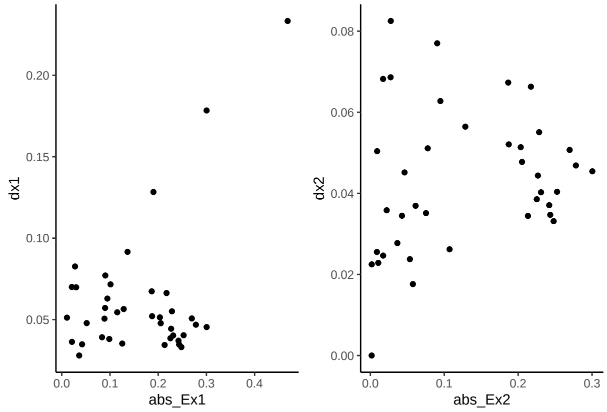


Figure 6: Relationship between change in trait value of Species 1 and Species 2 in the previous time step $|\Delta x_{t,t-1}|$ and distance from environmental optimum trait value to species trait value $|E_t - x_t|$.

### 4 Statistical model for evolution as a driver of species abundances

We hypothesize that $|E_t - x_{i,t}|$ will be the best predictor of species abundances, and that $|x_{t+1} - x_{i,t}|$ will have similar explanatory power.

```r
# prepare data in HMSC format
dat <- as.data.frame(cbind(log(N$N1), log(N$N2), x$x1, x$x2))
colnames(dat) <- c("N1", "N2", "x1", "x2")
dat$time <- 1:t
df <- data.frame(cbind(dat$N1[2:t], dat$N2[2:t], dat$x1[2:t], dat$x2[2:t]))
colnames(df) <- c("Nt1", "Nt2", "xt1", "xt2")
df$dx1 <- abs(dat$x1[2:t] - dat$x1[1:(t - 1)])
df$dx2 <- abs(dat$x2[2:t] - dat$x2[1:(t - 1)])
# Y matrix
Y <- as.matrix(cbind(df$Nt1, df$Nt2))
# X matrix
XData <- data.frame(cbind(dat$N1[1:(t - 1)], dat$N2[1:(t - 1)]), E[1:(t - 1)], E[1:(t -
    1)]^2, dat$x1[1:(t - 1)], dat$x2[1:(t - 1)], abs(dat$x1[2:t] - dat$x1[1:(t -
    1)]), abs(dat$x2[2:t] - dat$x2[1:(t - 1)]), abs(E[1:(t - 1)] - dat$x1[1:(t -
    1)]), abs(E[1:(t - 1)] - dat$x2[1:(t - 1)])))
colnames(XData) <- c("n1", "n2", "E", "Esq", "x1", "x2", "dx1", "dx2", "dEx1", "dEx2")
# Prepare HMSC model
studyDesign = data.frame(sample = as.factor(1:(t - 1)))
rL = HmscRandomLevel(units = studyDesign$sample)
# Design and fit 4 alternative HMSC models
models = list()
for (i in 1:4) {
    XFormula = switch(i, ~1, ~E + Esq + x1 + x2, ~E + Esq + dEx1 + dEx2, ~E + Esq +
```

```
        dx1 + dx2)
    m = Hmsc(Y = Y, XData = XData, XFormula = XFormula, studyDesign = studyDesign,
        ranLevels = list(sample = rL))
    models[[i]] = m
}
# Bayesian model parameters
nChains <- 2
thin <- 5
samples <- 2000
transient <- 100 * thin
verbose <- 500 * thin
for (i in 1:4) {
    models[[i]] = sampleMcmc(models[[i]], thin = thin, sample = samples, transient = transient,
        nChains = nChains, verbose = verbose)
}
```

We check the explanatory and predictive power of each model using cross-validation.

```
# After Ovaskainen & Abrego 2020, Chapter 7
partition = createPartition(m, nfolds = 2, column = "sample")
partition.sp = c(1, 2)
result = matrix(NA, nrow = 3, ncol = 4)
for (i in 1:4) {
    m = models[[i]]
    # Explanatory power
    preds = computePredictedValues(m)
    MF = evaluateModelFit(hM = m, predY = preds)
    result[1, i] = mean(MF$R2)
    # Predictive power based on cross-validation
    preds = computePredictedValues(m, partition = partition)
    MF = evaluateModelFit(hM = m, predY = preds)
    result[2, i] <- mean(MF$R2)
    # Predictive power based on conditional cross-validation
    preds = computePredictedValues(m, partition = partition, partition.sp = partition.sp,
        mcmcStep = 100)
    MF = evaluateModelFit(hM = m, predY = preds)
    result[3, i] = mean(MF$R2)
}
```

```
colnames(result) <- c("null", "xt", "dEx", "dx")
rownames(result) <- c("R2_expl", "R2_pred", "R2_pred_cross")
result
#>                      null        xt       dEx        dx
#> R2_expl        0.87023287 0.9973141 0.9947414 0.9960133
#> R2_pred       -0.01537272 0.9939087 0.9875206 0.9906780
#> R2_pred_cross  0.13943476 0.9939160 0.9879458 0.9906888
```

An important conclusion from this analysis is that the $|E_t - x_{i,t}|$ and $|x_{t+1} - x_{i,t}|$ do have similar explanatory power, and that - even though the species traits were the stronger predictor of species abundances - the model with total change in trait value can be used to assess the question "How does evolution impact species abundances?"

```r
m.post.hmsc <- convertToCodaObject(models[[4]])
bayesplot::mcmc_trace(m.post.hmsc$Beta)
```

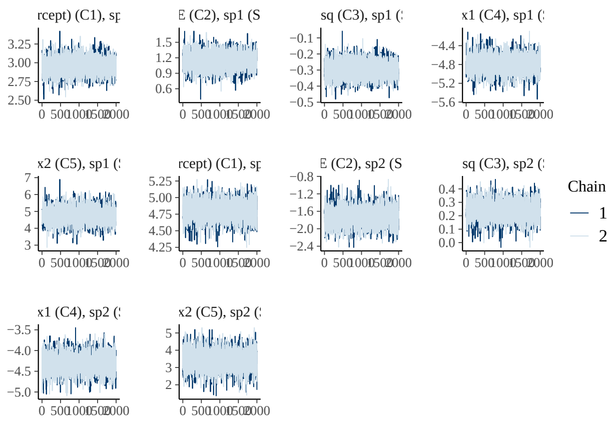


```r
bayesplot::mcmc_areas(m.post.hmsc$Beta, area_method = c("equal height"))
```

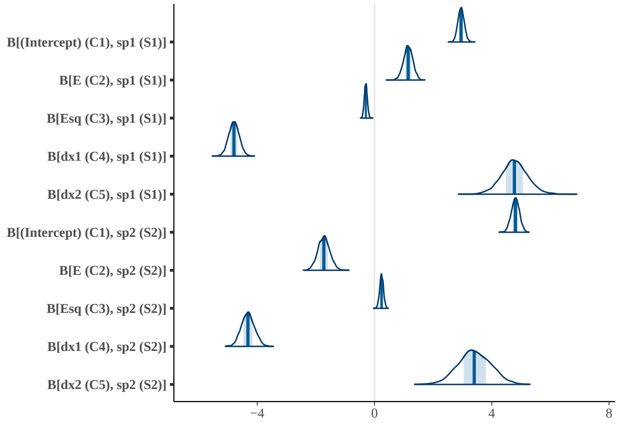


```r
# Bayesian estimates
cbind(summary(m.post.hmsc$Beta)$statistics[1:10, 1], summary(m.post.hmsc$Beta)$quantiles[,
    c(1, 5)])
#>                                              2.5%      97.5%
#> B[(Intercept) (C1), sp1 (S1)]  2.9521079  2.7485458  3.1660246
#> B[E (C2), sp1 (S1)]            1.1470368  0.8333290  1.4499025
#> B[Esq (C3), sp1 (S1)]         -0.2967196 -0.3951589 -0.1953133
#> B[dx1 (C4), sp1 (S1)]         -4.7933509 -5.1550466 -4.4359249
#> B[dx2 (C5), sp1 (S1)]          4.7750243  3.8733096  5.6787420
#> B[(Intercept) (C1), sp2 (S2)]  4.8026255  4.5270904  5.0716420
#> B[E (C2), sp2 (S2)]           -1.7264687 -2.1206607 -1.3233259
#> B[Esq (C3), sp2 (S2)]          0.2391028  0.1084473  0.3672534
#> B[dx1 (C4), sp2 (S2)]         -4.3164456 -4.7482722 -3.8745252
#> B[dx2 (C5), sp2 (S2)]          3.4113662  2.3122997  4.5109761
```

```r
OmegaCor = computeAssociations(models[[4]])
supportLevel = 0.7
toPlot = ((OmegaCor[[1]]$support > supportLevel) + (OmegaCor[[1]]$support < (1 -
```

```
    supportLevel)) > 0) * OmegaCor[[1]]$mean
corrplot::corrplot(toPlot, method = "color", col = colorRampPalette(c("blue", "white",
    "red"))(200), title = paste("random effect level:", models[[4]]$rLNames[1]),
    mar = c(0, 0, 1, 0))
```

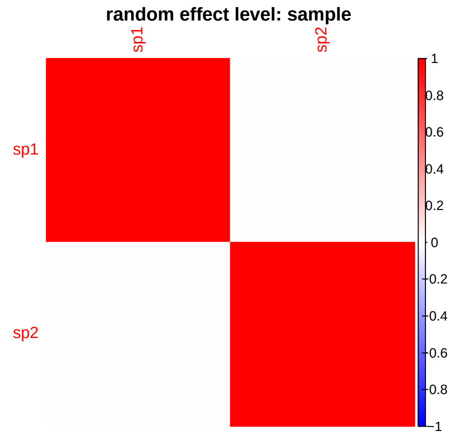


```
OmegaCor
#> [[1]]
#> [[1]]$mean
#>           sp1       sp2
#> sp1 1.0000000 0.1875533
#> sp2 0.1875533 1.0000000
#>
#> [[1]]$support
#>         sp1     sp2
#> sp1 1.00000 0.62075
#> sp2 0.62075 1.00000
```

We can also look at a gradient plot for the effect of evolution in Species 1 for abundance in Species 1 and Species 2.

```
Gradient <- constructGradient(models[[4]], focalVariable = "dx1", non.focalVariables = list(E = list(2)
    Esq = list(2), dx2 = list(2)), ngrid = 39)
Gradient$XDataNew$Esq <- Gradient$XDataNew$E^2
predY <- predict(models[[4]], XData = Gradient$XDataNew, expected = TRUE)
a <- plotGradient(models[[4]], Gradient, pred = predY, showData = T, measure = "Y",
    index = 1, main = "", xlab = "dx1_t", ylab = "predicted N1_t+1")
```

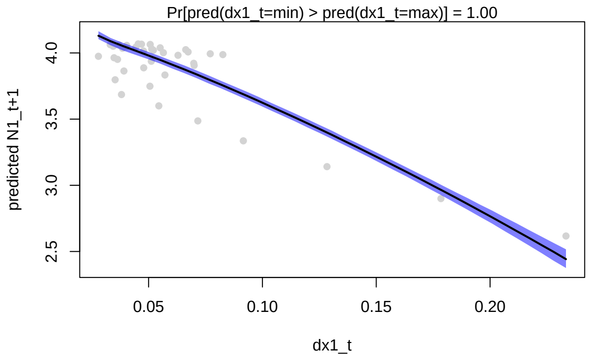


```r
b <- plotGradient(models[[4]], Gradient, pred = predY, showData = T, measure = "Y",
    index = 2, main = "", xlab = "dx1_t", ylab = "predicted N2_t+1")
```

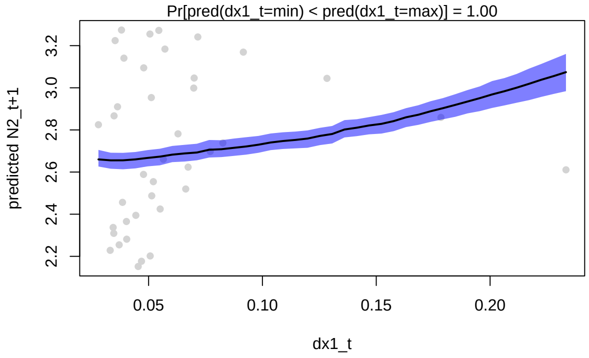


```r
postBeta = getPostEstimate(models[[4]], parName = "Beta")
plotBeta(models[[4]], post = postBeta, param = "Mean", supportLevel = 0.7)
```

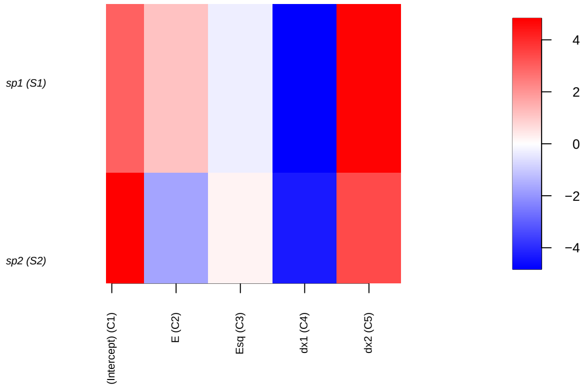


We repeat this plot for the effect of evolution in Species 2 for abundance in Species 1 and Species 2.

```r
Gradient <- constructGradient(models[[4]], focalVariable = "dx2", non.focalVariables = list(E = list(2)
    Esq = list(2), dx1 = list(2)), ngrid = 39)
Gradient$XDataNew$Esq <- Gradient$XDataNew$E^2
predY <- predict(models[[4]], XData = Gradient$XDataNew, expected = TRUE)
a <- plotGradient(models[[4]], Gradient, pred = predY, showData = T, measure = "Y",
    index = 1, main = "", xlab = "dx2_t", ylab = "predicted N1_t+1")
```

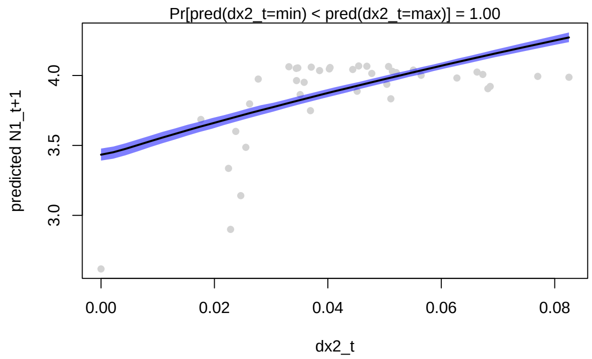


```r
b <- plotGradient(models[[4]], Gradient, pred = predY, showData = T, measure = "Y",
    index = 2, main = "", xlab = "dx2_t", ylab = "predicted N2_t+1")
```

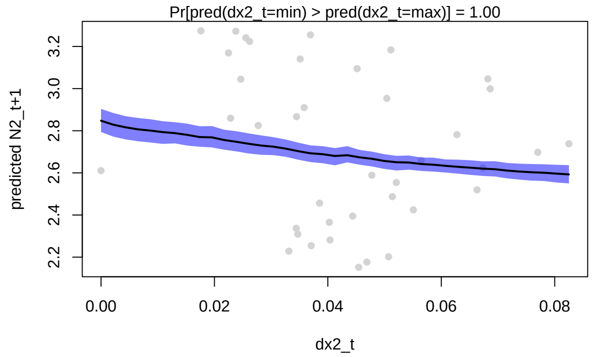


To evaluate the relative importance of evolution for species abundances, we use variation partition:

```r
VP <- computeVariancePartitioning(models[[4]], group = c(1, 1, 1, 2, 3), groupnames = c("Env",
    "Sp1", "Sp2"))
plotVariancePartitioning(models[[4]], VP, cols = c("white", "skyblue", "darkgrey",
    "orange"), args.legend = list(cex = 0.75, bg = "transparent"))
```

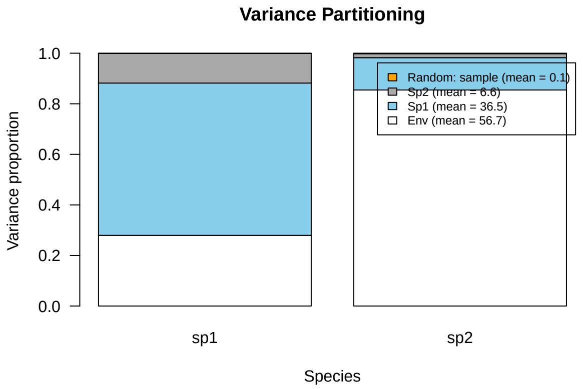


We see that trait evolution in Species 1 is an important driver of abundances in both Species 1 and Species 2.
