## Appendix S3 for "Detection of eco-evolutionary dynamics in communities using Joint Species Distribution Models"

### Appendix S3. The relative importance of evolution in populations and communities

Jelena H. Pantel* Ruben J. Hermann†

30 November, 2024

#### 1 Logistic growth and trait evolution

The goal of this analysis is to show that a hierarchical Bayesian linear model (HMSC) can successfully estimate the relative importance of different drivers of population dynamics, and to evaluate how the importance of evolution for population dynamics changes with heritability $h^2$ level. This analysis supports the findings for the **Evolution in metacommunities and HMSC** section in the main text. We create simulations of evolutionary rescue in a population, where adaptive evolution rescues a population from extinction, at a speed that depends on the heritability level in the system. Population dynamics are modelled as:

$$N_{i,t+1} = \frac{\hat{W} e^{\frac{-[(\frac{w+(1-h^2)P}{P+w})(E-x_{i,t})]^2}{2(P+w)}} N_{i,t}}{1+\alpha_{ii}N_{i,t}+\alpha_{ij}N_{j,t}}$$

where $\hat{W}$ is calculated as $\hat{W} = W_{max}\sqrt{(\frac{w}{P+w})}$, $W_{max}$ is the species' maximum per-capita growth rate, $w$ is the width of the Gaussian fitness function (which determines the strength of selection, as increasing values indicate a weaker reduction in fitness with distance from optimum trait value), $P$ is the width of the distribution of the phenotype $x$, and $h^2$ is the heritability of the trait $x$. For the simulation we use $W_{max} = 2$, $P = 1$, and $w = 2$.

The change in the average trait value each time step is given by:

$$d_{i,t+1} = kd_{i,t}$$

where $k = \frac{w+(1-h^2)P}{w+P}$ and $d_{i,t} = E_t - x_{i,t}$.

##### 1.1 Population dynamics simulation

We simulate population dynamics for 1 species, keeping all conditions constant except for heritability, which is evaluated at $h^2 = 0, 0.01, 0.02, 0.03, 0.04, 0.05, 0.075, 0.1, 0.4, 0.9, 1$. The simulation code for only $h^2 = 0$ is shown.

*Laboratoire Chrono-environnement,UMR 6249 CNRS-UFC, 16 Route de Gray, 25030 Besançon cedex, France,

†University of Duisburg-Essen, Universitätsstraße 5, 45141 Essen, Germany,

```
### 1 species, h2=0 ### Initial conditions
N0 <- 10
alpha.11 <- 0.01
alpha <- matrix(alpha.11, byrow = FALSE)
E.0 <- 0.8
P <- 1
w <- 0.5
# Draw initial trait value using degree of initial maladaptation
B0 <- 1
d0 = sqrt(B0 * (w + P))
x.0 = E.0 - d0
Wmax <- 2
h2 <- 0
k <- (w + (1 - h2) * P)/(P + w)
# Simulation of model for t time steps Simulation of model for t time steps
t <- 60
N <- array(NA, dim = c(t, length(N0)))
N <- as.data.frame(N)
colnames(N) <- paste0("N", 1:length(N0))
N[1, ] <- N0
E <- rep(NA, t)
E[1] <- E.0
x <- array(NA, dim = c(t, length(N0)))
x <- as.data.frame(x)
colnames(x) <- paste0("x", 1:length(N0))
x[1, ] <- x.0
r <- array(NA, dim = c(t, length(N0)))
r <- as.data.frame(r)
colnames(r) <- paste0("r", 1:length(N0))
What <- Wmax * sqrt(w/(P + w))
r[1, ] <- What * exp((-(((w + (1 - h2) * P)/(P + w)) * (E[1] - x[1, ]))^2)/(2 * (P +
    w)))
for (i in 2:t) {
    res <- disc_LV_evol(N0 = N[i - 1, ], alpha = alpha, E = E[i - 1], x = x[i - 1,
        ], P = P, w = w, Wmax = Wmax, h2 = h2)
    N[i, ] <- res$Nt1
    r[i, ] <- res$r
    # trait change
    d <- E[i - 1] - x[i - 1, ]
    d1 <- k * d
    x[i, ] <- E[i - 1] - d1
    # environmental change
    E[i] <- E[i - 1] + abs(rnorm(1, 0, 0))
}
# Arrange to save across scenarios
gdat <- N
gdat$time <- 1:t
gdat$h2 <- h2
gdat$x1 <- x$x1
gdat$E <- E
```

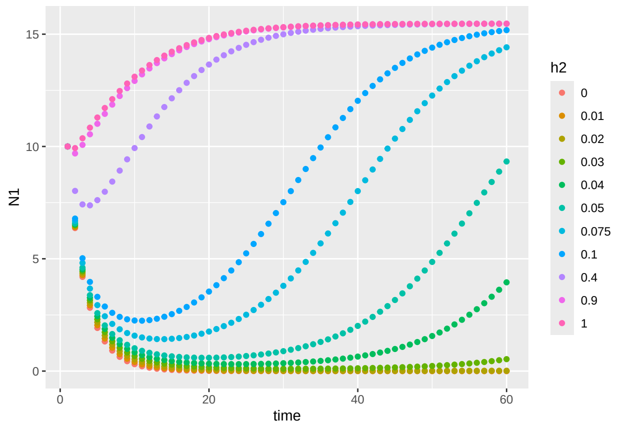


Figure 1: Plot of population size over time for a discete-time evolutionary rescue model, across a range of heritability $h^2$ values.

#### 1.2 HMSC model fit

The goal of fitting the data to a statistical model is to estimate the relative important of trait evolution for the population dynamics.

```
### Model 1, h2=00 ###
dat <- cbind(log(gdat$N1[gdat$h2 == 0]), gdat$x1[gdat$h2 == 0])
dat <- as.data.frame(dat)
colnames(dat) <- c("N1", "x1")
dat$time <- 1:t
dat <- as.data.frame(dat)
df <- data.frame(dat[(2:t), -3])
colnames(df) <- c("Nt1", "xt1")
df$dx1 <- abs(dat$x1[2:t] - dat$x1[1:(t - 1)])
# prepare data in HMSC format
Y <- as.matrix(df$Nt1)
XData <- data.frame(cbind(E[1:(t - 1)], E[1:(t - 1)]^2, abs(dat$x1[2:t] - dat$x1[1:(t -
    1)])))
colnames(XData) <- c("E", "Esq", "dx1")

studyDesign = data.frame(sample = as.factor(1:(t - 1)))
rL = HmscRandomLevel(units = studyDesign$sample)

m.1.hmsc = Hmsc(Y = Y, XData = XData, XFormula = ~E + Esq + dx1, studyDesign = studyDesign,
    ranLevels = list(sample = rL))
# Bayesian model parameters
nChains <- 2
thin <- 5
samples <- 2000
transient <- 1000 * thin
verbose <- 500 * thin
# sample MCMC
m.1.sample <- sampleMcmc(m.1.hmsc, thin = thin, sample = samples, transient = transient,
    nChains = nChains, verbose = verbose)
m1.post.hmsc <- convertToCodaObject(m.1.sample)
```

```
VP1 <- computeVariancePartitioning(m.1.sample, group = c(1, 1, 1, 2), groupnames = c("Env",
    "x1"))
plotVariancePartitioning(m.1.sample, VP1, cols = c("white", "skyblue", "darkgrey"),
    args.legend = list(cex = 0.75, bg = "transparent"))
```

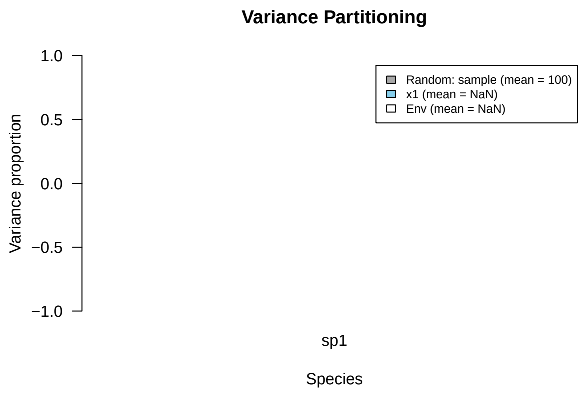


26

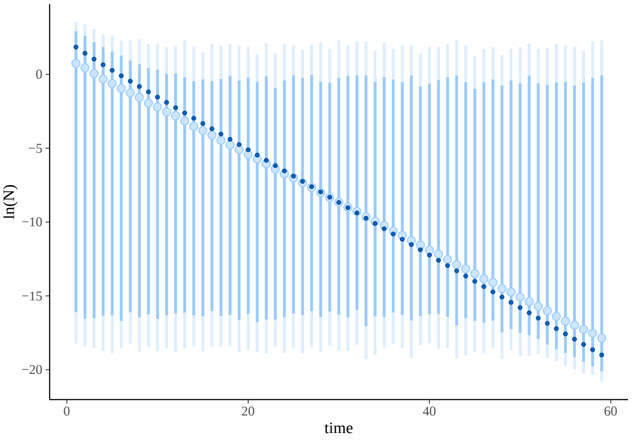


Figure 2: Simulated (dark blue points) population size over time for $h^2 = 0$, with posterior predicted mean (light blue points) and 95% highest density intervals (light blue bars).

```
### Model 2, h2=0.01 ###
dat <- cbind(log(gdat$N1[gdat$h2 == 0.01]), gdat$x1[gdat$h2 == 0.01])
dat <- as.data.frame(dat)
colnames(dat) <- c("N1", "x1")
dat$time <- 1:t
dat <- as.data.frame(dat)
df <- data.frame(dat[(2:t), -3])
colnames(df) <- c("Nt1", "xt1")
df$dx1 <- abs(dat$x1[2:t] - dat$x1[1:(t - 1)])
# prepare data in HMSC format
Y <- as.matrix(df$Nt1)
XData <- data.frame(cbind(E[1:(t - 1)], E[1:(t - 1)]^2, abs(dat$x1[2:t] - dat$x1[1:(t -
    1)])))
colnames(XData) <- c("E", "Esq", "dx1")

studyDesign = data.frame(sample = as.factor(1:(t - 1)))
```

```r
rL = HmscRandomLevel(units = studyDesign$sample)

m.2.hmsc = Hmsc(Y = Y, XData = XData, XFormula = ~E + Esq + dx1, studyDesign = studyDesign,
    ranLevels = list(sample = rL))
# Bayesian model parameters
nChains <- 2
thin <- 5
samples <- 2000
transient <- 1000 * thin
verbose <- 500 * thin
# sample MCMC
m.2.sample <- sampleMcmc(m.2.hmsc, thin = thin, sample = samples, transient = transient,
    nChains = nChains, verbose = verbose)
m2.post.hmsc <- convertToCodaObject(m.2.sample)
```

```r
m2.post.hmsc <- convertToCodaObject(m.2.sample)
summary(m2.post.hmsc$Beta)
#>
#> Iterations = 5005:15000
#> Thinning interval = 5
#> Number of chains = 2
#> Sample size per chain = 2000
#>
#> 1. Empirical mean and standard deviation for each variable,
#>    plus standard error of the mean:
#>
#>                                      Mean        SD  Naive SE Time-series SE
#> B[(Intercept) (C1), sp1 (S1)] -1.178e+15 6.264e+15 9.905e+13      1.566e+14
#> B[E (C2), sp1 (S1)]           -2.644e+15 3.470e+15 5.487e+13      9.209e+13
#> B[Esq (C3), sp1 (S1)]          5.145e+15 6.926e+15 1.095e+14      1.825e+14
#> B[dx1 (C4), sp1 (S1)]          4.345e+03 7.368e+01 1.165e+00      1.166e+00
#>
#> 2. Quantiles for each variable:
#>
#>                                      2.5%        25%        50%        75%
#> B[(Intercept) (C1), sp1 (S1)] -1.372e+16 -4.631e+15 -1.204e+15  2.190e+15
#> B[E (C2), sp1 (S1)]           -9.429e+15 -4.464e+15 -2.656e+15 -7.931e+14
#> B[Esq (C3), sp1 (S1)]         -8.793e+15  1.469e+15  5.155e+15  8.887e+15
#> B[dx1 (C4), sp1 (S1)]          4.198e+03  4.296e+03  4.345e+03  4.394e+03
#>                                    97.5%
#> B[(Intercept) (C1), sp1 (S1)] 1.140e+16
#> B[E (C2), sp1 (S1)]           4.298e+15
#> B[Esq (C3), sp1 (S1)]         1.913e+16
#> B[dx1 (C4), sp1 (S1)]         4.489e+03
bayesplot::mcmc_trace(m2.post.hmsc$Beta)
```

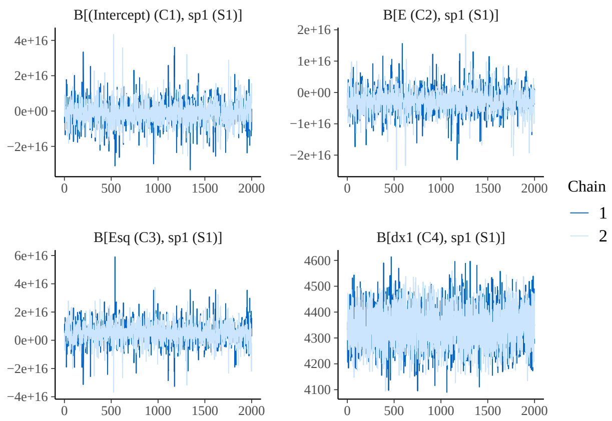


```
bayesplot::mcmc_areas(m2.post.hmsc$Beta, area_method = c("equal height"))
```

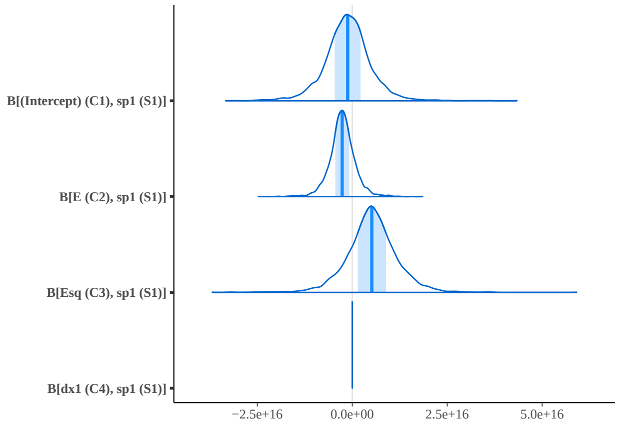


```
Gradient <- constructGradient(m.2.sample, focalVariable = "dx1", non.focalVariables = list(E = list(2),
    Esq = list(2)), ngrid = 59)
# Esq is manually constructed as gradient-produced E^2
Gradient$XDataNew$Esq <- Gradient$XDataNew$E^2
predY <- predict(m.2.sample, XData = Gradient$XDataNew, expected = TRUE)
plotGradient(m.2.sample, Gradient, pred = predY, showData = T, measure = "Y", main = "",
    xlab = "dx1_t", ylab = "predicted N_t+1")
```

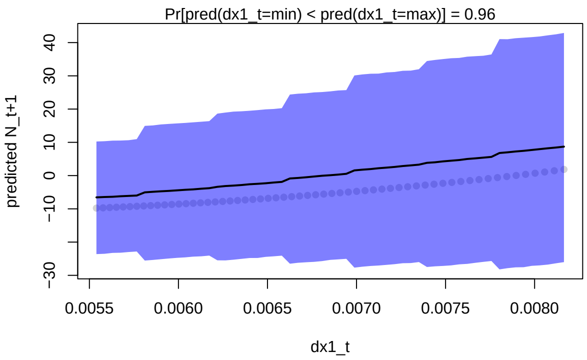

```
#> [1] 0.9595
```

We can see that the system is unpredictable, as population size is extremely low during the simulation due to maladaptation.

```r
VP2 <- computeVariancePartitioning(m.2.sample, group = c(1, 1, 1, 2), groupnames = c("Env",
    "x1"))
plotVariancePartitioning(m.2.sample, VP2, cols = c("white", "skyblue", "darkgrey"),
    args.legend = list(cex = 0.75, bg = "transparent"))
```

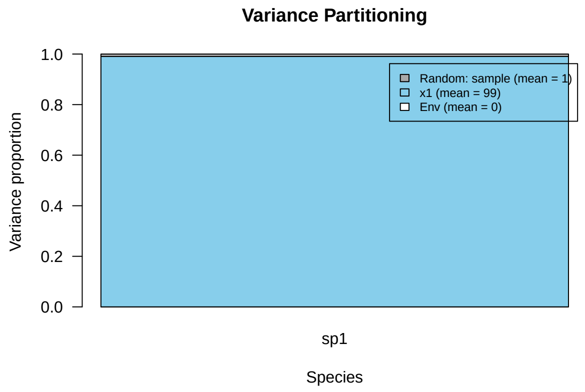


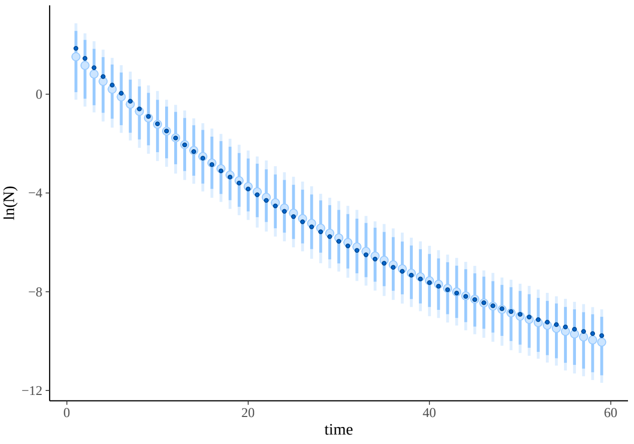


Figure 3: Simulated (dark blue points) population size over time for $h^2 = 0.01$, with posterior predicted mean (light blue points) and 95% highest density intervals (light blue bars).

```r
### Model 3, h2=0.02 ###
dat <- cbind(log(gdat$N1[gdat$h2 == 0.02]), gdat$x1[gdat$h2 == 0.02])
dat <- as.data.frame(dat)
colnames(dat) <- c("N1", "x1")
dat$time <- 1:t
dat <- as.data.frame(dat)
df <- data.frame(dat[(2:t), -3])
colnames(df) <- c("Nt1", "xt1")
df$dx1 <- abs(dat$x1[2:t] - dat$x1[1:(t - 1)])
# prepare data in HMSC format
Y <- as.matrix(df$Nt1)
```

```
XData <- data.frame(cbind(E[1:(t - 1)], E[1:(t - 1)]^2, abs(dat$x1[2:t] - dat$x1[1:(t -
    1)])))
colnames(XData) <- c("E", "Esq", "dx1")

studyDesign = data.frame(sample = as.factor(1:(t - 1)))
rL = HmscRandomLevel(units = studyDesign$sample)

m.3.hmsc = Hmsc(Y = Y, XData = XData, XFormula = ~E + Esq + dx1, studyDesign = studyDesign,
    ranLevels = list(sample = rL))
# Bayesian model parameters
nChains <- 2
thin <- 5
samples <- 2000
transient <- 1000 * thin
verbose <- 500 * thin
# sample MCMC
m.3.sample <- sampleMcmc(m.3.hmsc, thin = thin, sample = samples, transient = transient,
    nChains = nChains, verbose = verbose)
m3.post.hmsc <- convertToCodaObject(m.3.sample)
```

```
VP3 <- computeVariancePartitioning(m.3.sample, group = c(1, 1, 1, 2), groupnames = c("Env",
    "x1"))
plotVariancePartitioning(m.3.sample, VP3, cols = c("white", "skyblue", "darkgrey"),
    args.legend = list(cex = 0.75, bg = "transparent"))
```

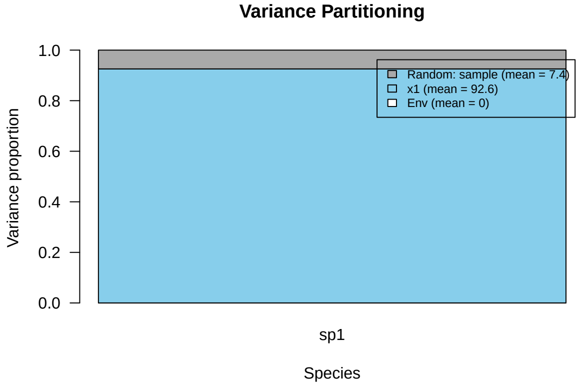

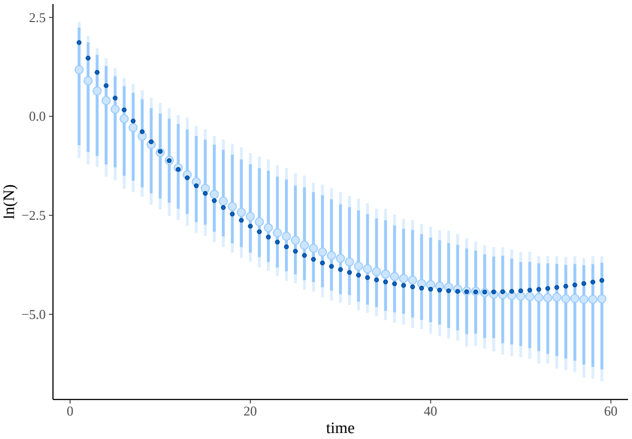


Figure 4: Simulated (dark blue points) population size over time for $h^2 = 0.02$, with posterior predicted mean (light blue points) and 95% highest density intervals (light blue bars).

```
### Model 4, h2=0.03 ###
dat <- cbind(log(gdat$N1[gdat$h2 == 0.03]), gdat$x1[gdat$h2 == 0.03])
dat <- as.data.frame(dat)
colnames(dat) <- c("N1", "x1")
dat$time <- 1:t
dat <- as.data.frame(dat)
df <- data.frame(dat[(2:t), -3])
colnames(df) <- c("Nt1", "xt1")
df$dx1 <- abs(dat$x1[2:t] - dat$x1[1:(t - 1)])
# prepare data in HMSC format
Y <- as.matrix(df$Nt1)
XData <- data.frame(cbind(E[1:(t - 1)], E[1:(t - 1)]^2, abs(dat$x1[2:t] - dat$x1[1:(t -
    1)])))
colnames(XData) <- c("E", "Esq", "dx1")

studyDesign = data.frame(sample = as.factor(1:(t - 1)))
rL = HmscRandomLevel(units = studyDesign$sample)

m.4.hmsc = Hmsc(Y = Y, XData = XData, XFormula = ~E + Esq + dx1, studyDesign = studyDesign,
    ranLevels = list(sample = rL))
# Bayesian model parameters
nChains <- 2
thin <- 5
samples <- 2000
transient <- 1000 * thin
verbose <- 500 * thin
# sample MCMC
m.4.sample <- sampleMcmc(m.4.hmsc, thin = thin, sample = samples, transient = transient,
    nChains = nChains, verbose = verbose)
m4.post.hmsc <- convertToCodaObject(m.4.sample)
```

```
VP4 <- computeVariancePartitioning(m.4.sample, group = c(1, 1, 1, 2), groupnames = c("Env",
    "x1"))
plotVariancePartitioning(m.4.sample, VP4, cols = c("white", "skyblue", "darkgrey"),
    args.legend = list(cex = 0.75, bg = "transparent"))
```

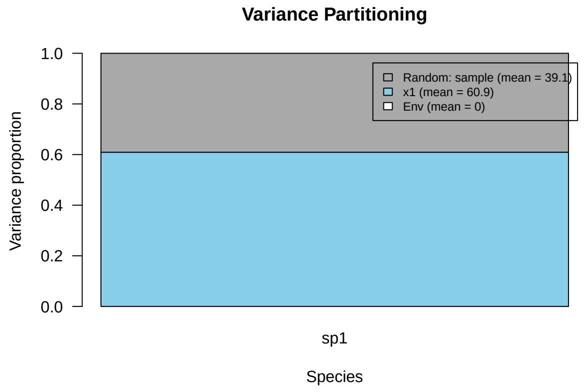


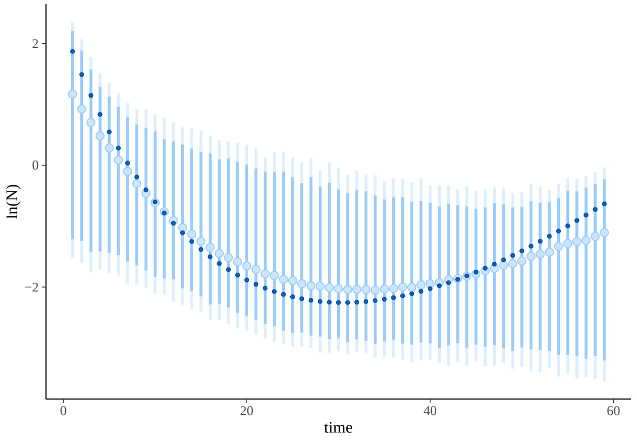


Figure 5: Simulated (dark blue points) population size over time for $h^2 = 0.03$, with posterior predicted mean (light blue points) and 95% highest density intervals (light blue bars).

```
### Model 5, h2=0.04 ###
dat <- cbind(log(gdat$N1[gdat$h2 == 0.04]), gdat$x1[gdat$h2 == 0.04])
dat <- as.data.frame(dat)
colnames(dat) <- c("N1", "x1")
dat$time <- 1:t
dat <- as.data.frame(dat)
df <- data.frame(dat[(2:t), -3])
colnames(df) <- c("Nt1", "xt1")
df$dx1 <- abs(dat$x1[2:t] - dat$x1[1:(t - 1)])
# prepare data in HMSC format
Y <- as.matrix(df$Nt1)
XData <- data.frame(cbind(E[1:(t - 1)], E[1:(t - 1)]^2, abs(dat$x1[2:t] - dat$x1[1:(t -
    1)])))
colnames(XData) <- c("E", "Esq", "dx1")

studyDesign = data.frame(sample = as.factor(1:(t - 1)))
rL = HmscRandomLevel(units = studyDesign$sample)

m.5.hmsc = Hmsc(Y = Y, XData = XData, XFormula = ~E + Esq + dx1, studyDesign = studyDesign,
    ranLevels = list(sample = rL))
# Bayesian model parameters
```

```
nChains <- 2
thin <- 5
samples <- 2000
transient <- 1000 * thin
verbose <- 500 * thin
# sample MCMC
m.5.sample <- sampleMcmc(m.5.hmsc, thin = thin, sample = samples, transient = transient,
    nChains = nChains, verbose = verbose)
m5.post.hmsc <- convertToCodaObject(m.5.sample)
```

```
VP5 <- computeVariancePartitioning(m.5.sample, group = c(1, 1, 1, 2), groupnames = c("Env",
    "x1"))
plotVariancePartitioning(m.5.sample, VP5, cols = c("white", "skyblue", "darkgrey"),
    args.legend = list(cex = 0.75, bg = "transparent"))
```

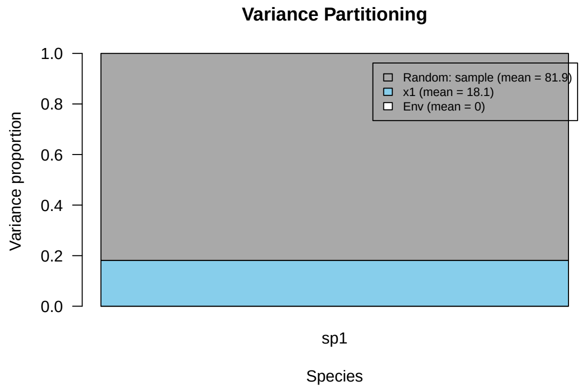


36

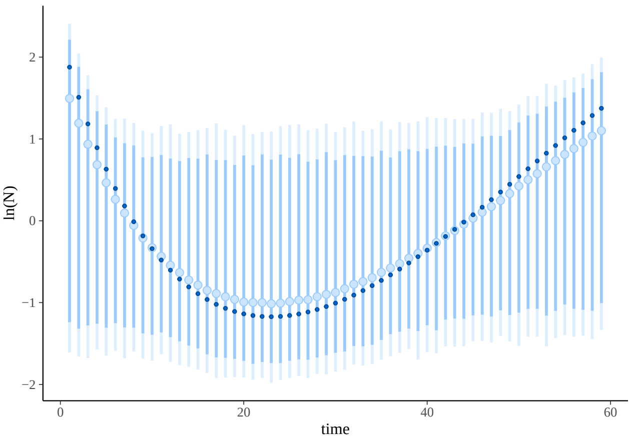


Figure 6: Simulated (dark blue points) population size over time for $h^2 = 0.01$, with posterior predicted mean (light blue points) and 95% highest density intervals (light blue bars).

```
### Model 6, h2=0.05 ###
dat <- cbind(log(gdat$N1[gdat$h2 == 0.05]), gdat$x1[gdat$h2 == 0.05])
dat <- as.data.frame(dat)
colnames(dat) <- c("N1", "x1")
dat$time <- 1:t
```

```
dat <- as.data.frame(dat)
df <- data.frame(dat[(2:t), -3])
colnames(df) <- c("Nt1", "xt1")
df$dx1 <- abs(dat$x1[2:t] - dat$x1[1:(t - 1)])
# prepare data in HMSC format
Y <- as.matrix(df$Nt1)
XData <- data.frame(cbind(E[1:(t - 1)], E[1:(t - 1)]^2, abs(dat$x1[2:t] - dat$x1[1:(t -
    1)])))
colnames(XData) <- c("E", "Esq", "dx1")

studyDesign = data.frame(sample = as.factor(1:(t - 1)))
rL = HmscRandomLevel(units = studyDesign$sample)

m.6.hmsc = Hmsc(Y = Y, XData = XData, XFormula = ~E + Esq + dx1, studyDesign = studyDesign,
    ranLevels = list(sample = rL))
# Bayesian model parameters
nChains <- 2
thin <- 5
samples <- 2000
transient <- 1000 * thin
verbose <- 500 * thin
# sample MCMC
m.6.sample <- sampleMcmc(m.6.hmsc, thin = thin, sample = samples, transient = transient,
    nChains = nChains, verbose = verbose)
m6.post.hmsc <- convertToCodaObject(m.6.sample)
```

```
VP6 <- computeVariancePartitioning(m.6.sample, group = c(1, 1, 1, 2), groupnames = c("Env",
    "x1"))
plotVariancePartitioning(m.6.sample, VP6, cols = c("white", "skyblue", "darkgrey"),
    args.legend = list(cex = 0.75, bg = "transparent"))
```

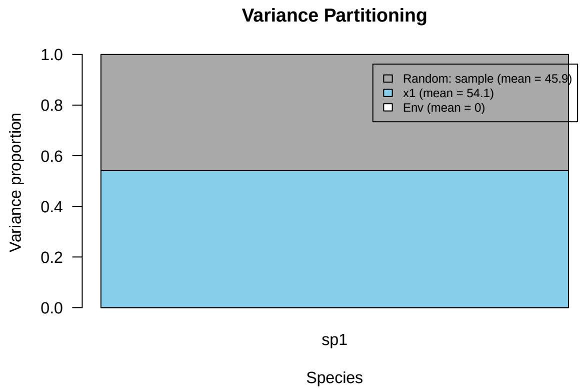

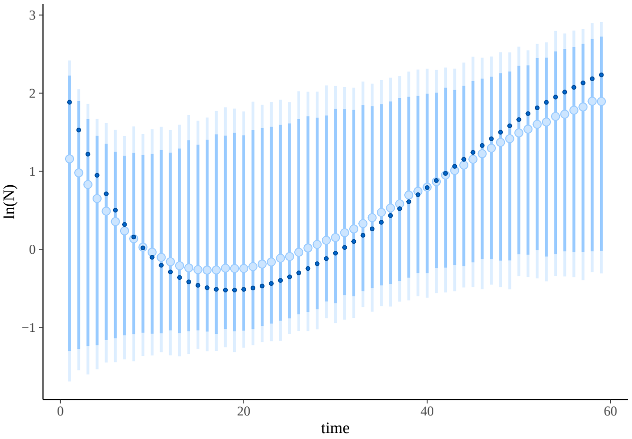


Figure 7: Simulated (dark blue points) population size over time for $h^2 = 0.02$, with posterior predicted mean (light blue points) and 95% highest density intervals (light blue bars).

```
### Model 7, h2=0.075 ###
dat <- cbind(log(gdat$N1[gdat$h2 == 0.075]), gdat$x1[gdat$h2 == 0.075])
dat <- as.data.frame(dat)
colnames(dat) <- c("N1", "x1")
dat$time <- 1:t
dat <- as.data.frame(dat)
df <- data.frame(dat[(2:t), -3])
colnames(df) <- c("Nt1", "xt1")
df$dx1 <- abs(dat$x1[2:t] - dat$x1[1:(t - 1)])
# prepare data in HMSC format
Y <- as.matrix(df$Nt1)
XData <- data.frame(cbind(E[1:(t - 1)], E[1:(t - 1)]^2, abs(dat$x1[2:t] - dat$x1[1:(t -
    1)])))
colnames(XData) <- c("E", "Esq", "dx1")

studyDesign = data.frame(sample = as.factor(1:(t - 1)))
rL = HmscRandomLevel(units = studyDesign$sample)

m.7.hmsc = Hmsc(Y = Y, XData = XData, XFormula = ~E + Esq + dx1, studyDesign = studyDesign,
    ranLevels = list(sample = rL))
# Bayesian model parameters
nChains <- 2
thin <- 5
samples <- 2000
transient <- 1000 * thin
verbose <- 500 * thin
# sample MCMC
m.7.sample <- sampleMcmc(m.7.hmsc, thin = thin, sample = samples, transient = transient,
    nChains = nChains, verbose = verbose)
m7.post.hmsc <- convertToCodaObject(m.7.sample)
```

```
VP7 <- computeVariancePartitioning(m.7.sample, group = c(1, 1, 1, 2), groupnames = c("Env",
    "x1"))
plotVariancePartitioning(m.7.sample, VP7, cols = c("white", "skyblue", "darkgrey"),
    args.legend = list(cex = 0.75, bg = "transparent"))
```

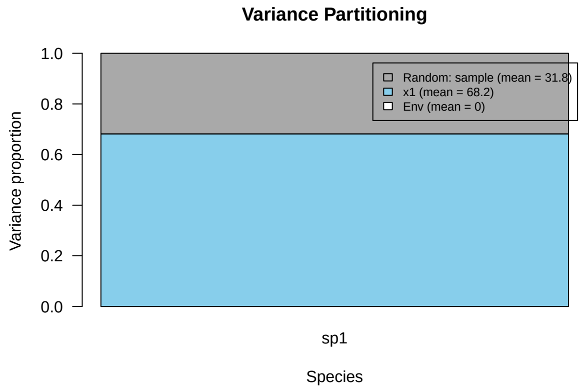


38

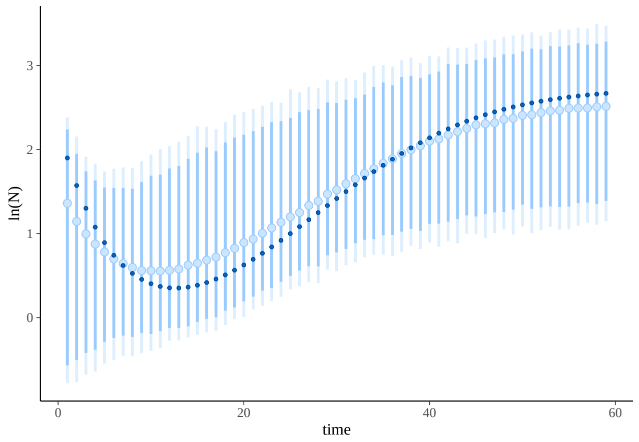


Figure 8: Simulated (dark blue points) population size over time for $h^2 = 0.075$, with posterior predicted mean (light blue points) and 95% highest density intervals (light blue bars).

```
### Model 8, h2=0.1 ###
dat <- cbind(log(gdat$N1[gdat$h2 == 0.1]), gdat$x1[gdat$h2 == 0.1])
dat <- as.data.frame(dat)
colnames(dat) <- c("N1", "x1")
dat$time <- 1:t
dat <- as.data.frame(dat)
df <- data.frame(dat[(2:t), -3])
colnames(df) <- c("Nt1", "xt1")
df$dx1 <- abs(dat$x1[2:t] - dat$x1[1:(t - 1)])
# prepare data in HMSC format
Y <- as.matrix(df$Nt1)
XData <- data.frame(cbind(E[1:(t - 1)], E[1:(t - 1)]^2, abs(dat$x1[2:t] - dat$x1[1:(t -
    1)])))
colnames(XData) <- c("E", "Esq", "dx1")

studyDesign = data.frame(sample = as.factor(1:(t - 1)))
rL = HmscRandomLevel(units = studyDesign$sample)

m.8.hmsc = Hmsc(Y = Y, XData = XData, XFormula = ~E + Esq + dx1, studyDesign = studyDesign,
    ranLevels = list(sample = rL))
# Bayesian model parameters
```

```
nChains <- 2
thin <- 5
samples <- 2000
transient <- 1000 * thin
verbose <- 500 * thin
# sample MCMC
m.8.sample <- sampleMcmc(m.8.hmsc, thin = thin, sample = samples, transient = transient,
    nChains = nChains, verbose = verbose)
m8.post.hmsc <- convertToCodaObject(m.8.sample)
```

```
VP8 <- computeVariancePartitioning(m.8.sample, group = c(1, 1, 1, 2), groupnames = c("Env",
    "x1"))
plotVariancePartitioning(m.8.sample, VP8, cols = c("white", "skyblue", "darkgrey"),
    args.legend = list(cex = 0.75, bg = "transparent"))
```

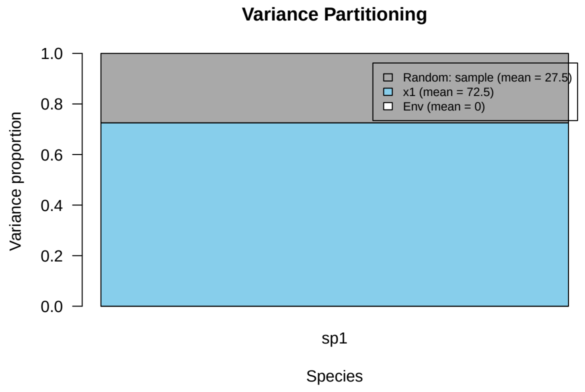


39

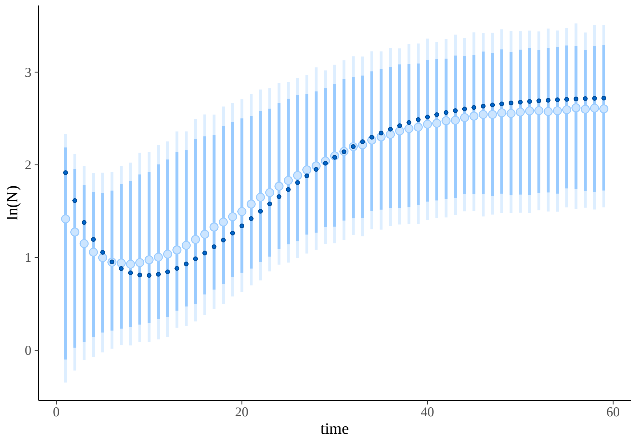


Figure 9: Simulated (dark blue points) population size over time for $h^2 = 1$, with posterior predicted mean (light blue points) and 95% highest density intervals (light blue bars).

```
### Model 9, h2=0.4 ###
dat <- cbind(log(gdat$N1[gdat$h2 == 0.4]), gdat$x1[gdat$h2 == 0.4])
dat <- as.data.frame(dat)
colnames(dat) <- c("N1", "x1")
dat$time <- 1:t
```

```r
dat <- as.data.frame(dat)
df <- data.frame(dat[(2:t), -3])
colnames(df) <- c("Nt1", "xt1")
df$dx1 <- abs(dat$x1[2:t] - dat$x1[1:(t - 1)])
# prepare data in HMSC format
Y <- as.matrix(df$Nt1)
XData <- data.frame(cbind(E[1:(t - 1)], E[1:(t - 1)]^2, abs(dat$x1[2:t] - dat$x1[1:(t -
    1)])))
colnames(XData) <- c("E", "Esq", "dx1")

studyDesign = data.frame(sample = as.factor(1:(t - 1)))
rL = HmscRandomLevel(units = studyDesign$sample)

m.9.hmsc = Hmsc(Y = Y, XData = XData, XFormula = ~E + Esq + dx1, studyDesign = studyDesign,
    ranLevels = list(sample = rL))
# Bayesian model parameters
nChains <- 2
thin <- 5
samples <- 2000
transient <- 1000 * thin
verbose <- 500 * thin
# sample MCMC
m.9.sample <- sampleMcmc(m.9.hmsc, thin = thin, sample = samples, transient = transient,
    nChains = nChains, verbose = verbose)
m9.post.hmsc <- convertToCodaObject(m.9.sample)
```

```r
VP9 <- computeVariancePartitioning(m.9.sample, group = c(1, 1, 1, 2), groupnames = c("Env",
    "x1"))
plotVariancePartitioning(m.9.sample, VP9, cols = c("white", "skyblue", "darkgrey"),
    args.legend = list(cex = 0.75, bg = "transparent"))
```

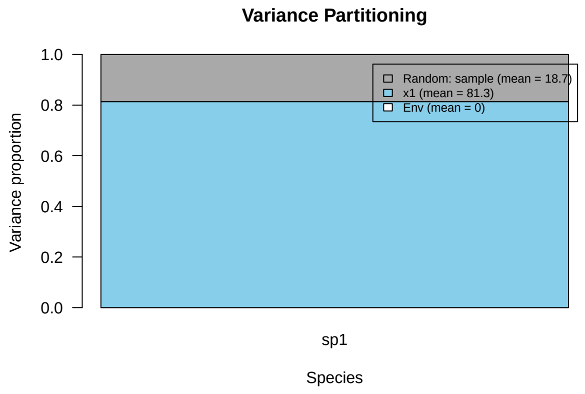

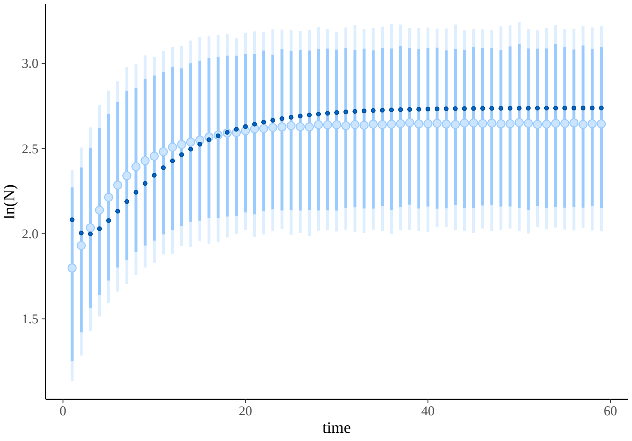


Figure 10: Simulated (dark blue points) population size over time for $h^2 = 0.4$, with posterior predicted mean (light blue points) and 95% highest density intervals (light blue bars).

```
### Model 10, h2=0.9 ###
dat <- cbind(log(gdat$N1[gdat$h2 == 0.9]), gdat$x1[gdat$h2 == 0.9])
dat <- as.data.frame(dat)
colnames(dat) <- c("N1", "x1")
dat$time <- 1:t
dat <- as.data.frame(dat)
df <- data.frame(dat[(2:t), -3])
colnames(df) <- c("Nt1", "xt1")
df$dx1 <- abs(dat$x1[2:t] - dat$x1[1:(t - 1)])
# prepare data in HMSC format
Y <- as.matrix(df$Nt1)
XData <- data.frame(cbind(E[1:(t - 1)], E[1:(t - 1)]^2, abs(dat$x1[2:t] - dat$x1[1:(t -
    1)])))
colnames(XData) <- c("E", "Esq", "dx1")

studyDesign = data.frame(sample = as.factor(1:(t - 1)))
rL = HmscRandomLevel(units = studyDesign$sample)

m.10.hmsc = Hmsc(Y = Y, XData = XData, XFormula = ~E + Esq + dx1, studyDesign = studyDesign,
    ranLevels = list(sample = rL))
# Bayesian model parameters
nChains <- 2
thin <- 5
samples <- 2000
transient <- 1000 * thin
verbose <- 500 * thin
# sample MCMC
m.10.sample <- sampleMcmc(m.10.hmsc, thin = thin, sample = samples, transient = transient,
    nChains = nChains, verbose = verbose)
m10.post.hmsc <- convertToCodaObject(m.10.sample)
```

```
VP10 <- computeVariancePartitioning(m.10.sample, group = c(1, 1, 1, 2), groupnames = c("Env",
    "x1"))
plotVariancePartitioning(m.10.sample, VP10, cols = c("white", "skyblue", "darkgrey"),
    args.legend = list(cex = 0.75, bg = "transparent"))
```

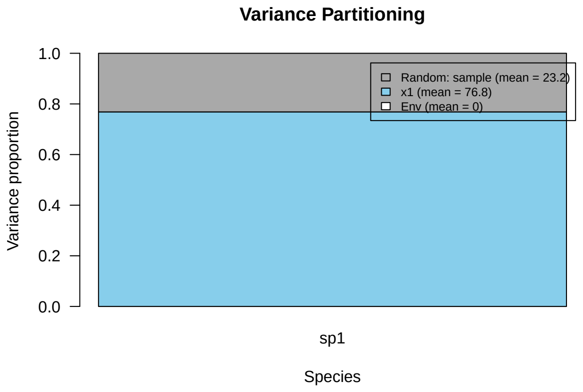


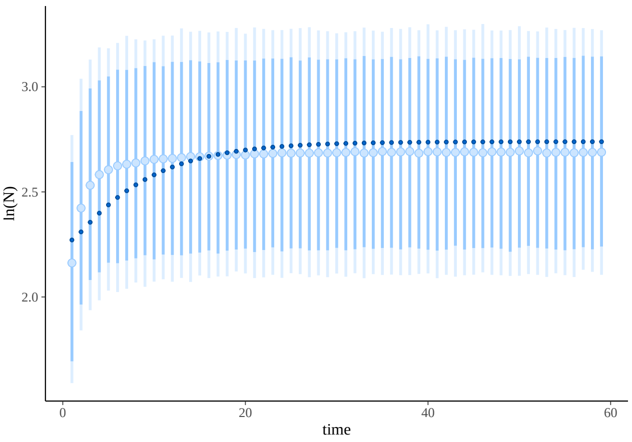


Figure 11: Simulated (dark blue points) population size over time for $h^2 = 0.9$, with posterior predicted mean (light blue points) and 95% highest density intervals (light blue bars).

```
### Model 11, h2=1 ###
dat <- cbind(log(gdat$N1[gdat$h2 == 1]), gdat$x1[gdat$h2 == 1])
dat <- as.data.frame(dat)
colnames(dat) <- c("N1", "x1")
dat$time <- 1:t
dat <- as.data.frame(dat)
df <- data.frame(dat[(2:t), -3])
colnames(df) <- c("Nt1", "xt1")
df$dx1 <- abs(dat$x1[2:t] - dat$x1[1:(t - 1)])
# prepare data in HMSC format
Y <- as.matrix(df$Nt1)
XData <- data.frame(cbind(E[1:(t - 1)], E[1:(t - 1)]^2, abs(dat$x1[2:t] - dat$x1[1:(t -
    1)])))
colnames(XData) <- c("E", "Esq", "dx1")

studyDesign = data.frame(sample = as.factor(1:(t - 1)))
rL = HmscRandomLevel(units = studyDesign$sample)

m.11.hmsc = Hmsc(Y = Y, XData = XData, XFormula = ~E + Esq + dx1, studyDesign = studyDesign,
    ranLevels = list(sample = rL))
# Bayesian model parameters
```

```
nChains <- 2
thin <- 5
samples <- 2000
transient <- 1000 * thin
verbose <- 500 * thin
# sample MCMC
m.11.sample <- sampleMcmc(m.11.hmsc, thin = thin, sample = samples, transient = transient,
    nChains = nChains, verbose = verbose)
m11.post.hmsc <- convertToCodaObject(m.11.sample)
```

```
m11.post.hmsc <- convertToCodaObject(m.11.sample)
summary(m11.post.hmsc$Beta)
#>
#> Iterations = 5005:15000
#> Thinning interval = 5
#> Number of chains = 2
#> Sample size per chain = 2000
#>
#> 1. Empirical mean and standard deviation for each variable,
#>    plus standard error of the mean:
#>
#>                                    Mean        SD  Naive SE Time-series SE
#> B[(Intercept) (C1), sp1 (S1)]  6.626e+14 3.185e+15 5.036e+13      5.037e+13
#> B[E (C2), sp1 (S1)]            1.322e+15 1.749e+15 2.765e+13      2.765e+13
#> B[Esq (C3), sp1 (S1)]         -2.688e+15 3.551e+15 5.615e+13      5.615e+13
#> B[dx1 (C4), sp1 (S1)]         -6.066e-01 1.036e-01 1.639e-03      1.639e-03
#>
#> 2. Quantiles for each variable:
#>
#>                                    2.5%        25%        50%        75%
#> B[(Intercept) (C1), sp1 (S1)] -5.464e+15 -1.346e+15  6.827e+14  2.667e+15
#> B[E (C2), sp1 (S1)]           -2.066e+15  2.588e+14  1.298e+15  2.399e+15
#> B[Esq (C3), sp1 (S1)]         -9.582e+15 -4.847e+15 -2.729e+15 -4.805e+14
#> B[dx1 (C4), sp1 (S1)]         -8.063e-01 -6.774e-01 -6.070e-01 -5.368e-01
#>                                   97.5%
#> B[(Intercept) (C1), sp1 (S1)]  6.735e+15
#> B[E (C2), sp1 (S1)]            4.812e+15
#> B[Esq (C3), sp1 (S1)]          4.319e+15
#> B[dx1 (C4), sp1 (S1)]         -3.982e-01
bayesplot::mcmc_trace(m11.post.hmsc$Beta)
```

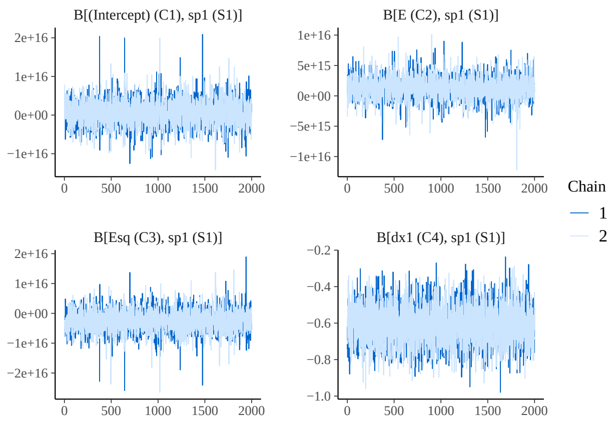


42

```
bayesplot::mcmc_areas(m11.post.hmsc$Beta, area_method = c("equal height"))
```

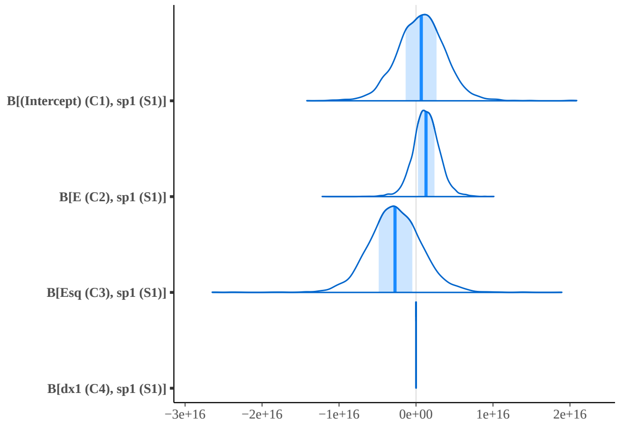


43

```
VP11 <- computeVariancePartitioning(m.11.sample, group = c(1, 1, 1, 2), groupnames = c("Env",
    "x1"))
plotVariancePartitioning(m.11.sample, VP11, cols = c("white", "skyblue", "darkgrey"),
    args.legend = list(cex = 0.75, bg = "transparent"))
```

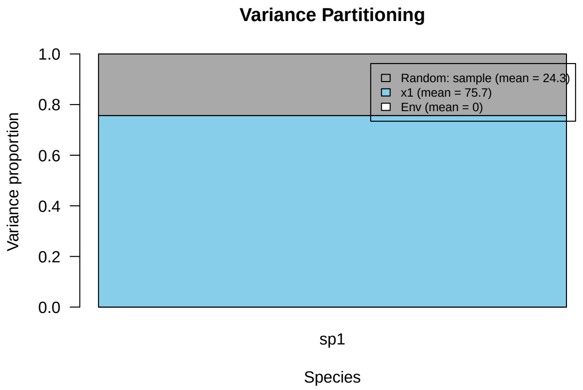


44

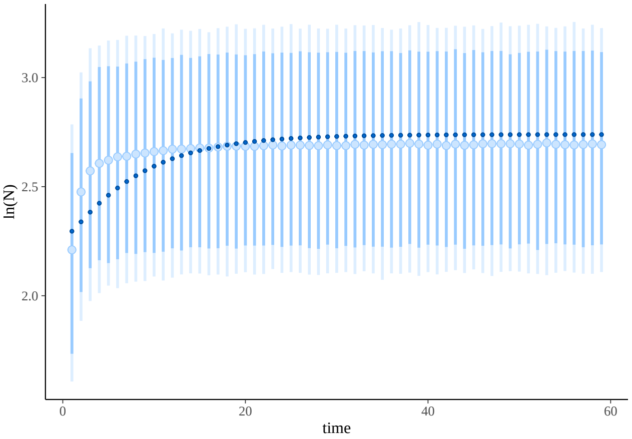


Figure 12: Simulated (dark blue points) population size over time for $h^2 = 1$, with posterior predicted mean (light blue points) and 95% highest density intervals (light blue bars).

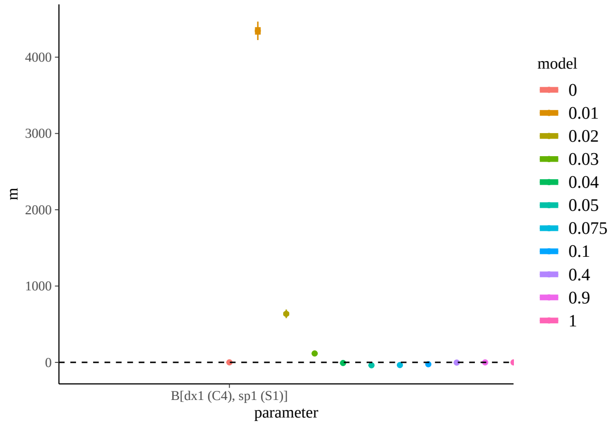


Figure 13: Estimation of linear regression coefficient for trait evolution across $h^2$ levels.

We can see that the variation in Y is a combination of the relative importance of density-dependence and of trait evolution. We can see that relative importance changes across $h^2$ levels. What drives changes in this relative importance?

```
d_dat <- gdat[gdat$time != 1, c(1, 2, 3, 5)]
d_dat <- as.data.frame(d_dat)
d_dat$dxt <- NA
d_dat$ddt <- NA
d_dat$dN <- NA
d_dat$dd <- NA
for (i in 1:11) {
    sub <- gdat[gdat$h2 == levels(gdat$h2)[i], ]
    d_dat$dxt[d_dat$h2 == levels(d_dat$h2)[i]] <- abs(sub$x1[2:60] - sub$x1[1:59])  # absolute value of
    d_dat$ddt[d_dat$h2 == levels(d_dat$h2)[i]] <- alpha[1, 1] * sub$N1[2:60] - alpha[1,
        1] * sub$N1[1:59]
    d_dat$dN[d_dat$h2 == levels(d_dat$h2)[i]] <- sub$N1[2:60] - sub$N1[1:59]
    d_dat$dd[d_dat$h2 == levels(d_dat$h2)[i]] <- alpha[1, 1] * d_dat$N1[d_dat$h2 ==
        levels(d_dat$h2)[i]]
```

```
}
```

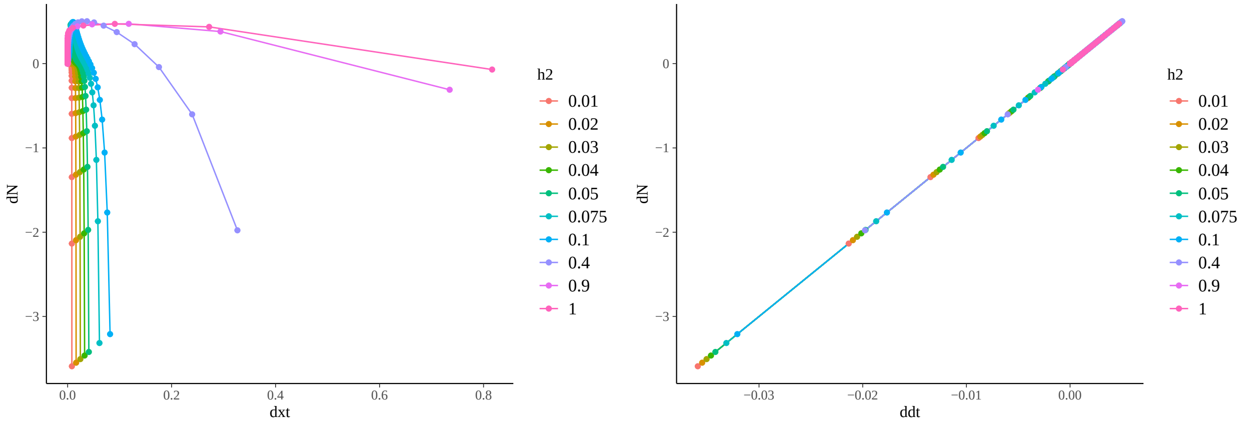


Figure 14: Density dependence varies in importance for change in population size (dN) across heritability levels. When heritability is very low, $|\Delta x|$ can be higher for more time steps, as the population is slow to adapt to the optimum trait value. This effect diminishes with increasing $h^2$. Density dependence ($dd = (\alpha_{ii} N_t) - (\alpha_{ii} N_{t-1})$) on the other hand increases with increasing heritability.
