## Appendix S4 for "Detection of eco-evolutionary dynamics in communities using Joint Species Distribution Models"

### Appendix S4. The interpretation of the estimated trait change effect size coefficient

Jelena H. Pantel* Ruben J. Hermann†

30 November, 2024

#### 1 Empirical data

To correctly interpret correlation of trait change with population density of the species (as seen in Fig. 3) we first need to check the raw data.

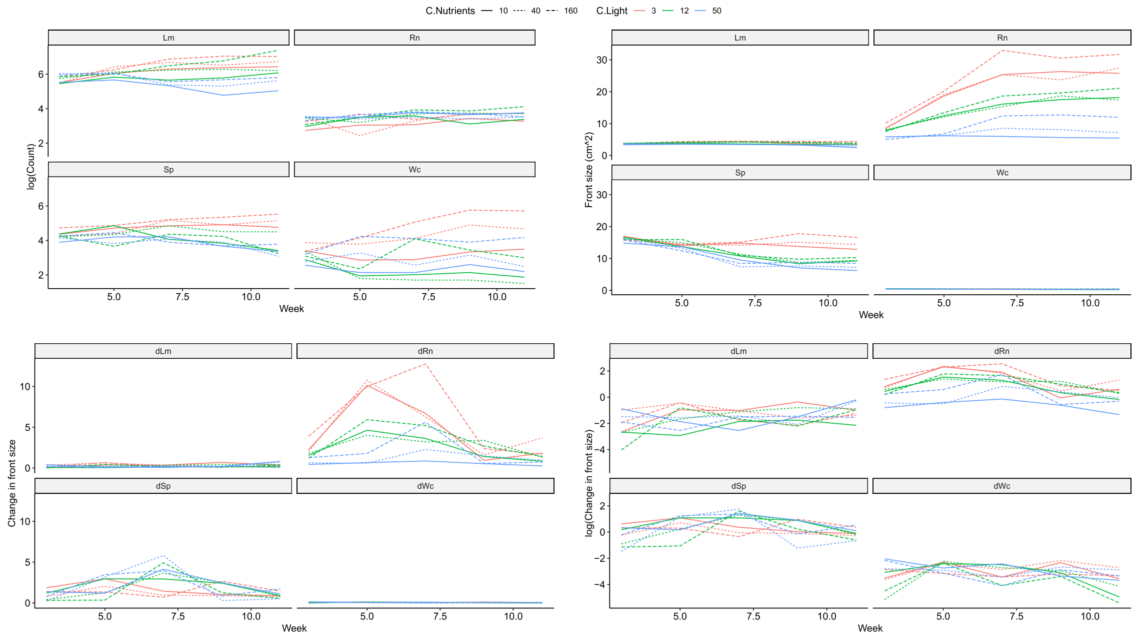


Figure 1: Figure S05 1

From the empirical data we can detect that species Rn is drastically increasing in frond size over the course of the experiment, with a peak in trait change during week 5 and 7. Simultaneously it is only slowly increasing in population density, with slightly lower density during week 5. This leads to the highest value in $|\Delta x|$ during a lower value in the population density of species Rn. Similarly species Wc had large trait changes, during time points when species Lm and Sp showed an increase in population density. HMSC draws a linear correlation between both and in (2) it can be seen how HMSC infers these correlations (the logarithmic scale of $|\Delta x|$ was used for the figure).

*Laboratoire Chrono-environnement,UMR 6249 CNRS-UFC, 16 Route de Gray, 25030 Besançon cedex, France,

†University of Duisburg-Essen, Universitätsstraße 5, 45141 Essen, Germany,

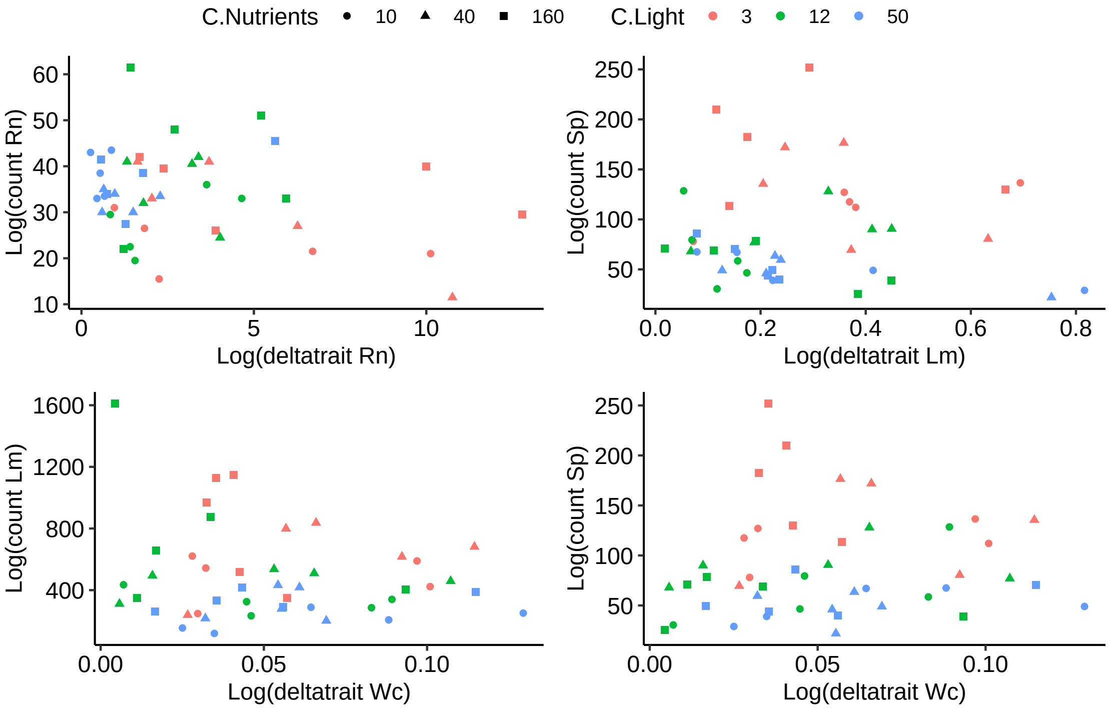


Figure 2: Figure S05 2

It should be noted that species Rn was the only to strongly increase in frond size in this experiment, which may the result from its decrease in population size. Species Wc decreased in frond size, thus occupying less space on the water surface and potentially freeing space for the other species to occupy. We speculate on this, but the manuscript for the original data will have more information.
